# In *silico* characterisation of PAL homologs and metabolomic profiling of shade response indicates potential presence of PTALs and Tyr as a probable precursor of lignin biosynthesis in conifers

**DOI:** 10.1101/2024.08.20.608774

**Authors:** Sonali Sachin Ranade, María Rosario García-Gil

## Abstract

Norway spruce and Scots pine show enhanced lignin synthesis under shade along with differential expression of defense-related genes that renders disease resilience. In general, phenylalanine (Phe) is the precursor for lignin synthesis in plants and tyrosine (Tyr) forms an additional lignin precursor in grasses. Phenylalanine ammonia-lyase (PAL) and tyrosine ammonia-lyase (TAL) from lignin biosynthesis pathway use either Phe or Tyr as precursors for lignin production, respectively. Grasses possess bifunctional phenylalanine/tyrosine ammonia-lyase (PTAL) that potentially can use both Phe and Tyr for lignin biosynthesis. Metabolomic profiles of seedlings revealed a relatively higher amount of Phe and Tyr under shade in Scots pine, while Norway spruce showed differential regulation of only Tyr under shade. Sequence analysis and phylogeny of PAL homologs in the two conifers coupled with correlation of up-regulation of precursors for lignin synthesis (Phe/Tyr) and enhanced lignin synthesis along with differential expression of PAL homologs under shade, suggests potential presence of PTALs in conifers, which is novel. Exome sequence analysis revealed latitudinal variation in allele frequencies of SNPs from coding regions of putative PAL and PTAL in Norway spruce, which may impact enzyme activity affecting lignin synthesis. Metabolomic analysis additionally identified metabolites involved in plant immunity, defense and stress response.

## Introduction

Light is one of the essential environmental factors that plays a vital role in the regulation of plant growth and development. Shade comprises of low red (R): far-red (FR) ratio and is a stressful condition for plants (Hussain et al., 2019). Norway spruce (*Picea abies* (L.) H. Karst.) and Scots pine (*Pinus sylvestris* L.), which are economically important conifer species for the Swedish forest industry have a contrasting response to shade or low R:FR ratio; Norway spruce is shade-tolerant while Scots pine is a shade-intolerant species (Ranade, Delhomme, & García-Gil, 2019). Norway spruce can grow, survive and thrive under shade as compared to Scots pine which is shade-intolerant and requires full sunlight (Grebner, Bettinger, Siry, & Kevin, 2021). Shade-tolerant species have a slow relative growth rate and a strong defense strategy as against the shade-intolerant which exhibit rapid growth and reduced defense response (Martinez-Garcia & Rodriguez-Concepcion, 2023). However, shade is perceived as stress and an unfavourable condition in both species. Scots pine displays shade avoidance syndrome (SAS) and Norway spruce exhibits shade tolerance response (STR) in response to low R:FR or shade conditions (Ranade et al., 2019).

During the growth season the northern forests in Sweden daily receive more hours of FR-enriched light/twilight or shade-like conditions (low R:FR) as compared to southern forests, due to Sweden’s geographical location. Although Norway spruce and Scots pine show contrasting responses to shade, they have adapted to latitudinal variation in twilight characterised by a northward increase in FR requirement to maintain growth (Clapham et al., 1998; Clapham, Ekberg, Eriksson, Norell, & Vince-Prue, 2002; Ranade & García-Gil, 2013; Ranade, Seipel, Gorzsás, & García-Gil, 2022b). In Norway spruce, recently we identified a latitudinal cline in SNPs that belong to the coding regions of phytochrome which are the central regulators of the light pathway (Ranade & García-Gil, 2023). These clinal variations in SNPs correlate with the latitudinal gradient in response to variable light quality and are proposed to represent signs of local adaptation to light quality in Norway spruce by this study.

Lignin is the second most abundant polymer in the secondary cell wall that renders mechanical strength, protection against pathogens and enables transport of solutes in plants (Lee et al., 2019). Lignin is synthesised in plant cells through the phenylpropanoid metabolic pathway. Phenylalanine (Phe) and tyrosine (Tyr) are the two key amino acid precursors for lignin biosynthesis, thus defining two ways of lignification in plants (Liu, Luo, & Zheng, 2018). Phenylalanine ammonia-lyase (PAL) is the first enzyme of the general phenylpropanoid pathway in most plants, where Phe is the precursor of lignin synthesis. In some bacteria and fungi, lignin biosynthesis is mediated by tyrosine ammonia-lyase (TAL) where Tyr forms the precursor of lignin synthesis. Additionally, few bacteria and fungi have the bifunctional phenylalanine/tyrosine ammonia-lyase (PTAL) that can use both Phe and Tyr for the biosynthesis of lignin (Barros & Dixon, 2020). In plants, the presence of PTAL has been reported only in grasses (monocots) (Barros et al., 2016). The enzymatic active site of PAL/PTAL contains a highly conserved Ala-Ser-Gly – MIO (4-methylidene-imidazole-5-one) electrophilic group (Peng, Engel, Aliyu, & Rudat, 2022).

Light is one of the factors that regulates the synthesis of lignin. There is a decrease in lignin synthesis under shade conditions (low R:FR) in most angiosperms, which makes the plant weak and susceptible to diseases (Hussain et al., 2019; Wu et al., 2017). However, in contrast, there is enhanced lignin synthesis under shade in the two conifers species - Norway spruce and Scots pine, irrespective of their contrasting shade tolerance responses (Ranade, Seipel, Gorzsás, & García-Gil, 2022a; Ranade et al., 2022b). In addition, a clinal variation in the regulation of defense gene expression was observed in these two species; the northern populations of both conifers in Sweden showed a higher number of defense-related genes being expressed in response to shade as compared to the southern populations. These investigations suggested that the northern populations of both conifers may be disease resilient as compared to the southern ones. Furthermore, the same studies proposed that these variations could be attributed as one of the underlying factors responsible for adapting to local environmental conditions, in this case, it being the adaptation to the light quality (twilight or FR-enriched light). Adapting to local environmental conditions renders higher mean fitness to the plants. However, comprehension of the local adaptation strategies and detection of the underlying phenomenon is challenging in forest trees. In the current study, seedlings from southern and northern populations of Norway spruce and Scots pine in Sweden were grown under continuous shade conditions in growth cabinets. Metabolomic profiling of these seedlings was carried out to reveal the metabolites involved in enhanced lignin synthesis and differential defense response under shade in the southern and northern populations of both conifer species.

Moreover, we reconfirmed the enhanced lignin synthesis in both conifer species in response to shade with pyrolysis in agreement with earlier investigations (Ranade et al., 2022a, 2022b). Sequence analysis and phylogenetic analysis of the PAL and PTAL homologs were carried out in addition to analysis of the differential expression of these genes in both conifer species in response to shade conditions. We also analysed the latitudinal variation in the SNPs from PAL/PTAL homologs in the different Norway spruce populations with the exome sequencing technology.

## Material and Methods

### Seedling growth and sampling

Seeds were collected and grown as described previously (Ranade et al., 2022a, 2022b). Briefly, seeds were collected from natural populations in Sweden from unrelated trees. Northern Norway spruce seeds were collected from Pellonhuhta (67°2′N) and the seeds for the southern population were collected between latitude 56°N and 58°N. Scots pine seeds were sampled from Kaunisvaara (67°5′N) and Lammhult (56°2′N), referred to as northern and southern populations, respectively. Seeds were germinated and grown in Percival (LED-30 Elite series) growth cabinets under continuous Shade (R:FR ratio of 0.2 and total light intensity of 36 μmol m−2 s−1) and Sun (R:FR ratio equal to 1.2 and a total light intensity of 65 μmol m−2 s−1) conditions at a temperature of 22°C on moist vermiculite (Ranade et al., 2019). These light conditions were used as they were able to trigger the shade responses in Scots pine and Norway spruce as described in our earlier work; R and FR light qualities are the two main responsible elements that plants use to determine the shade conditions and respond accordingly (Ranade et al., 2019). Seedlings were harvested at the same developmental stage when the hypocotyl is fully developed i.e., when the seed drops off and cotyledons are set free (Ranade et al., 2019). The number of days from sowing of the seeds to fully developed hypocotyl was approximately 17 ± 2 days for Norway spruce while it was 14 ± 2 days for Scots pine, under both the light treatments. Harvested seedlings were collected in liquid nitrogen and stored at −80°C until further processing. Whole seedling was used for metabolomics and pyrolysis.

### Metabolite extraction and LC-MS/GC-MS

Metabolomics including liquid chromatography–mass spectrometry (LC-MS) and gas chromatography–mass spectrometry (GC-MS) was performed on the whole seedling. Eight seedlings per population, per light treatment, from Norway spruce and Scots pine were used for metabolomics. The untargeted metabolomic approach was followed and the identification of the compounds was carried out by referring to the standards/library. The seedlings were ground into fine powder in frozen conditions using liquid nitrogen. 10-15 mg sample per seedling was used for metabolite extraction. Detailed information regarding sample preparation, mass spectrometry and data processing is included in supplementary data (Supplementary file1).

### Multivariate data analysis and pathway analysis of metabolomic data

Principal component analysis (PCA) was performed to create an overview of the data, investigate data integrity, identify potential outliers and explore possible trends and groupings of the samples (Jolliffe & Cadima, 2016). Orthogonal projections to latent structures discriminant analysis (OPLS-DA) (Trygg & Wold, 2002) were used to investigate differences in the metabolic profiles between the studied groups. A 1+0 component model (predictive + orthogonal) was used to avoid the risk of over-fitting (Trygg & Wold, 2002). The significance of a metabolite for classification in the OPLS-DA model was specified by calculating the 95% confidence interval for the loadings using jackknifing (Efron & Gong, 1983). All data was centered and scaled to unit variance. The OPLS-DA model was validated with a seven-fold cross-validation (Wold, 1978) and ANOVA of the cross validated models was used to define the model significance (Eriksson, Trygg, & Wold, 2008). All multivariate data analysis and model plots were performed in SIMCA 16.0 (Sartorius Stedim Data Analytics AB, Umeå, Sweden). Pathway analysis of the significant metabolites detected by LCMS and GCMS was performed using MetaboAnalyst 6.0 (Pang et al., 2024).

### Measurement of total lignin content

Measurement of total lignin content was performed in young seedlings (17 ± 2 days old in Norway spruce, 14 ± 2 days old in Scots pine) grown under different light treatments using Pyrolysis-Gas Chromatography/Mass Spectrometry (Py-GC/MS). Py-GC/MS was performed using two approaches, one by preparing the alcohol insoluble residue (AIR1/AIR2) and other by using the crude seedling powder. Norway spruce and Scots pine seedlings grown under the Sun and Shade conditions were dried and ball-milled into fine powder. Five seedlings per species, per population and per light treatment, were used for each pyrolysis method. Total lignin was also measured in trees ranging from 12-24 years of age from field trials by using near infrared spectroscopy (NIR).

For the AIR1/AIR2 method, firstly, ball-milled fine powder of the seedling was washed sequentially in 80% ethanol (v:v in water) for 30 min at 95°C, 70% ethanol (v:v in water) for 30 min at 95°C, 95% MeOH (v:v in water) for 10 min at room temperature, methanol:chloroform 1:1(v:v) for 10 min at room temperature and twice with acetone. The residue was dried in vacuum-desiccator overnight to obtain alcohol insoluble residue 1 (AIR1). Starch was removed from AIR1 by treating with *α*-amylase from pig pancreas (100 units per 100 mg of AIR1) in 0.1M potassium phosphate buffer, pH 7.0 containing 10mM NaCI overnight at 37 °C. The residue (AIR2) was washed with 0.1M potassium phosphate buffer, water, followed by acetone. It was dried in vacuum-desiccator overnight. 50 µg (± 10 µg) of the dried residue was applied to a pyrolyzer equipped with an auto sampler (PY-2020iD and AS-1020E, Frontier Lab, Japan) connected to a GC/MS (7890A/5975C; Agilent Technologies AB, Sweden). The pyrolysate was separated and analysed according to Gerber et al. (Gerber, Eliasson, Trygg, Moritz, & Sundberg, 2012).

To perform Py-GC/MS with the crude seedling powder, 50 µg (± 10 µg) of ball-milled fine powder of the seedling was directly applied to a pyrolyzer equipped with an auto sampler (PY-2020iD and AS-1020E, Frontier Lab, Japan) connected to a GC/MS (7890A/5975C; Agilent Technologies AB, Sweden). The pyrolysate was separated and analysed according to Gerber et al. (Gerber et al., 2012).

For NIR, 12-24 years trees from two populations maintained by Skogforsk were included – Sävar (63.89°N) and Höreda (57°N). Both populations comprised of unrelated individuals including 782 trees from Sävar (clonal archive of unrelated elite genotypes) and 1244 trees from Höreda (progeny trial of half-sib families, one tree per family has been sampled) (Morales et al., 2024). The detailed procedure for NIR has been included in the is included in supplementary data (Supplementary file2).

### Sequence analysis and phylogeny of PAL/PTAL homologs

The PAL/PTAL protein sequences of monocots and dicots were retrieved from GenBank (https://www.ncbi.nlm.nih.gov/genbank/) (Benson et al., 2013) and The Arabidopsis Information Resource (TAIR, https://www.arabidopsis.org/). The protein sequences of putative PAL/PTAL for the conifer species were retrieved from Gymno PLAZA, 1.0 (https://bioinformatics.psb.ugent.be/plaza/versions/gymno-plaza/) (Proost et al., 2015) using BLAST searches performed with the characterised PAL from *Arabidopsis thaliana* and PTAL from *Brachypodium distachyon*. Multiple sequence alignment was performed using CLUSTAL O (1.2.4) (Sievers & Higgins, 2018). The phylogenetic tree was constructed using Phylogeny.fr with the default settings (https://www.phylogeny.fr/) (Dereeper et al., 2008). In brief, alignments were created using MUSCLE (Edgar, 2004), phylogeny was done using PhyML (maximum-likelihood principle) (Guindon et al., 2010) and TreeDyn (Chevenet, Brun, Banuls, Jacq, & Christen, 2006) was used to construct the tree.

### Detection of SNP variation in PAL/PTAL genes in Norway spruce

The exome capture dataset was recruited from the previous study described in Ranade and Garcia-Gil, 2023 (Ranade & García-Gil, 2023). In short, 1654 individuals (unrelated parents from natural forests) originating from different latitudes across Sweden, were included in this study. The trees were divided into six populations following the same rationality described in Ranade and García-Gil, 2023 (Ranade & García-Gil, 2023), considering the latitude-wise variation in the amount of FR light across Sweden – the northern latitudes receive an extended period of FR light as compared with the southern ones. Trees were divided into six populations, S1-S6 as described previously (Ranade & García-Gil, 2023): S1 comprised 245 trees from latitudes 55-57, S2 – 213 trees from latitude 58, S3 – 187 trees from latitudes 59-60, S4 – 213 trees from latitudes 61-62, S5 – 573 trees from latitudes 63-64 and S6 – 223 trees from latitudes 65-67. Exome capture details have been described in Baison et al. (Baison et al., 2019). Variant calling was performed using GATK HAPLOTYPECALLER v.3.6 (Van der Auwera et al., 2013) and SNPs were annotated using SNPEFF 4 (Cingolani et al., 2012). Only bi-allelic SNPs were included in this study. The vcf file was filtered using settings; --min-alleles 2 --max-alleles 2 --maf 0.01 --remove-indels --minQ 10 --max-missing 0.9. The vcf file from the current analysis is deposited in Zenodo, which is the open-access repository developed under the European OpenAIRE program and operated by CERN (Ranade & García-Gil, 2024). SNPassoc was used to determine the allele and genotype frequencies (Gonzalez et al., 2007) and Analysis of variance and Tukey’s posthoc tests (Bonferroni *p values*) were applied to determine the statistical significance of their difference. Genetic diversity among the six different populations (pairwise FST estimates) was estimated using DnaSP 6 (Rozas et al., 2017) including both the synonymous + missense SNPs, for each of the PAL/PTAL genes. Allele frequencies in each population regarding the PAL/PTAL genes were calculated and then regressed on population latitude. R2 of the linear regression was computed as the proportion of the total variance of latitude explained by the frequency of each marker (Berry & Kreitman, 1993), where R2 is the goodness-of-fit of the linear regression model.

### PAL/PTAL gene expression in response to shade

The differential expression of putative *PAL/PTAL* genes in response to Shade was derived from the transcriptome data from our earlier study in Norway spruce (Ranade et al., 2022b) and Scots pine (Ranade et al., 2022a), where the transcriptome was analysed in seedling samples in both species from the northern and southern latitudes in Sweden, grown under Shade and Sun conditions. In brief, single-gene differential expression between the northern and southern latitudes in response to Shade was determined where Sun was the control, using DESeq2 (v1.12.0) (Love, Huber, & Anders, 2014). False discovery rate (FDR) adjusted *p values* were used to assess the significance of the expression of the genes; a common threshold of 5% was used throughout.

## Results

### Detection of metabolites in response to Shade

LCMS detected a total of 799 metabolites (targeted + untargeted) in Norway spruce and 781 metabolites (targeted + untargeted) in Scots pine (Supplementary file3, Table S1). GCMS detected and identified 69 metabolites in Norway spruce and 68 metabolites in Scots pine. Further details on the metabolites detected with LCMS and GCMS in both conifer species are included in the supplementary data (Tables S1-S5). A total of 30 identified metabolites (LCMS + GCMS) were significantly down-regulated under Shade and 21 identified metabolites were significantly up-regulated under Shade in the northern Norway spruce samples. Likewise, 41 and 25 identified metabolites (LCMS + GCMS) were significantly down-regulated and significantly up-regulated respectively, in response to Shade in the southern Norway spruce population. In the case of Scots pine, 63 identified metabolites (LCMS + GCMS) were significantly down-regulated and 29 were significantly up-regulated under Shade in the northern population, while 63 and 29 identified metabolites (LCMS + GCMS) were significantly down-regulated and significantly up-regulated respectively, in response to Shade in the southern population. Overall, a larger number of metabolites were down-regulated under Shade in both conifers.

The PCA of LCMS and GCMS data for Norway spruce and Scots pine demonstrated a clear separation between all groups – North_Shade, North_Sun, South_Shade and South_Sun (Supplementary file4, Figure S1-S3), except in case of GCMS data for Scots pine where the PCA showed clear separation between Shade and Sun conditions, but the Northern and Southern samples were not completely separated (Supplementary file4, Figure S4). The OPLS-DA (LCMS and GCMS data) showed differences between Shade and Sun conditions in both populations in both species (Supplementary file4, Figure S5-S8). Metabolite loadings for Northern ecotype plotted against Southern ecotype (OPLS-DA for South vs North models) in Norway Spruce and Scots pine showed similarity between models and highly similar metabolic response to Sun and Shade conditions for both ecotypes i.e. up/down regulation of metabolites under the Sun and Shade conditions (Supplementary file4, Figure S10-S12), except in the case of Norway Spruce LCMS data which showed only a mild similarity in the metabolic response to light condition (Supplementary file4, Figure S9). Furthermore, Scots pine exhibited a more similar metabolic response under both light conditions compared to Norway spruce (higher Q2Y in Scots pine).

Table 1 represents the list of selected statistically significant metabolites detected in response to Shade along with their fold change under Shade as compared to the Sun conditions. Table S2-S5 provided in the supplementary data gives the complete list of statistically significant metabolites that were identified by LCMS and GCMS in response to Shade in both the conifer species, at both latitudes. Up/down-regulation refers to up/down-regulation of the compound under Shade conditions in these tables.

**Table 1.** Summary of statistically significant metabolites detected in response to Shade in Norway spruce and Scots pine – Up/Down-regulation refers to Up/Down-regulation of the compound under Shade conditions.

|  | Compound | Role | Regulation under Shade: Spruce North (Fold change under Shade in parenthesis) | Regulation under Shade: Spruce South (Fold change under Shade in parenthesis) | Regulation under Shade: Pine North (Fold change under Shade in parenthesis) | Regulation under Shade: Pine South (Fold change under Shade in parenthesis) | Reference (Regarding function) |
| --- | --- | --- | --- | --- | --- | --- | --- |
| 1 | L-Leucine | Defense | Up-regulated (2.3) | Up-regulated (2.5) | Up-regulated (3.6) | Up-regulated (3.8) | (Jones & Jones, 1997) |
| 2 | gamma-Aminobutyric acid (GABA) | Defense | Up-regulated (3.2) | Up-regulated (1.6) | Up-regulated (2.1) | Up-regulated (1.2) | (Guo, Gong, Luo, Zuo, & Shen, 2023) |
| 3 | L-Serine | Plant immunity | Up-regulated (1.9) | Up-regulated (2.2) | Up-regulated (1.8) | Up-regulated (1.1) | (X. M. Zhang et al., 2023) |
| 4 | Betaine | Abiotic stress tolerance | Up-regulated (2.7) | Up-regulated (2.4) | Up-regulated (2.8) | Up-regulated (3.2) | (Ashraf & Foolad, 2007; Giri, 2011) |
| 5 | L-Threonine | Stress response | Up-regulated (1.6) | Up-regulated (1.9) | Up-regulated (1.5) | Up-regulated (1.8) | (Charlton et al., 2008) |
| 6 | Valine | Stress response, reduces vegetative growth | Up-regulated (5.4) | Up-regulated (4.1) | Up-regulated (4.6) | Up-regulated (4.9) | (Charlton et al., 2008; S. H. Li et al., 2020) |
| 7 | Threonic acid | Defense | Up-regulated (1.9) |  | Up-regulated (1.8) |  | (Wen et al., 2023) |
| 8 | L-Lysine | Defense | Up-regulated (1.9) |  | Up-regulated (1.9) | Up-regulated (1.4) | (Wen et al., 2023) |
| 9 | Ornithine | Defense | Up-regulated (1.9) |  |  | Up-regulated (1.04) | (Hussein, Mekki, Abd El-Sadek, & El Lateef, 2019) |
| 10 | L-Leucine/L-Isoleucine | Disease resistance |  | Up-regulated (3.6) | Up-regulated (3.8) |  | (Jones & Jones, 1997; Y. Li et al., 2021) |
| 11 | Tryptophan | Defense |  |  | Up-regulated (3.1) | Up-regulated (1.1) | (Zhao et al., 2022) |
| 12 | L-Proline | Biotic/abiotic stress tolerance | Up-regulated (1.6) |  |  |  | (Ashraf & Foolad, 2007) |
| 13 | Xanthone | Defense |  |  | Up-regulated (2.2) |  | (Tocci et al., 2011) |
| 14 | Caffeoyl quinic acid 4 | Defense |  |  | Down-regulated (0.4) | Down-regulated (0.2) | (Mondolot et al., 2006) |
| 15 | Coumaroyl quinic acid 2 | Defense |  |  | Down-regulated (0.6) | Down-regulated (0.4) | (Koskimäki et al., 2009) |
| 16 | Coumaroyl quinic acid 3 | Defense |  |  |  | Down-regulated (0.5) | (Koskimäki et al., 2009) |
| 17 | Kaempferol | Defense |  | Down-regulated (0.6) | Down-regulated (0.2) | Down-regulated (0.1) | (Likic, Sola, Ludwig-Müller, & Rusak, 2014) |
| 18 | Abietic acid | Defense | Down-regulated<br>(0.9) |  | Down-regulated<br>(0.5) | Down-regulated<br>(0.1) | (Trapp & Croteau, 2001) |
| 19 | Tyrosine | Lignin synthesis,<br>antioxidant, defense |  | Up-regulated<br>(5.7) | Up-regulated<br>(3.6) | Up-regulated<br>(3.1) | (Yadav & Chattopadhyay, 2023) |
| 20 | L-Phenylalanine | Lignin synthesis, defense |  |  | Up-regulated<br>(4.3) | Up-regulated<br>(1.7) | (Yadav & Chattopadhyay, 2023) |
| 21 | Shikimic acid | Lignin synthesis | Down-regulated<br>(0.6) | Down-regulated<br>(0.7) | Down-regulated<br>(0.5) | Down-regulated<br>(0.5) | (Santos-Sánchez et al., 2019) |
| 22 | L-Arginine | Nitrogen reserve | Up-regulated<br>(2.5) | Up-regulated<br>(1.5) | Up-regulated<br>(1.6) | Up-regulated<br>(1.2) | (Yang & Gao, 2007) |
| 23 | Asparagine | Nitrogen reserve | Up-regulated<br>(5.5) | Up-regulated<br>(4.1) | Up-regulated<br>(1.9) | Up-regulated<br>(1.6) | (Lea et al., 2007) |
| 24 | Oxoglutaric acid | TCA - energy-yielding<br>metabolism | Down-regulated<br>(0.6) | Down-regulated<br>(0.6) | Down-regulated<br>(0.4) | Down-regulated<br>(0.5) | (Zhang & Fernie, 2018) |
| 25 | Pyruvic acid | TCA - energy-yielding<br>metabolism | Down-regulated<br>(0.9) | Down-regulated<br>(0.6) | Down-regulated<br>(0.3) | Down-regulated<br>(0.2) | (Zhang & Fernie, 2018) |
| 26 | L-Glutamine | Metabolic fuel, defense |  |  | Down-regulated<br>(0.5) | Down-regulated<br>(0.2) | (X. M. Zhang et al., 2023) |
| 27 | Dehydroascorbic acid<br>(DHAA) | Cellular homeostasis | Down-regulated<br>(0.7) | Down-regulated<br>(0.5) | Down-regulated<br>(0.2) | Down-regulated<br>(0.2) | (Deutsch, 2000; Dreyer, 2021) |
| 28 | L-Aspartic acid | Growth and defense | Up-regulated<br>(1.2) | Up-regulated<br>(1.3) | Down-regulated<br>(0.8) | Down-regulated<br>(0.8) | (Han et al., 2021) |
| 29 | Glucose | Growth & development |  |  | Down-regulated<br>(0.6) | Down-regulated<br>(0.5) | (Yoon et al., 2021) |
| 30 | Sucrose | Growth & development | Down-regulated<br>(0.8) | Down-regulated<br>(0.7) | Down-regulated<br>(0.5) | Down-regulated<br>(0.5) | (Yoon et al., 2021) |

Generally, amino acids are the building blocks of proteins and they play a vital role in the overall growth and development throughout the plant life cycle. However, studies in plant model systems have reported a few amino acids that are also associated with defense. Amino acids involved in plant immunity (e.g. serine) (X. M. Zhang et al., 2023), defense (leucine) (Jones & Jones, 1997) and stress response (e.g. threonine, valine) (Charlton et al., 2008; Y. Li et al., 2021) were found to be up-regulated in response to Shade in both conifers at both latitudes. A common resin component involved in plant defense e.g. abietic acid (Trapp & Croteau, 2001), was found to be down-regulated under Shade at both latitudes in Scots pine and in the northern Norway spruce population, while threonic acid, engaged in plant defense (Wen et al., 2023), was detected to be up-regulated specifically in the northern populations of both the conifers. Amino acid like proline (Ashraf & Foolad, 2007) and a secondary metabolite like xanthone (Tocci et al., 2011), both involved in stress and defense response, were detected to be up-regulated, specifically in Norway spruce and Scots pine respectively (Table 1). A higher number of phenolic compounds (e.g. Caffeoyl quinic acid 4, (Mondolot et al., 2006)) related to the flavonoid biosynthesis pathway, were down-regulated in Scots pine as compared to Norway spruce populations. Tyrosine and phenylalanine involved in defense and lignin synthesis (Yadav & Chattopadhyay, 2023) were upregulated under Shade in Scots pine in both populations, while in the case of Norway spruce, only tyrosine was observed to be up-regulated under Shade in the southern population. Shikimic acid involved in the biosynthesis of lignin, aromatic amino acids (phenylalanine, tyrosine and tryptophan) and most alkaloids of plants (Santos-Sánchez, Salas-Coronado, Hernández-Carlos, & Villanueva-Cañongo, 2019), was found to be downregulated under Shade at both latitudes in both the conifers. Amino acids (e.g. arginine and asparagine) related to nitrogen reserve in plants (Yang & Gao, 2007) (Lea, Sodek, Parry, Shewry, & Halford, 2007) were found to be up-regulated in response to Shade in both conifer species and both populations, while oxoglutaric acid and pyruvic acid related to energy-yielding metabolism (Zhang & Fernie, 2018) were observed to be down-regulated in both species. Growth-related metabolites (e.g. glutamine, glucose (Yoon, Cho, Tun, Jeon, & An, 2021; X. M. Zhang et al., 2023)) were found to be down-regulated in Scots pine while these were up-regulated in Norway spruce. Figures S13-S16 from the supplementary data (Supplementary file4) represent an overview of metabolic pathways that were impacted under Shade in both conifer species. Alanine, aspartate and glutamate metabolism was the most impacted pathway in both the species under Shade.

### Total lignin content

Pyrolysis-Gas Chromatography/Mass Spectrometry (Py-GC/MS) was performed using two approaches – using AIR1/AIR2 and with the crude seedling powder. Although the Py-GC/MS by AIR1/AIR2 method showed higher lignin under Shade as compared to the Sun condition in all cases except for the southern Scots pines, it was not statistically significant. The means of the percentage (proportion) of total lignin by AIR1/AIR2 in each condition were as follows – north-Norway spruce Shade: 18.3, north-Norway spruce Sun: 18.0; south-Norway spruce Shade: 17.8, south-Norway spruce Sun: 17.6; north-Scots pine Shade: 19.1, north-Scots pine Sun: 19.0; south-Scots pine Shade: 18.2, south-Scots pine Sun: 18.4.

Py-GC/MS performed with the crude wood powder showed higher lignin under Shade as compared to the Sun condition in all cases, which was statistically significant (*p value*<0.05) in the southern populations of Norway spruce and Scots pine, but it was not statistically significant in the northern populations of both conifers. The means of the percentage (proportion) of total lignin from crude samples were – north-Norway spruce Shade: 7.5, north-Norway spruce Sun: 6.5; south-Norway spruce Shade: 8.5, south-Norway spruce Sun: 6.4; north-Scots pine Shade: 9.0, north-Scots pine Sun: 8.3; south-Scots pine Shade: 10.0, south-Scots pine Sun: 8.2.

The proportion of total lignin content derived by NIR from Sävar (north of Sweden) was detected to be higher than the proportion of total lignin in trees from Höreda (south of Sweden) (*p value*<0.05). The mean percentage (proportion) of total lignin content in trees from Höreda was 25.8, while it was 26.1 in the case of Sävar.

### Sequence analysis and phylogeny of PAL/PTAL homologs

The sequence details of all the PAL/PTAL sequences from monocots, dicots and putative PAL/PTAL sequences from conifer species included in the analysis are represented in the supplementary data (Supplementary file3, Table S6-S7). The partial alignment of PALs and PTALs from the model plants and putative PALs and PTALs from conifer species (Figure 1) suggests that the functional domains/residues are well conserved across angiosperms and conifers. The catalytically essential MIO region formed from an alanine-serine-glycine triad which is conserved in model plants was found to be present in all the conifer species except PabPAL2. Several other key amino acid residues required for the functioning of the lyase were also found to be conserved in conifers such as tyrosine as the catalytic base, arginine interacting with the carboxylic group of the substrate and, tyrosine and asparagine involved in stabilisation of the electrophilic MIO within the catalytic site (Varga et al., 2021). Residues involved in substrate specificity for Phe/Tyr in PALs/TALs/PTALs according to the previous studies were detected in the putative PALs and PTALs from conifer species (F/H, A/S, L/V, I/L, D/E) (Barros et al., 2016; Feduraev et al., 2020; Hsieh, Ma, Yang, & Lee, 2010; Jun et al., 2018; Xue, McCluskey, Cantera, Sariaslani, & Huang, 2007). The F/H residue is crucial for substrate specificity; F is conserved between all the PALs in angiosperms while H residue is conserved in the PTALs (Barros et al., 2016). Although none of the conifers possessed the H residue at the position, the other residues involved in substrate specificity for Phe/Tyr seem to be conserved in conifers with exceptions (Figure 1). Similar to the bifunctional PTALs from grasses, PtaPTAL2, PabPTAL3 and PmePTAL3 have a conserved E residue rather than D residue present in PALs (refer to D/E position in Figure 1). Likewise, PtaPTAL3 and PmePTAL4 have S and E (refer A/S and D/E position in Figure 1), while PsiPTAL, PmePTAL1, PmePTAL2, PsyPTAL, PtaPTAL1, PabPTAL1, PabPTAL2 have S and V (refer A/S and L/V position in Figure 1), similar to PTALs from grasses. The alignment of the complete sequences of PAL/PTALs (Supplementary file4, Figure S17) showed high sequence similarity among the angiosperms and conifers especially in the regions containing the functional domains/residues including the conserved motif “GTIT<u>ASG</u>DLVPLSYIA” with the MIO region (<u>ASD</u>) (He et al., 2020). The putative PALs and PTALs from conifer species do not tend to blend and instead appear to be well separated into distinct clades; moreover, they also form distinct clades separated from the dicots and monocots (Figure 2).

**Figure 1:**
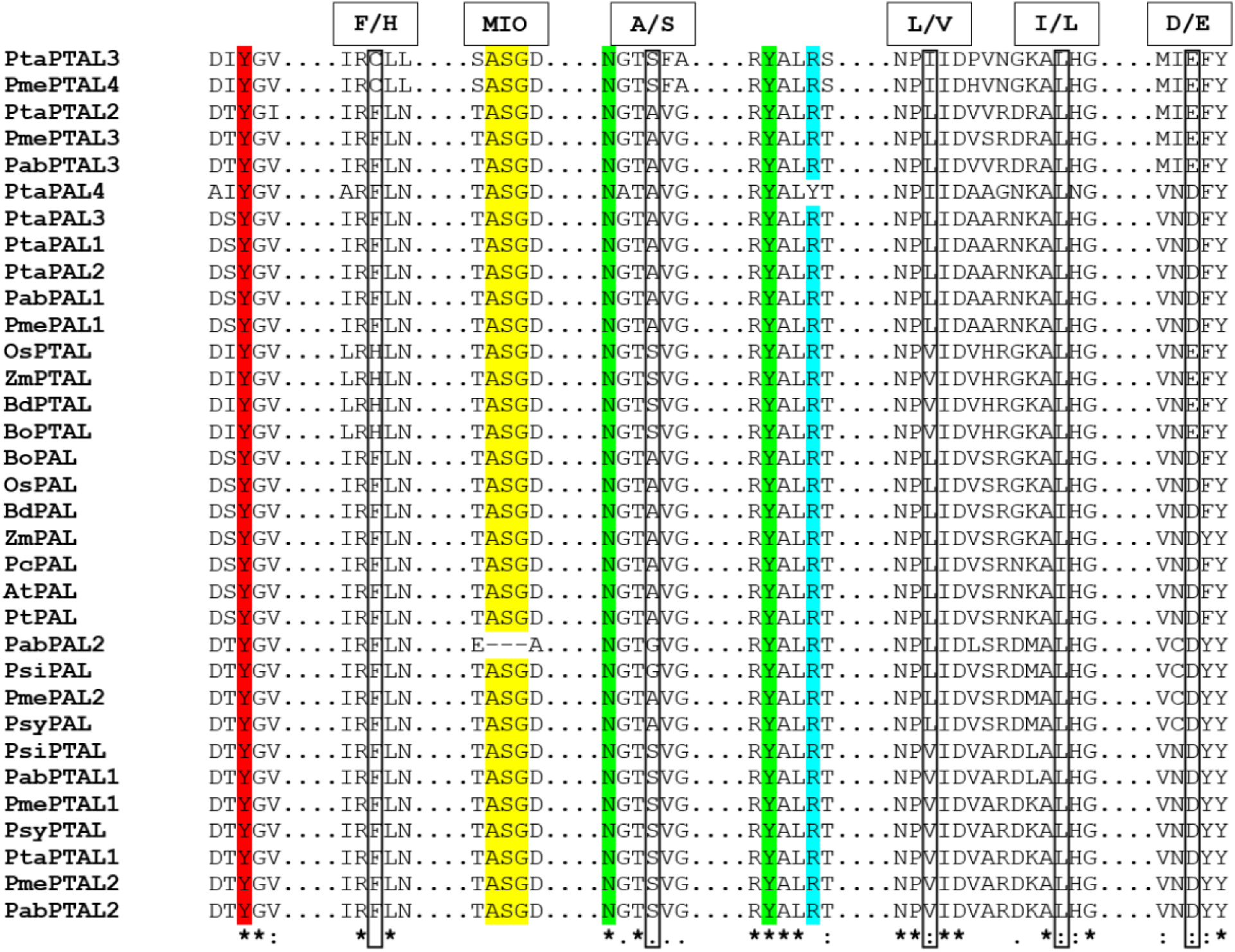
Partial alignment of PALs and PTALs from plant model systems and putative PALs and PTALs from conifer species showing conserved amino acid residues. Colour codes – red: catalytically essential tyrosine; yellow: MIO region; green: asparagine and tyrosine stabilizing the MIO group; blue: arginine responsible for binding the carboxylic group of the substrate. Residues involved in substrate specificity for phenylalanine/tyrosine are marked with boxes (F/H, S/A, V/L, L/I, E/D). At: *Arabidopsis thaliana*; Bd: *Brachypodium distachyon*; Bo: *Bambusa oldhamii*; Pc: *Petroselinum crispum*; Pt: *Populus trichocarpa*; Os: *Oryza sativa*; Zm: *Zea mays*; Pab: *Picea abies*; Pme: *Pseudotsuga menziesii*; Psi: *Picea sitchensis*; Psy: *Pinus sylvestris*; Pta: *Pinus taeda*.

**Figure 2:**
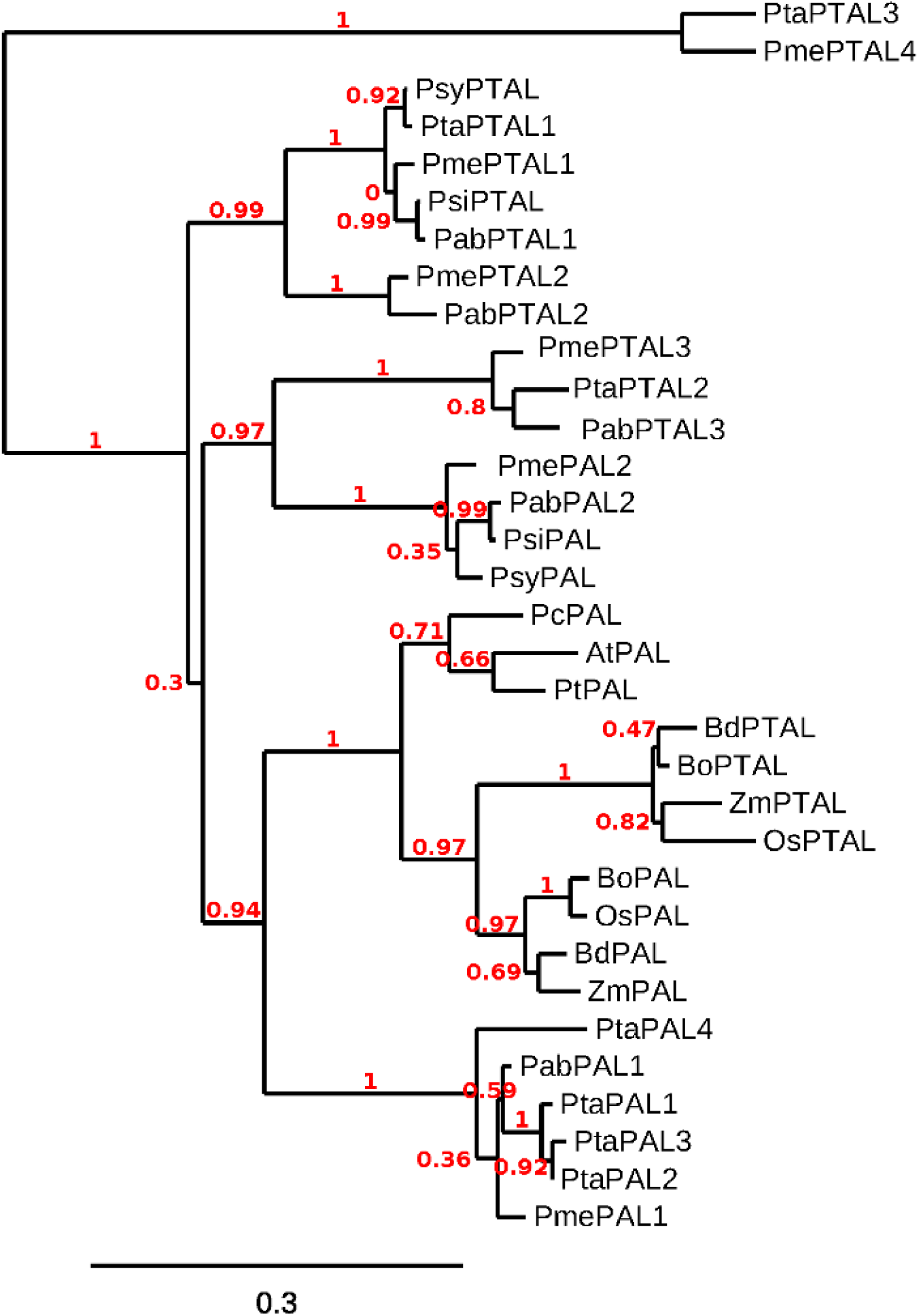
Phylogenetic tree of PTALs and PALs in plant model systems and putative PTALs and PALs in conifer species. At: *Arabidopsis thaliana*; Bd: *Brachypodium distachyon*; Bo: *Bambusa oldhamii*; Pc: *Petroselinum crispum*; Pt: *Populus trichocarpa*; Os: *Oryza sativa*; Zm: *Zea mays*; Pab: *Picea abies*; Pme: *Pseudotsuga menziesii*; Psi: *Picea sitchensis*; Psy: *Pinus sylvestris*; Pta: *Pinus taeda*.

### Clinal variation in SNPs detected in PAL/PTAL in Norway spruce

Out of the two PALs and three PTALs from Norway spruce, SNPs were detected in the coding regions of one PAL (PabPAL2; 14 missense and 20 synonymous) and two PTALs (PabPTAL1; 3 missense and 5 synonymous and PabPTAL3: 4 missense and 2 synonymous) (Supplementary file3, Table S8). Statistically significant clinal variation in the allele and genotype frequencies was detected in two synonymous and one missense mutation in PabPAL2, while PabPTAL1 and PabPTAL3 showed significant latitudinal variation in two missense mutations, respectively (Table 2). One-way ANOVA of the allele frequencies and genotype frequencies of SNPs detected in PAL/PTAL showing cline is represented in the supplementary data (Supplementary file3, Table S9-S10).

**Table 2.** PAL/PTAL SNPs in Norway spruce showing clinal variation and the population-wise allele frequencies of the reference and alternate alleles.

| Gene | Mutation | Variation | Allele | Population-wise allele frequency |  |  |  |  |  |
| --- | --- | --- | --- | --- | --- | --- | --- | --- | --- |
|  |  |  |  | S1 | S2 | S3 | S4 | S5 | S6 |
| PabPAL2 | Synonymous | Ala228Ala: Reference T, alternate C; GCT → GCC | Reference (T) | 0.50 | 0.50 | 0.50 | 0.49 | 0.48 | 0.47 |
|  |  | (Nonpolar, hydrophobic → Nonpolar, hydrophobic) | Alternate (C) | 0.50 | 0.50 | 0.50 | 0.51 | 0.52 | 0.53 |
|  | Synonymous | Pro356Pro: Reference G, alternate C; CCG → CCC | Reference (G) | 0.97 | 0.96 | 0.96 | 0.95 | 0.92 | 0.86 |
|  |  | (Nonpolar, hydrophobic → Nonpolar, hydrophobic) | Alternate (C) | 0.03 | 0.04 | 0.04 | 0.05 | 0.08 | 0.14 |
|  | Missense | Ala383Thr: Reference G, alternate A; GCA → ACA | Reference (G) | 0.90 | 0.90 | 0.87 | 0.86 | 0.85 | 0.77 |
|  |  | (Nonpolar, hydrophobic → Polar, uncharged) | Alternate (A) | 0.10 | 0.10 | 0.13 | 0.14 | 0.15 | 0.23 |
| PabPTAL1 | Missense | Ala535Val: Reference C, alternate T; GCC → GTC | Reference (C) | 0.90 | 0.90 | 0.87 | 0.88 | 0.84 | 0.80 |
|  |  | (Nonpolar, hydrophobic → Nonpolar, hydrophobic) | Alternate (T) | 0.10 | 0.10 | 0.13 | 0.12 | 0.16 | 0.20 |
|  | Missense | Ala537Val: Reference C, alternate T; GCT → GTC | Reference (C) | 0.97 | 0.97 | 0.95 | 0.93 | 0.89 | 0.86 |
|  |  | (Nonpolar, hydrophobic → Nonpolar, hydrophobic) | Alternate (T) | 0.03 | 0.03 | 0.05 | 0.07 | 0.11 | 0.14 |
| PabPTAL3 | Missense | Ser595Phe: Reference C, alternate T; TCC → TTC | Reference (C) | 0.94 | 0.95 | 0.91 | 0.87 | 0.89 | 0.83 |
|  |  | (Nonpolar, uncharged → Nonpolar, hydrophobic) | Alternate (T) | 0.06 | 0.05 | 0.09 | 0.13 | 0.11 | 0.17 |
|  | Missense | Leu601Phe: Reference A, alternate T; TTA → TTT | Reference (A) | 0.95 | 0.95 | 0.92 | 0.89 | 0.89 | 0.85 |
|  |  | (Nonpolar, hydrophobic → Nonpolar, hydrophobic) | Alternate (T) | 0.05 | 0.05 | 0.08 | 0.11 | 0.11 | 0.15 |

Ala537Val from PabPTAL1 displayed the steepest cline among all (Figure 3), meaning the difference between the allele frequencies between the extreme southern and northern populations was the highest. Other SNPs from PabPAL2, PabPTAL1 and PabPTAL3 that showed latitudinal variation in the allele and genotype frequencies are represented in Figures S18-S23 included in the supplementary data. The variations that did not show any latitudinal variation in their allele/genotype frequencies could be referred to as controls (Supplementary file3, Table S8). These include several SNPs from PabPAL2, PabPTAL1 and PabPTAL3. No SNPs were detected in PabPAL1 and PabPTAL2.

**Figure 3:**
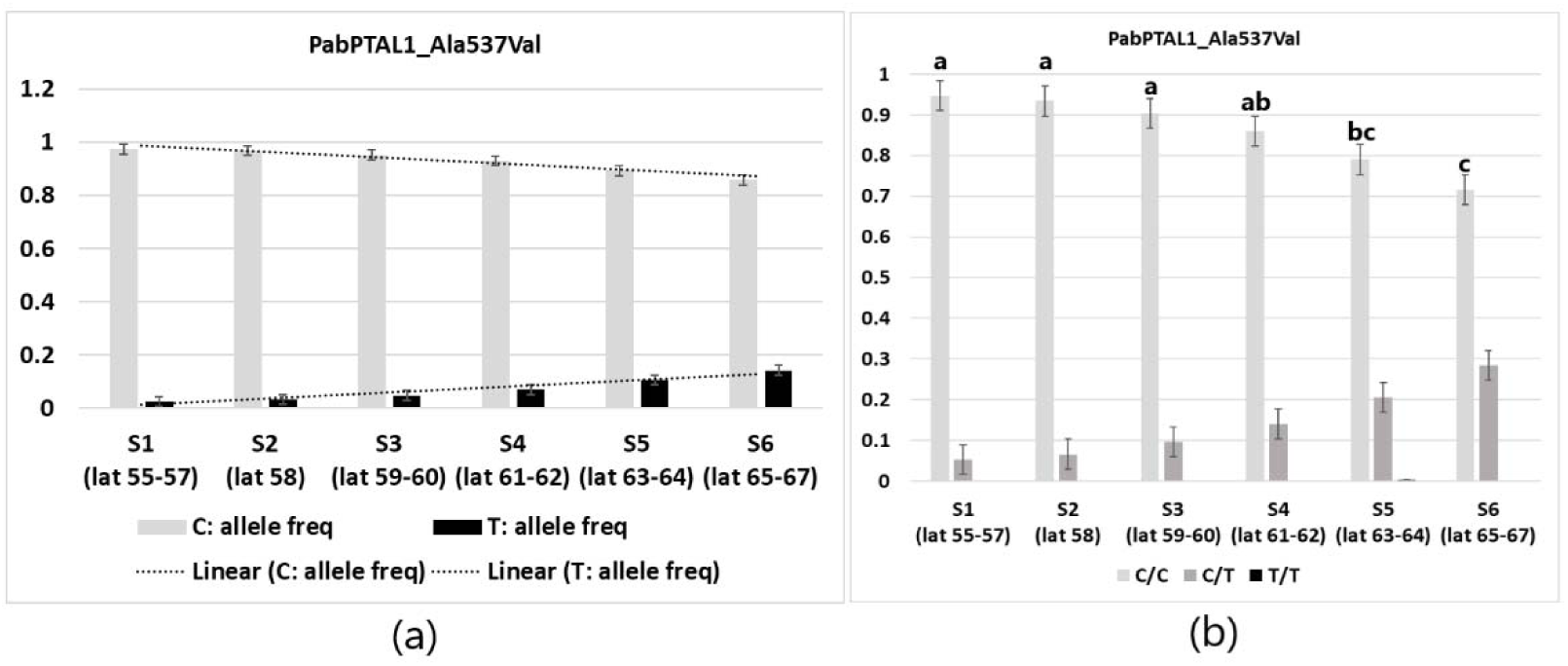
Latitudinal variation in the SNP (Ala537Val) from PabPTAL1 gene in Norway spruce populations across Sweden, which was the steepest among all the SNPs detected in the PAL/PTAL genes. (a) Cline in the allele frequencies of Ala537Val. (b) Cline in the genotype frequencies of Ala537Val. One-way ANOVA and Tukey’s posthoc test was performed with the genotype frequencies. Tukey’s posthoc categorisation is indicated above the bars.

PabPTAL3 showed the highest *F_ST_* values along with higher and less dispersed R^2^ values as compared to PabPAL2 and PabPTAL1 which suggests that it may exhibit more precise clinal variation among the three genes (Figure 4), considering all the SNPs taken together. However, overall, in accordance with the literature available on Norway spruce summarised previously (Ranade & García-Gil, 2021, 2023), the pair-wise *F_ST_* estimates for six populations of Norway spruce across Sweden (Table 3) were low, indicating low population genetic differentiation.

**Figure 4:**
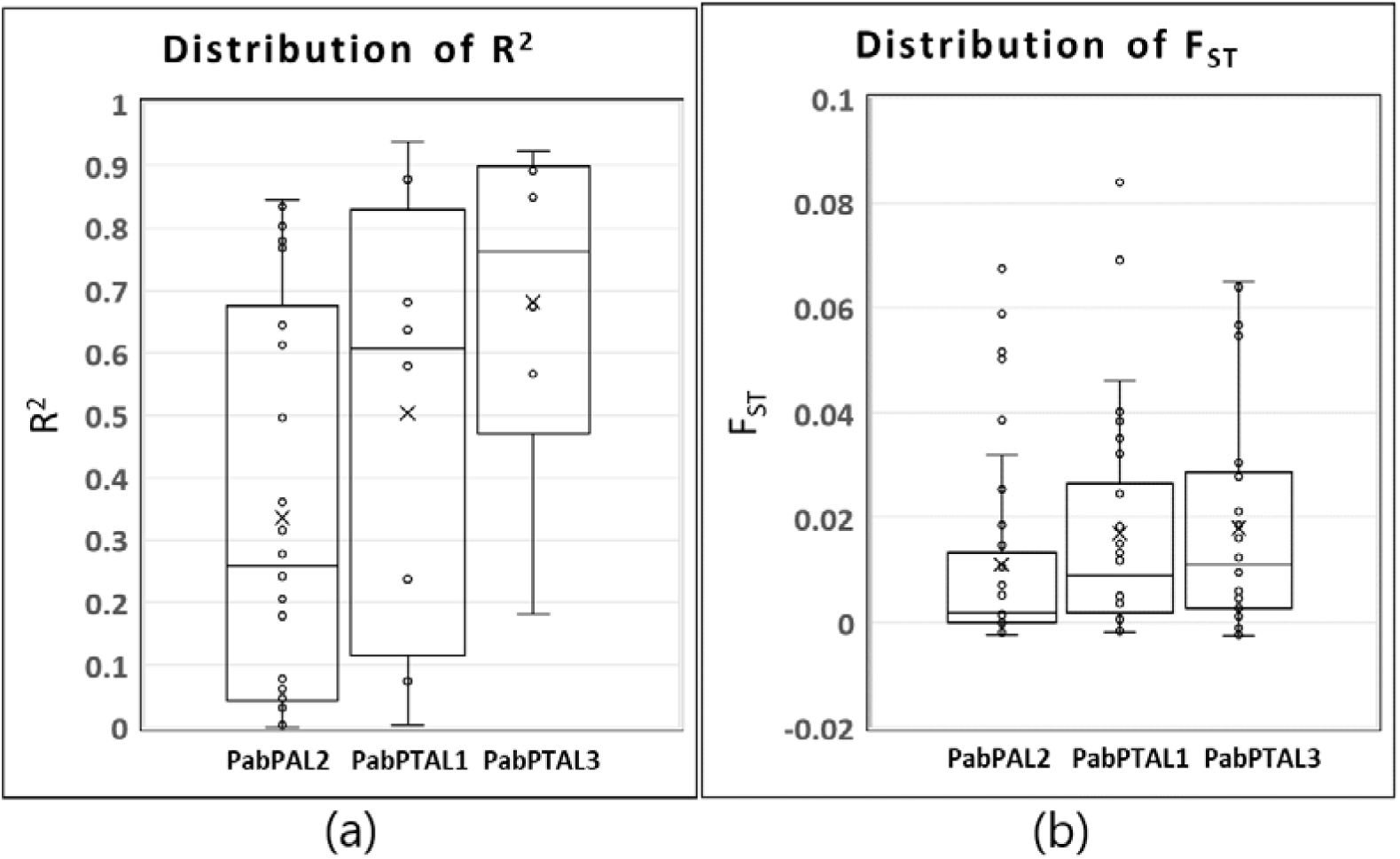
R2 and FST were calculated with the allele frequencies of all the missense and synonymous SNPs detected in the PAL/PTAL genes of the Norway spruce populations in Sweden. (a) The distribution of R2. (b) The distribution of F_ST_.

**Table 3.** Pairwise F_ST_ estimates for six populations of Norway spruce across Sweden involved in the detection of clinal variation in the SNPs in PAL/PTAL genes.

| Population | S2 | S3 | S4 | S5 | S6 |
| --- | --- | --- | --- | --- | --- |
| S1 | 0 | 0.00198 | 0.00349 | 0.00143 | 0.01225 |
| S2 |  | 0 | 0 | 0 | 0.00604 |
| S3 |  |  | 0 | 0 | 0.00797 |
| S4 |  |  |  | 0 | 0.01026 |
| S5 |  |  |  |  | 0.00375 |

## Discussion

### Presence of potential PTAL gene family in conifers that catalyses Tyr assimilation into the lignin biosynthetic pathway

Lignin is synthesised from Phe/Tyr via the phenylpropanoid metabolic pathway in plant cells (Liu et al., 2018). In most plants (dicots), lignin is synthesised from Phe, where the first step involves deamination of Phe by PAL producing cinnamic acid. Cinnamate 4-hydroxylase (C4H) produces p-coumaric acid from cinnamic acid by introducing a hydroxyl group into the phenyl ring of cinnamic acid and p-coumaric acid forms the precursor for all monolignols. Monolignol biosynthesis in grasses is reported with fewer steps; p-coumaric acid is produced directly from Tyr through the PTAL enzyme (Barros et al., 2016). Thus, in dicots, lignin is synthesised only from Phe while both Phe and Tyr form the basis for lignin synthesis in monocots. The biosynthesis of lignin in plants requires the provision of its essential precursors, Phe or Tyr. The supply of exogenous Phe or Tyr leads to a lignin deposition which indicates that the availability of Phe or Tyr is a deciding factor for lignin synthesis (Feduraev et al., 2020; P. Wang et al., 2018).

In the current work, the metabolomic analysis detected up-regulation of Phe and Tyr together with higher lignin synthesis in Scots pine, whereas in Norway spruce, up-regulation of only Tyr was detected along with higher lignin synthesis. In grasses, although Tyr is preferentially incorporated into the S-units of lignin, Phe is also utilised in the formation of the S-units (Barros et al., 2016). In this regard, it is worth noting that conifers mainly contain G-units with little or no S-units of lignin (Ralph, Lapierre, & Boerjan, 2019), therefore they may follow alternative mechanisms to efficiently utilise Tyr for lignin formation, which needs further validation. Lastly, although there is enough evidence for an increase in lignin synthesis under Shade, the concurrent down-regulation of shikimic acid that is one of the key elements involved in the biosynthesis of lignin, also warrants further investigation in conifers.

Multiple PAL/PTAL genes are found in monocots, e.g. in grasses that represent distinct PAL/PTAL monocot-specific clades in phylogenetic tree (Barros et al., 2016; Schaker et al., 2017). Likewise, many PAL genes have been described in dicots (e.g. *Arabidopsis* and tomato) which form a dicot-specific PAL clade in the phylogenetic tree (Schaker et al., 2017; F. L. Zhang et al., 2023). Previous research has documented a phylogenetically diverse set of PAL enzymes in gymnosperms (Bagal, Leebens-Mack, Lorenz, & Dean, 2012) and suggested the existence of a gymnosperm-specific lineage of PAL genes (Bagal et al., 2012; Neale et al., 2017). Our findings on the PAL homologs in Scots pine and Norway spruce align with these studies; the phylogenetic tree with the putative PAL/PTAL sequences detected in the conifers showed distinct clades comprising either PAL or PTAL, also the respective clades are distinct from the dicots and monocots. Additionally, our research contributes to a novel discovery of potential PTAL gene family in conifers, which was previously thought to be absent (Barros et al., 2016).

Finally, the novel detection of potential PTALs in both conifer species with supporting evidence from expression data, sequence analysis and phylogeny coupled with the up-regulation of Tyr and enhanced lignin synthesis under Shade, strongly supports the concept of Tyr as a probable precursor of lignin biosynthesis in conifer species which is a new finding reported by this investigation. Further research is required to validate the functionality of PTALs and lignin synthesis using Tyr in conifers.

### Possible role of PAL and PTAL gene families in local adaptation through lignin content regulation

Our previous investigation regarding transcriptomic data in Scots pine and Norway spruce in response to Shade (Ranade et al., 2022a, 2022b), indicates that the PTAL gene family is expressed in conifers (Supplementary file3, Table S6-S7). In Norway spruce, *PabPAL2* was found to be up regulated under Shade together with enhanced lignin synthesis in the northern population but not in the southern. Furthermore, our current results revealed a latitudinal cline in the missense variations from the coding regions of PAL and PTALs in Norway spruce populations across Sweden. Ala537Val from PabPTAL1 may lead to minor or no alterations in the chemical properties of amino acids, as both Ala and Val are nonpolar/hydrophobic. However, Ala383Thr from PabPAL2 and Ser595Phe from PabPTAL3 represent the change in the properties of amino acids, as the former represents a conversion from nonpolar/hydrophobic to polar/uncharged while the latter is the conversion from nonpolar/uncharged to nonpolar/hydrophobic. Variations in SNPs in the coding regions of the putative PAL/PTAL genes in Norway spruce, especially when they result in amino acid substitutions with different chemical properties, may alter protein function. This can affect enzyme activity and may lead to differential lignin synthesis, independent of gene expression regulation. Such clinal variation suggests the potential role of the PAL/PTAL gene family in local adaptation through lignin synthesis regulation, meriting further investigation.

Another aspect that supplements the higher lignin synthesis in northern Norway spruce populations in response to Shade is differential expression of the MYB3 transcription factor which is involved in the repression of lignin synthesis. Lower expression of two copies of the MYB3 transcription factors was detected in the northern Norway spruce populations as compared to the southern ones in response to Shade (Ranade et al., 2022b). Interestingly, one of these MYB3 copies also showed a steep latitudinal cline in allelic and genotypic frequencies of a missense SNP in its coding region (Ranade & García-Gil, 2021).

Regarding the lignin pathway, monolignols are synthesised in the cytoplasm and are then translocated to the cell wall where they undergo polymerisation leading to lignin formation and deposition (Alejandro et al., 2012). AIR1 removes unpolymerised lignin (monolignols) from the samples (cytoplasm and cell wall) and thus AIR1/AIR2 gives the measurement of only the polymerised lignin present in the cell wall. Py-GC-MS of the crude sample measures unpolymerised and polymerised lignin i.e. monolignols present in cytoplasm and cell wall, and the polymerised lignin from the cell wall.

Enhanced lignin synthesis under Shade reported in the current study aligns with our previous studies in Norway spruce and Scots pine (Ranade et al., 2022a, 2022b). Although the higher lignin detected under Shade in most cases was not statistically significant (e.g. AIR1/AIR2), it is important to highlight that there is a trend for enhanced lignin synthesis under Shade conditions in both conifer species in both northern and southern ecotypes. Yet, above all, this is contrasting to most angiosperms where there is a significant decrease in lignin synthesis in response to shade (Hussain et al., 2019; Wu et al., 2017). The probable reason for lignin content detected by AIR1/AIR2 not being statistically significant as compared to the Py-GC-MS with the crude sample may be because the monomers are yet being synthesised and are being transported to the cell wall but have not yet undergone polymerisation to form lignin. Therefore, the measurement of lignin monomers + polymerised lignin (Py-GC-MS with the crude sample) was statistically significant as compared to the lignin content measured with Py-GC-MS performed by AIR1/AIR2 method which measures only the polymerised lignin. Lignin formation in the cell wall is an irreversible process and it is this irreversibility that governs the necessity for the strict regulation of the lignification process (Y. Wang, Chantreau, Sibout, & Hawkins, 2013). Therefore, another reason may be attributed to seedlings representing a very early stage of development in the process of lignification in tree species (Ruzicka, Ursache, Hejátko, & Helariutta, 2015), probably not a very likely stage to see the striking differences in lignin content. Regarding the north versus south comparisons in both conifers, no significant difference in lignin content was detected and this could also be attributed to the early developmental stage of the seedlings. Yet, the older Norway spruce trees from the north (Sävar) showed a significantly higher lignin content (NIR) as compared to the trees from the southern population (Höreda). Likewise, in the case of Scots pine, the fold change in the increase of Phe and Tyr under Shade is higher in the northern populations as compared to the southern (Table1). These findings support the hypothesis that higher lignin content in northern conifer species is due to the increased exposure to FR-rich light during the growth season, suggesting a combination of plastic and adaptive genetic responses (Ranade & García-Gil, 2021; Ranade et al., 2022a, 2022b).

### Detection of metabolites related to plant defense

In addition to lignin biosynthesis, Phe and Tyr also serve as precursors for several metabolites having diverse physiological functions as antioxidants and defense compounds (Pascual et al., 2016; Schenck & Maeda, 2018). Threonic acid, which is specifically involved in defense (Wen et al., 2023), was found to be up-regulated under Shade in the northern populations of both conifers and amino acids generally involved in defense, e.g. L-leucine (Jones & Jones, 1997) were found to be up-regulated in response to Shade in both conifers at both latitudes. Recently, it was demonstrated that threonic acid along with lysine was critical for recruiting beneficial microorganisms that protected the plants from pathogen attacks in cucumbers (Wen et al., 2023). The key role played by the leucine-rich repeat proteins in plant defense has been well documented (Jones & Jones, 1997). Threonic acid is a weak sugar acid derived from threose while L-threonic acid is produced by the degradation of ascorbic acid under oxidative conditions at alkaline pH (Loewus, 1999). Ascorbic acid helps plants to defend against oxidative stress as it is the most abundant water-soluble antioxidant (Shen et al., 2021). Ascorbic acid is reversibly oxidised into dehydroascorbic acid (DHAA) upon exposure to light, heat, transition metal ions and low alkaline pH (Thurnham, 2000; Yin et al., 2022); the stability of DHAA lasts only for a few minutes and it is further irreversibly hydrolyses to form 2,3-diketogulonic acid (Zilva, 1928). The reduction of DHAA leads to the formation of ascorbic acid which takes part in regulating cellular homeostasis (Deutsch, 2000; Dreyer, 2021); cellular homeostasis is crucial for the establishment of balanced conditions for the controlled commencement and performance of various biochemical processes. DHAA was detected to be down-regulated in both species at both latitudes in response to Shade. However, gamma-aminobutyric acid (GABA) that not only helps the plant to respond to biotic and abiotic stresses but is also involved in maintaining cellular homeostasis (Guo et al., 2023), was up-regulated under Shade in both species at both latitudes.

Flavonoids play a key role in plant defense by protecting plants from different biotic and abiotic stresses (Divekar et al., 2022; Panche, Diwan, & Chandra, 2016). The phenolic compounds (e.g. caffeoyl quinic acid 4, coumaroyl quinic acid 2 and coumaroyl quinic acid 3; Table 1, Tables S2-S5 from Supplementary file3) related to flavonoid biosynthesis were down-regulated in Scots pine, while in Norway spruce these compounds were not statistically significant. This could be extrapolated as the SAS response in Scots pine associated with reduced defense response as compared to the STR response in Norway spruce where no significant difference in the number of defense-related genes was reported under the Sun and Shade conditions (Ranade et al., 2019). In this context, the regulation of defense-related metabolites is more pronounced in Norway spruce in general, meaning that overall, the fold change of the up-regulated defense-related metabolites is higher and the fold change of down-regulated defense-related metabolites is lower in Norway spruce as compared to Scots pine.

### Detection of metabolites related to plant growth and development

The metabolic pathways commonly involved in the growth and development of plants such as glycolysis/gluconeogenesis, starch and sucrose metabolism, pentose phosphate pathway, carbon fixation in photosynthetic organisms and TCA cycle were altered in response to shade in both the conifer species (Figures S13-S16 from the Supplementary file4). As Norway spruce is shade-tolerant, it can grow, survive and thrive under shade as compared to Scots pine which is shade-intolerant. Growth-related metabolite like aspartic acid was found to be up-regulated under shade in Norway spruce, while it was down-regulated in Scots pine. Like-wise, Shade did not significantly affect glucose regulation in Norway spruce, but glucose was found to be down-regulated in Scots pine. Glucose is synthesised during photosynthesis using carbon dioxide and water using light; glucose is the key carbon source acting as the signalling molecule regulating various metabolomic processes (Siddiqui, Sami, & Hayat, 2020). Glucose affects plant growth, improves the harmful effects of abiotic stress by increasing the level of antioxidants and induces the synthesis of chlorophyll, thereby regulating photosynthesis (Siddiqui et al., 2020). While sucrose is down-regulated under Shade in both conifers, the fold change is lower in Scots pine. Sucrose is synthesised from monosaccharides (e.g. fructose and glucose) in photosynthetically active cells. As sucrose is a disaccharide, its usage is more energy efficient for transport and storage as compared to fructose and glucose (Geiger, 2020). In addition, as sucrose is a non-reducing sugar, therefore it cannot be oxidised and intermediate reactions with other molecules do not take place (Geiger, 2020).

Oxoglutaric acid and pyruvic acid which are considered as the energy-yielding metabolites (Zhang & Fernie, 2018) were detected to be down-regulated under Shade in both conifers, following the similar trend as observed in sugars. However, the fold change of their down-regulation was lower in Scots pine as compared to Norway spruce which may be attributed to its shade-tolerant nature. Glutamine, which is known to function as metabolic fuel, apart from taking part in defense (X. M. Zhang et al., 2023) was found to be down-regulated in Scots pine, whereas Shade did not alter its regulation in Norway spruce. This again can be due to the shade-tolerant characteristic feature exhibited by Norway spruce.

## Conclusion

Overall, the results from the earlier investigations (Ranade et al., 2022a, 2022b) together with the current analysis further strongly support enhanced lignin synthesis under shade in conifers. The current analysis reports new findings regarding the lignin pathway in conifers. Based on the sequence analysis and phylogeny of potential PAL/PTAL homologs in conifers coupled with the correlation of up-regulation of the precursors of lignin (Phe/Tyr), differential expression of PAL/PTAL homologs and enhanced lignin synthesis under shade conditions, we propose the prospective of the presence of PTALs and biosynthesis of lignin using Tyr in conifers. The results suggest that the PAL/PTAL gene families thus play a role in the local adaptation/enhanced defence through modulating the lignin synthesis. Yet, additional research is needed to validate the functionality of PTALs and to reveal the underlying molecular mechanism involved in lignin biosynthesis using Tyr as a precursor in conifers.

## Acknowledgments

This work was supported by grants from the SLU Skogsskadecentrum, the Knut and Alice Wallenberg Foundation (KAW 2016.0352 and KAW 2020.0240), the Swedish Governmental Agency for Innovation Systems (VINNOVA, 2016-00504), FORMAS, the Swedish Foundation for Strategic Research (SSF) and the Swedish Research Council (VR). Swedish Metabolomics Centre, Umeå, Sweden (www.swedishmetabolomicscentre.se) is acknowledged for metabolic profiling by GC-MS and LC-MS. Data analysis support for metabolomics data was provided by the Computational Analytics Support Platform (CASP), Umeå University, Sweden. We thank the Biopolymer Analytical Platform (BAP) hosted by SLU and KBC of Umeå University and supported by Bio4Energy for the Pyrolysis-GC/MS analysis. We thank Skogforsk for providing access to the field trials for lignin measurement in trees. We thank RISE Research Institutes of Sweden for the analysis of lignin content in trees. We are grateful to Dr. Junko Takahashi-Schmidt, BAP and Prof. Ewa Mellerowicz, UPSC for helpful suggestions and discussions regarding pyrolysis.

## Author contributions

SSR contributed to experimental design, experiment performance, data collection, data analysis and interpretation, and manuscript writing. MRGG contributed with experimental design, data analysis and interpretation, and manuscript writing. Both the authors read and approved the manuscript.

## Conflict of interests

The authors declare no conflict of interest.

## Data availability

The vcf file from the current analysis containing data from the exome sequencing results is deposited in Zenodo (https://doi.org/10.5281/zenodo.12605324). All other data are included in the supplementary data.

## Notes

https://doi.org/10.5281/zenodo.12605324

### Competing Interest Statement

The authors have declared no competing interest.

