## Supplementary file1 for "In *silico* characterisation of PAL homologs and metabolomic profiling of shade response indicates potential presence of PTALs and Tyr as a probable precursor of lignin biosynthesis in conifers"

**Metabolic profiling by GC-MS and LC-MS**

***Sample preparation:*** Sample preparation was performed according to Gullberg et al (Gullberg et al. 2004) using 10-15 mg sample per seedling. In detail, 1000 µL of extraction buffer (20/20/60 v/v chloroform:water:methanol) including internal standards for both GC-MS and LC-MS. LC-MS internal standards were as follows: 13C9-Phenylalanine, 13C3-Caffeine, D4-Cholic acid, 13C9-Caffeic Acid were obtained from Sigma (St. Louis, MO, USA). GC-MS internal standards were as follows: L-proline-13C5, alpha-ketoglutarate-13C4, myristic acid-13C3, cholesterol-D7 were obtained from Cil (Andover, MA, USA); Succinic acid-D4, salicylic acid-D6, L-glutamic acid-13C5,15N, putrescine-D4, hexadecanoic acid-13C4, D-glucose-13C6, D-sucrose-13C12 were obtained from Sigma (St. Louis, MO, USA). The sample was shaken with two tungsten beads in a mixer mill at 30 Hz for 3 minutes, the beads were removed, and the sample was centrifuged at +4 °C, 14 000 rpm (18 620g), for 10 minutes. The supernatant, 100µL for LC-MS analysis and 50µL for GC-MS analysis, was transferred to micro vials and evaporated to dryness in a speed-vac concentrator. Solvents were evaporated and the samples were stored at -80 °C until analysis. A small aliquot of the remaining supernatants was pooled and used to create quality control (QC) samples. MSMS analysis (LC-MS) was run on the QC samples for identification purposes. The samples were analyzed in batches according to a randomized run order on both GC-MS and LC-MS.

***GC-MS analysis*:** Derivatization and GC-MS analysis were performed as described previously (Gullberg et al. 2004). 0.5 µL of the derivatized sample was injected in split less mode by a L-PAL3 autosampler (CTC Analytics AG, Switzerland) into an Agilent 7890B gas chromatograph equipped with a 10 m x 0.18 mm fused silica capillary column with a chemically bonded 0.18 µm Rxi-5 Sil MS stationary phase (Restek Corporation, U.S.) The injector temperature was 270 °C, the purge flow rate was 20 mL min-1 and the purge was turned on after 60 seconds. The gas flow rate through the column was 1 mL min-1, the column temperature was held at 70 °C for 2 minutes, then increased by 40 °C min-1 to 320 °C, and held there for 2 minutes. The column effluent was introduced into the ion source of a Pegasus BT time-of-flight mass spectrometer, GC/TOFMS (Leco Corp., St Joseph, MI, USA). The transfer line and the ion source temperatures were 250 °C and 200 °C, respectively. Ions were generated by a 70 eV electron beam at an ionization current of 2.0 mA, and 30 spectra s-1 were recorded in the mass range m/z 50 - 800. The acceleration voltage was turned on after a solvent delay of 150 seconds. The detector voltage was 1800-2300 V.

***LC-MS analysis***: Before LC-MS analysis the sample was re-suspended in 10 + 10 µL methanol and water. Each batch of samples was first analyzed in positive mode. After all samples within a batch had been analyzed, the instrument was switched to negative mode and a second injection of each sample was performed. The chromatographic separation was performed on an Agilent 1290 Infinity UHPLC-system (Agilent Technologies, Waldbronn, Germany). 2 µL of each sample were injected onto an Acquity UPLC HSS T3, 2.1 x 50 mm, 1.8 µm C18 column in combination with a 2.1 mm x 5 mm, 1.8 µm VanGuard precolumn (Waters Corporation, Milford, MA, USA) held at 40 °C. The gradient elution buffers were A (H2O, 0.1 % formic acid) and B (75/25 acetonitrile:2-propanol, 0.1 % formic acid), and the flow-rate was 0.5 mL min-1. The compounds were eluted with a linear gradient consisting of 0.1 - 10 % B over 2 minutes, B was increased to 99 % over 5 minutes and held at 99 % for 2 minutes; B was decreased to 0.1 % for 0.3 minutes and the flow-rate was increased to 0.8 mL min-1 for 0.5 minutes; these conditions were held for 0.9 minutes, after which the flow-rate was reduced to 0.5 mL min-1 for 0.1 minutes before the next injection.

The compounds were detected with an Agilent 6546 Q-TOF mass spectrometer equipped with a jet stream electrospray ion source operating in positive or negative ion mode. The settings were kept identical between the modes, with exception of the capillary voltage. A reference interface was connected for accurate mass measurements; the reference ions purine (4 µM) and HP-0921 (Hexakis(1H, 1H, 3H-tetrafluoropropoxy)phosphazine) (1 µM) were infused directly into the MS at a flow rate of 0.05 mL min-1 for internal calibration, and the monitored ions were purine m/z 121.05 and m/z 119.03632; HP-0921 m/z 922.0098 and m/z 966.000725 for positive and negative mode respectively. The gas temperature was set to 150°C, the drying gas flow to 8 L min-1 and the nebulizer pressure 35 psig. The sheath gas temp was set to 350°C and the sheath gas flow 11 L min-1. The capillary voltage was set to 4000 V in positive ion mode, and to 4000 V in negative ion mode. The nozzle voltage was 300 V. The fragmentor voltage was 120 V, the skimmer 65 V and the OCT 1 RF Vpp 750 V. The collision energy was set to 0 V. The m/z range was 70 - 1700, and data was collected in centroid mode with an acquisition rate of 4 scans s-1 (1977 transients/spectrum).

***Data analysis***: For the GC-MS data, all non-processed MS-files from the metabolic analysis were exported from the ChromaTOF software in NetCDF format to MATLAB R2021a (Mathworks, Natick, MA, USA), where all data pre-treatment procedures, such as base-line correction, chromatogram alignment, data compression and Multivariate Curve Resolution were performed. The extracted mass spectra were identified by comparisons of their retention index and mass spectra with libraries of retention time indices and mass spectra (Schauer et al. 2005). Mass spectra and retention index comparison was performed using NIST MS 2.2 software. Annotation of mass spectra was based on reverse and forward searches in the library. Masses and ratio between masses indicative of a derivatized metabolite were especially notified. The mass spectrum with the highest probability indicative of a metabolite and the retention index between the sample and library for the suggested metabolite was ± 5 (usually less than 3) the deconvoluted "peak" was annotated as an identification of a metabolite.

For the LC-MS data, all data processing was performed using the Agilent Masshunter Profinder version B.10.0.2 (Agilent Technologies Inc., Santa Clara, CA, USA). The processing was performed both in a target and an untargeted fashion. For target processing, a pre-defined list of metabolites was searched for using the Batch Targeted feature extraction in Masshunter Profinder. An in-house LC-MS library built up by authentic standards run on the same system with the same chromatographic and mass-spec settings, were used for the targeted processing, in addition an in-house library for plant metabolites. The identification of the metabolites was based on MS, MSMS and retention time information. Batch Recursive Feature Extraction algorithm within Masshunter Profinder, was used for the untargeted data pre-processing.

***Chemicals***:

Solvents: Methanol, HPLC-grade was obtained from Fischer Scientific (Waltham, MA, USA) Chloroform, Suprasolv for GC was obtained from Merck (Darmstadt, Germany) Acetonitrile, HPLC-grade was obtained from Fischer Scientific (Waltham, MA, USA) 2-Propanol, HPLC-grade was obtained from VWR (Radnor, PA, USA) H2O, Milli-Q. Reference and tuning standards: Purine, 4 µM, Agilent Technologies (Santa Clara, CA, USA) HP-0921 (Hexakis(1H, 1H, 3Htetrafluoropropoxy)phosphazine), 1 µM, Agilent Technologies (Santa Clara, CA, USA) Calibrant, ESI-TOF, ESI-L Low Concentration Tuning Mix, Agilent Technologies (Santa Clara, CA, USA) HP-0321 (Hexamethoxyphosphazine), 0.1 mM, Agilent Technologies (Santa Clara, CA, USA).
