## Supplementary file2 for "In *silico* characterisation of PAL homologs and metabolomic profiling of shade response indicates potential presence of PTALs and Tyr as a probable precursor of lignin biosynthesis in conifers"

**Near infrared spectroscopy (NIR)**

The analyses for estimating concentration of wood chemical components (lignin (LIG), cellulose (CELL), and hemicelluloses (HEM) [%]) were based on increment cores. Wet chemical determination of the macromolecular constituents of the wood samples was performed at MoRe Research, using a subset of samples which were selected with the aim of covering as much variability in the wood as possible. The carbohydrates and lignin content for all samples were analysed by the SCAN-CM 71:09 and Tappi T222 methods, respectively. With the SCAN-CM 71:09 method, the different sugar monomers were obtained and from the monomer content, the percentage of cellulose and hemicellulose were quantified, following the formula developed by Sjöström (Sjöström 1993) for softwoods. Near infrared (NIR, 960 – 2500 nm) spectra were collected at 6 nm intervals with a hyperspectral imaging camera. The chemical data were associated with corresponding NIR spectra and partial least squares multivariate models were created and validated with the data. The coefficient of determination ($R^2$) and root mean square error of cross-validation (RMSEcv) for each model are given in Table 1.

Table 1: Performance metrics of the chemistry models employed in the study

| Model | $R^2$ | RMSEcv |
|---|---|---|
| Cellulose | 0.76 | 1.92 |
| Hemicellulose | 0.79 | 1.40 |
| Lignin | 0.70 | 1.39 |

The SilviScan samples were then polished and scanned with the NIR-camera at 30 µm radial resolution on one longitudinal *x* radial surface, facing the side of the tracheids, and the chemical composition was predicted for all samples. The chemical composition at each point in the sample was spatially matched to the physical characteristics as determined by SilviScan, using an algorithm designed for the purpose, which allowed calculating ring-mean averages of the chemical composition.“
