## Supplementary file3 for "In *silico* characterisation of PAL homologs and metabolomic profiling of shade response indicates potential presence of PTALs and Tyr as a probable precursor of lignin biosynthesis in conifers"

| Table S1 Number of metabolites detected in Norway spruce and Scots pine in response to shade | | | | |
|---|---|---|---|---|
| Species | Processing type | Metabolites detected | Significantly down-regulated metabolites | Significantly up-regulated metabolites |
| Spruce North | LCMS | 799 | 141 (identified + non identified) | 45 (identified + non identified) |
| Spruce South | LCMS | 799 | 137 (identified + non identified) | 58 (identified + non identified) |
| Pine North | LCMS | 781 | 264 (identified + non identified) | 73 (identified + non identified) |
| Pine South | LCMS | 781 | 218 (identified + non identified) | 69 (identified + non identified) |
| Spruce North | GCMS | 69 | 15 (identified) | 3 (identified) |
| Spruce South | GCMS | 69 | 13 (identified) | 9 (identified) |
| Pine North | GCMS | 68 | 18 (identified) | 15 (identified) |
| Pine South | GCMS | 68 | 16 (identified) | 13 (identified) |

| Table S2 Metabolites detected in Norway Spruce_north: Sun vs Shade | | | | |
|---|---|---|---|---|
| **Compound Name** | **HMDB** | **KEGG** | **Direction in Shade** | **Pathway** |
| (±)-Taxifolin | HMDB0303943 | C01617 | Down | |
| 1-Aminocyclopropanecarboxylic acid | HMDB0036458 | C01234 | Up | |
| 4-Guanidinobutanoic acid | HMDB0003464 | C01035 | Down | Arginine and proline metabolism |
| Abietic acid | | C06087 | Down | |
| Alanine | HMDB0000161 | C00041 | Up | Alanine, Aspartate, Glutamate metabolism<br>Carbon fixation in photosynthetic organisms |
| alpha-Tocopherol | HMDB0001893 | C02477 | Down | |
| Benzyl-HCH fr | HMDB0304271 | | Down | |
| beta-Alanine | HMDB0000056 | C00099 | Up | beta-Alanine metabolism<br>Pantothenate and CoA biosynthesis |
| Betaine | HMDB0000043 | C00719 | Up | Glycine, serine and threonine metabolism |
| Catechin | HMDB0002780 | C06562 | Down | |
| chiro-Inositol | HMDB0240209 | C19891 | Down | |
| Cinnamoyl acetate fr | | | Down | |
| Dehydroascorbic acid (DHAA) | HMDB0001264 | C00425 | Down | |
| Dihydrouracil | HMDB0000076 | C00429 | Up | beta-Alanine metabolism<br>Pantothenate and CoA biosynthesis |
| Dihydroxy benzoic acid fr1 | | | Down | |
| Dihydroxy benzoic acid fr2 | | | Down | |
| Dihydroxy benzoic acid fr3 | | | Down | |
| Epigallocatechin dimer | | | Down | |
| Eriodictyol glucoside 1 | HMDB0304673 | | Down | |
| Fucose | HMDB0029196 | C01018 | Down | |
| Galactinol | HMDB0005826 | C01235 | Down | Galactose metabolism |
| Gallocatechin | HMDB0038365 | C12127 | Down | |
| gamma-Aminobutyric acid (GABA) | HMDB0000112 | C00334 | Up | Alanine, aspartate and glutamate metabolism<br>Butanoate metabolism<br>Arginine and proline metabolism |
| Indoleacetaldehyde | HMDB0001190 | C00637 | Up | |
| L-Arginine | HMDB0000517 | C00062 | Up | Arginine biosynthesis<br>Arginine and proline metabolism |
| L-Asparagine | HMDB0000168 | C00152 | Up | Alanine, aspartate and glutamate metabolism<br>Cyanoamino acid metabolism |
| L-Aspartic acid | HMDB0000191 | C00049 | Up | Alanine, aspartate and glutamate metabolism<br>Arginine biosynthesis<br>Glycine, serine and threonine metabolism<br>beta-Alanine metabolism<br>Cyanoamino acid metabolism<br>Carbon fixation in photosynthetic organisms<br>Glycolysis / Gluconeogenesis<br>Cysteine and methionine metabolism |
| L-Histidine | HMDB0000177 | C00135 | Up | |
| L-Leucine / L-Isoleucine | HMDB0000687 | C00123 | Up | Valine, leucine and isoleucine biosynthesis |
| L-Lysine | HMDB0000182 | C00047 | Up | |
| L-Proline | HMDB0000162 | C00148 | Up | Arginine and proline metabolism |
| L-Serine | HMDB0000187 | C00065 | Up | Glycine, serine and threonine metabolism<br>Cyanoamino acid metabolism<br>Glyoxylate and dicarboxylate metabolism<br>Cysteine and methionine metabolism |
| L-Threonine | HMDB0000167 | C00188 | Up | Glycine, serine and threonine metabolism<br>Valine, leucine and isoleucine biosynthesis |
| LysoPE(18:2(9Z,12Z)/0:0) | HMDB0011507 | | Down | |
| Ornithine | HMDB0000214 | C00077 | Up | Arginine and proline metabolism<br>Arginine biosynthesis |

| | | | | |
|---|---|---|---|---|
| Oxoglutaric acid | HMDB0000208 | C00026 | Down | Alanine, aspartate and glutamate metabolism<br>Arginine biosynthesis<br>Butanoate metabolism<br>Glyoxylate and dicarboxylate metabolism<br>TCA cycle<br>Glycolysis/Gluconeogenesis |
| Phosphorylcholine | HMDB0001565 | C00588 | Up | |
| Procyanidin B-type dimer 3 | HMDB0033974 | | Down | |
| Procyanidin B-type trimer 3 | | | Down | |
| Pyroglutamic acid | HMDB0000267 | C01879 | Up | |
| Pyruvic acid | HMDB0000243 | C00022 | Down | Alanine, aspartate and glutamate metabolism<br>Glycine, serine and threonine metabolism<br>Valine, leucine and isoleucine biosynthesis<br>Butanoate metabolism<br>Pantothenate and CoA biosynthesis<br>TCA cycle<br>Glycolysis/Gluconeogenesis<br>Tyrosine metabolism<br>Pyruvate metabolism<br>Carbon fixation in photosynthetic organisms<br>Cysteine and methionine metabolism |
| Quinic acid | HMDB0003072 | C00296 | Down | |
| Raffinose | HMDB0003213 | C00492 | Down | Galactose metabolism |
| Ribitol | HMDB0000508 | C00474 | Down | |
| Salicyl aldehyde/Benzoic acid fr 1 | HMDB0001870 | C00539 | Down | |
| Salicylic acid fr 3 | HMDB0001895 | C00805 | Down | |
| Shikimic acid | HMDB0003070 | C00493 | Down | Phenylalanine, tyrosine and tryptophan biosynthesis |
| Sucrose | HMDB0000258 | C00089 | Down | Galactose metabolism<br>Starch and sucrose metabolism<br>Glycolysis/Gluconeogenesis |
| Taxifolin glucopyranoside | | | Down | |
| Threonic acid | HMDB0000943 | C01620 | Up | |
| Valine | HMDB0000883 | C00183 | Up | Valine, leucine and isoleucine biosynthesis<br>Pantothenate and CoA biosynthesis |

| Table S3 Metabolites detected in Norway Spruce_south: Sun vs Shade | | | | |
|---|---|---|---|---|
| Compound Name | HMDB | KEGG | Direction in Shade | Pathway |
| (±)-Taxifolin | HMDB0303943 | C01617 | Down | |
| 1-Aminocyclopropanecarboxylic acid | HMDB0036458 | C01234 | Up | Cysteine and methionine metabolism |
| 2,3,4,5,6,7-Hexahydroxyheptanoic acid | HMDB0240292 | | Down | |
| 2-Ketobutyric acid | HMDB0000005 | C00109 | Down | Valine, leucine and isoleucine biosynthesis<br>Glycine, serine and threonine metabolism<br>Cysteine and methionine metabolism |
| Adenosine | HMDB0000050 | C00212 | Down | |
| Allothreonine | HMDB0004041 | C05519 | Up | Glycine, serine and threonine metabolism |
| Asymmetric dimethylarginine | HMDB0001539 | C03626 | Up | |
| Benzyl-HCH fr | HMDB0304271 | | Down | |
| beta-Cyanoalanine | METPA0300 | C02512 | Up | Cyanoamino acid metabolism |
| Betaine | HMDB0000043 | C00719 | Up | Glycine, serine and threonine metabolism |
| Catechin | HMDB0002780 | C06562 | Down | |
| chiro-Inositol | HMDB0240209 | C19891 | Down | |
| Cinnamoyl acetate fr | | | Down | |
| Dehydroascorbic acid (DHAA) | HMDB0001264 | C00425 | Down | |
| Dihydrouracil | HMDB0000076 | C00429 | Up | beta-Alanine metabolism<br>Pantothenate and CoA biosynthesis |
| Dihydroxy benzoic acid fr1 | | | Down | |
| Dihydroxy benzoic acid fr2 | | | Down | |
| Dihydroxy benzoic acid fr3 | | | Down | |
| Epigallocatechin dimer | | | Down | |
| Fumaric acid | HMDB0000134 | C00122 | Up | Alanine, aspartate and glutamate metabolism<br>TCA cycle<br>Arginine biosynthesis<br>Tyrosine metabolism<br>Pyruvate metabolism<br>Glycolysis / Gluconeogenesis |
| Galactinol | HMDB0005826 | C01235 | Down | Galactose metabolism |
| Galactose 1-phosphate | HMDB0000645 | C00446 | Down | Galactose metabolism |
| Gallocatechin | HMDB0038365 | C12127 | Down | |
| gamma-Aminobutyric acid (GABA) | HMDB0000112 | C00334 | Up | Alanine, aspartate and glutamate metabolism<br>Butanoate metabolism<br>Arginine and proline metabolism |
| gamma-Glutamylleucine / gamma-Glutam | HMDB0011171 | | Up | |
| Guanosine | HMDB0000133 | C00387 | Down | |
| Isoleucine | HMDB0000172 | C00407 | Up | Valine, leucine and isoleucine biosynthesis |
| Isopropylmalate | HMDB0000402 | C02504 | Down | Valine, leucine and isoleucine biosynthesis<br>Pyruvate metabolism |
| Kaempferol | HMDB0005801 | C05903 | Down | Flavone and flavonol biosynthesis<br>Flavonoid biosynthesis |
| L-2-Hydroxyglutaric acid / 3-Hydroxyglutar | HMDB0000694 | C03196 | Down | |
| L-Arginine | HMDB0000517 | C00062 | Up | Arginine biosynthesis<br>Arginine and proline metabolism |
| L-Asparagine | HMDB0000168 | C00152 | Up | Alanine, aspartate and glutamate metabolism<br>Cyanoamino acid metabolism |
| L-Aspartic acid | HMDB0000191 | C00049 | Up | Alanine, aspartate and glutamate metabolism<br>Arginine biosynthesis<br>Glycine, serine and threonine metabolism<br>beta-Alanine metabolism<br>Cyanoamino acid metabolism<br>Carbon fixation in photosynthetic organisms<br>Glycolysis / Gluconeogenesis<br>Cysteine and methionine metabolism |
| Leucyl-Aspartate / gamma-Glutamylvaline | HMDB0028925 | | Up | |

| | | | | |
|---|---|---|---|---|
| L-Leucine / L-Isoleucine | HMDB0000687 | C00123 | Up | Valine, leucine and isoleucine biosynthesis |
| L-Serine | HMDB0000187 | C00065 | Up | Glycine, serine and threonine metabolism<br>Cyanoamino acid metabolism<br>Glyoxylate and dicarboxylate metabolism<br>Cysteine and methionine metabolism |
| L-Threonine | HMDB0000167 | C00188 | Up | Glycine, serine and threonine metabolism<br>Valine, leucine and isoleucine biosynthesis |
| LysoPC(16:0/0:0) | HMDB0010382 | C04230 | Down | |
| LysoPC(18:2(9Z,12Z)/0:0) | HMDB0010386 | C04230 | Down | |
| LysoPC(18:3(9Z,12Z,15Z)/0:0) | HMDB0010388 | C04230 | Down | |
| LysoPE(0:0/18:2(9Z,12Z)) | HMDB0011477 | | Down | |
| Malic acid | HMDB0000156 | C00149 | Up | TCA cycle<br>Glyoxylate and dicarboxylate metabolism<br>Pyruvate metabolism<br>Carbon fixation in photosynthetic organisms<br>Glycolysis / Gluconeogenesis |
| myo-Inositol | HMDB0000211 | C00137 | Down | Galactose metabolism<br>Inositol phosphate metabolism |
| Naringenin-7-O-Glucoside (Prunin) | | C09099 | Down | |
| N-gamma-Glutamylglutamine | HMDB0029147 | | Up | |
| Oxoglutaric acid | HMDB0000208 | C00026 | Down | Alanine, aspartate and glutamate metabolism<br>Arginine biosynthesis<br>Butanoate metabolism<br>Glyoxylate and dicarboxylate metabolism<br>TCA cycle<br>Glycolysis/Gluconeogenesis |
| Pantothenic acid | HMDB0000210 | C00864 | Up | beta-Alanine metabolism<br>Pantothenate and CoA biosynthesis |
| Procyanidin B-type dimer 3 | HMDB0033974 | | Down | |
| Procyanidin B-type trimer 1 | | | Down | |
| Procyanidin B-type trimer 3 | | | Down | |
| Pyroglutamic acid | HMDB0000267 | C01879 | Up | |
| Pyruvic acid | HMDB0000243 | C00022 | Down | Alanine, aspartate and glutamate metabolism<br>Glycine, serine and threonine metabolism<br>Valine, leucine and isoleucine biosynthesis<br>Butanoate metabolism<br>Pantothenate and CoA biosynthesis<br>TCA cycle<br>Glycolysis/Gluconeogenesis<br>Tyrosine metabolism<br>Pyruvate metabolism<br>Carbon fixation in photosynthetic organisms |
| Quinic acid | HMDB0003072 | C00296 | Down | |
| Ribitol | HMDB0000508 | C00474 | Down | |
| Salicyl aldehyde/Benzoic acid fr 1 | HMDB0001870 | C00539 | Down | |
| Salicylic acid fr 3 | HMDB0001895 | C00805 | Down | |
| Shikimic acid | HMDB0003070 | C00493 | Down | Phenylalanine, tyrosine and tryptophan biosynthesis |
| Spermidine | HMDB0001257 | C00315 | Down | beta-Alanine metabolism<br>Arginine and proline metabolism |
| Succinic acid | HMDB0000254 | C00042 | Down | Alanine, aspartate and glutamate metabolism<br>Butanoate metabolism<br>Glyoxylate and dicarboxylate metabolism<br>TCA cycle<br>Glycolysis/Gluconeogenesis |
| Sucrose | HMDB0000258 | C00089 | Down | Galactose metabolism<br>Starch and sucrose metabolism<br>Glycolysis/Gluconeogenesis |
| Taxifolin glucopyranoside | | | Down | |

| | | | | |
|---|---|---|---|---|
| Trigonelline | HMDB0000875 | C01004 | Up | |
| Tyrosine | HMDB0000158 | C00082 | Up | Phenylalanine, tyrosine and tryptophan biosynthesis<br>Tyrosine metabolism<br>Isoquinoline alkaloid biosynthesis |
| Valine | HMDB0000883 | C00183 | Up | Valine, leucine and isoleucine biosynthesis<br>Pantothenate and CoA biosynthesis |
| Vanillic acid fr | HMDB0000484 | C06672 | Down | |
| Vanilloyl glucoside 1 | | | Up | |

| Table S4 Metabolites detected in Scots Pine_north: Sun vs Shade | | | | |
|---|---|---|---|---|
| Compound Name | HMDB | KEGG | Direction in Shade | Pathway |
| (±)-Taxifolin | HMDB0303943 | C01617 | Down | |
| 2,3,4,5,6,7-Hexahydroxyheptanoic acid | HMDB0240292 | | Down | |
| 3-Hydroxyisovaleric acid | HMDB0000754 | | Up | |
| 4-Guanidinobutanoic acid | HMDB0003464 | C01035 | Down | Arginine and proline metabolism |
| Abietic acid | | C06087 | Down | |
| Adenosine | HMDB0000050 | C00212 | Down | |
| Allothreonine | HMDB0004041 | C05519 | Up | Glycine, serine and threonine metabolism |
| Benzyl hexoside pentoside 2 | | | Down | |
| Benzyl-HCH fr | HMDB0304271 | | Down | |
| beta-Cyanoalanine | METPA0300 | C02512 | Up | Cyanoamino acid metabolism |
| Betaine | HMDB0000043 | C00719 | Up | Glycine, serine and threonine metabolism |
| beta-Sitosterol | HMDB0000852 | C01753 | Down | |
| Caffeoyl glucose 3 | HMDB0036937 | C10433 | Down | |
| Caffeoyl quinic acid 4 | HMDB0003164 | C00852/ | Down | Phenylpropanoid biosynthesis<br>Flavonoid biosynthesis |
| Caffeoylshikimate | HMDB0304282 | | Down | |
| Campesterol | HMDB0002869 | C01789 | Down | |
| Catechin | HMDB0002780 | C06562 | Down | |
| chiro-Inositol | HMDB0240209 | C19891 | Down | |
| Cinnamoyl acetate fr | | | Down | |
| Citrulline | HMDB0000904 | C00327 | Up | Arginine biosynthesis |
| Coumaroyl quinic acid 2 | HMDB0029681 | C12208 | Down | Phenylpropanoid biosynthesis<br>Flavonoid biosynthesis |
| Cyanidin 3-glucoside | HMDB0030684 | | Down | Anthocyanin biosynthesis |
| Dehydroabietate | HMDB0061925 | C12078 | Down | |
| Dehydroascorbic acid (DHAA) | HMDB0001264 | C00425 | Down | |
| Dehydromyricetin fr | | | Up | |
| Dihydrouracil | HMDB0000076 | C00429 | Up | beta-Alanine metabolism<br>Pantothenate and CoA biosynthesis |
| Dihydroxy benzoic acid fr1 | | | Down | |
| Dihydroxy benzoic acid fr2 | | | Down | |
| Dihydroxy benzoic acid fr3 | | | Down | |
| Epigallocatechin dimer | | | Down | |
| Erythrose | HMDB0002649 | C01796 | Down | |
| Ferulic acid fr | HMDB0000954 | C01494 | Up | Phenylpropanoid biosynthesis |
| Fructose 6-phosphate | HMDB0000124 | C00085 | Down | Starch and sucrose metabolism<br>Pentose phosphate pathway<br>Glycolysis/Gluconeogenesis<br>Fructose and mannose metabolism |
| Fucose | HMDB0029196 | C01018 | Down | |
| Galactinol | HMDB0005826 | C01235 | Down | Galactose metabolism |
| Galactose 1-phosphate | HMDB0000645 | C00446 | Down | Galactose metabolism |
| Gallocatechin | HMDB0038365 | C12127 | Down | |
| gamma-Aminobutyric acid (GABA) | HMDB0000112 | C00334 | Up | Alanine, aspartate and glutamate metabolism<br>Butanoate metabolism<br>Arginine and proline metabolism |
| Gluconic acid | HMDB0000625 | C00257 | Down | Pentose phosphate pathway<br>Glycolysis/Gluconeogenesis |
| Glucose | HMDB0000122 | C00221 | Down | Galactose metabolism<br>Starch and sucrose metabolism<br>Glycolysis/Gluconeogenesis |
| Glucose 6-phosphate | HMDB0001401 | C00092 | Down | Starch and sucrose metabolism<br>Glycolysis/Gluconeogenesis |
| Glutathione (GSSG) | HMDB0003337 | C00127 | Down | |
| Indoleacetaldehyde | HMDB0001190 | C00637 | Up | Tryptophan metabolism |

| | | | | |
|---|---|---|---|---|
| Isoleucine | HMDB0000172 | C00407 | Up | Valine, leucine and isoleucine biosynthesis |
| Isopropylmalate | HMDB0000402 | C02504 | Down | Valine, leucine and isoleucine biosynthesis<br>Pyruvate metabolism |
| Kaempferol | HMDB0005801 | C05903 | Down | Flavone and flavonol biosynthesis<br>Flavonoid biosynthesis |
| Kaempferol 3-O-b -rutinoside (Nicotiflorin) | | | Down | |
| L-Arginine | HMDB0000517 | C00062 | Up | Arginine biosynthesis<br>Arginine and proline metabolism |
| L-Asparagine | HMDB0000168 | C00152 | Up | Alanine, aspartate and glutamate metabolism<br>Cyanoamino acid metabolism |
| L-Aspartic acid | HMDB0000191 | C00049 | Down | Alanine, aspartate and glutamate metabolism<br>Arginine biosynthesis<br>Glycine, serine and threonine metabolism<br>beta-Alanine metabolism<br>Cyanoamino acid metabolism<br>Carbon fixation in photosynthetic organisms<br>Glycolysis / Gluconeogenesis<br>Cysteine and methionine metabolism |
| Leucyl-Aspartate / gamma-Glutamylvaline | HMDB0028925 | | Up | |
| L-Glutamine | HMDB0000641 | C00064 | Down | Alanine, aspartate and glutamate metabolism<br>Arginine biosynthesis<br>Glyoxylate and dicarboxylate metabolism |
| L-Histidine | HMDB0000177 | C00135 | Up | |
| Linoleic acid | HMDB0000673 | C01595 | Down | Linoleic acid metabolism |
| L-Leucine / L-Isoleucine | HMDB0000687 | C00123 | Up | Valine, leucine and isoleucine biosynthesis |
| L-Lysine | HMDB0000182 | C00047 | Up | |
| L-Phenylalanine | HMDB0000159 | C00079 | Up | Cyanoamino acid metabolism<br>Phenylalanine, tyrosine and tryptophan biosynthesis<br>Phenylpropanoid biosynthesis<br>Phenylalanine metabolism |
| L-Serine | HMDB0000187 | C00065 | Up | Glycine, serine and threonine metabolism<br>Cyanoamino acid metabolism<br>Glyoxylate and dicarboxylate metabolism<br>Cysteine and methionine metabolism |
| L-Threonine | HMDB0000167 | C00188 | Up | Glycine, serine and threonine metabolism<br>Valine, leucine and isoleucine biosynthesis |
| LysoPC(0:0/18:3) | | | Down | |
| LysoPC(16:0/0:0) | HMDB0010382 | C04230 | Down | |
| LysoPC(18:1(9Z)/0:0) | HMDB0002815 | C04230 | Down | |
| LysoPC(18:2(9Z,12Z)/0:0) | HMDB0010386 | C04230 | Down | |
| LysoPE(0:0/18:2(9Z,12Z)) | HMDB0011477 | C04230 | Down | |
| LysoPE(16:0/0:0) | HMDB0011503 | | Down | |
| Oxoglutaric acid | HMDB0000208 | C00026 | Down | Alanine, aspartate and glutamate metabolism<br>Arginine biosynthesis<br>Butanoate metabolism<br>Glyoxylate and dicarboxylate metabolism<br>TCA cycle<br>Glycolysis/Gluconeogenesis |
| Palatinose | | C01742 | Down | |
| Palmitic acid | HMDB0000220 | C00249 | Down | |
| Pelargonic acid | HMDB0000847 | C01601 | Up | |
| Phosphorylcholine | HMDB0001565 | C00588 | Up | |
| Procyanidin B-type dimer 1 | HMDB0029754 | | Down | |
| Procyanidin B-type dimer 2 | HMDB0033973 | | Down | |
| Procyanidin B-type dimer 3 | HMDB0033974 | | Down | |
| Procyanidin B-type trimer 1 | | | Down | |
| Procyanidin B-type trimer 2 | | | Down | |

| | | | | |
|---|---|---|---|---|
| Protocatechuic acid glucoside | HMDB0303826 | | Up | |
| Pyruvic acid | HMDB0000243 | C00022 | Down | Alanine, aspartate and glutamate metabolism<br>Glycine, serine and threonine metabolism<br>Valine, leucine and isoleucine biosynthesis<br>Butanoate metabolism<br>Pantothenate and CoA biosynthesis<br>TCA cycle<br>Glycolysis/Gluconeogenesis<br>Tyrosine metabolism<br>Pyruvate metabolism<br>Carbon fixation in photosynthetic organisms |
| Quercetin fr 2 | | | Up | |
| Quinic acid | HMDB0003072 | C00296 | Down | |
| Ribitol | HMDB0000508 | C00474 | Down | |
| Salicyl aldehyde/Benzoic acid fr 1 | HMDB0001870 | C00539 | Down | |
| Salicylic acid fr 3 | HMDB0001895 | C00805 | Down | |
| Salirepin 1 | | | Down | |
| Shikimic acid | HMDB0003070 | C00493 | Down | Phenylalanine, tyrosine and tryptophan biosynthesis |
| Spermidine | HMDB0001257 | C00315 | Down | beta-Alanine metabolism<br>Arginine and proline metabolism |
| Succinic acid | HMDB0000254 | C00042 | Down | Alanine, aspartate and glutamate metabolism<br>Butanoate metabolism<br>Glyoxylate and dicarboxylate metabolism<br>TCA cycle<br>Glycolysis/Gluconeogenesis |
| Sucrose | HMDB0000258 | C00089 | Down | Galactose metabolism<br>Starch and sucrose metabolism<br>Glycolysis/Gluconeogenesis |
| Threonic acid | HMDB0000943 | C01620 | Up | |
| Tryptophan | HMDB0000929 | C00078 | Up | Glycine, serine and threonine metabolism<br>Phenylalanine, tyrosine and tryptophan biosynthesis<br>Tryptophan metabolism |
| Tyrosine | HMDB0000158 | C00082 | Up | Phenylalanine, tyrosine and tryptophan biosynthesis<br>Tyrosine metabolism<br>Isoquinoline alkaloid biosynthesis |
| Valine | HMDB0000883 | C00183 | Up | Valine, leucine and isoleucine biosynthesis<br>Pantothenate and CoA biosynthesis |
| Xanthone der 1 | HMDB0029463 | C10065 | Up | |

| Table S5 Metabolites detected in Scots Pine_south: Sun vs Shade | | | | |
|---|---|---|---|---|
| **Compound Name** | **HMDB** | **KEGG** | **Direction in Shade** | **Pathway** |
| (±)-Taxifolin | HMDB0303943 | C01617 | Down | |
| 2,3,4,5,6,7-Hexahydroxyheptanoic ac | HMDB0240292 | | Down | |
| 2-Hydroxyhexadecanoic acid | HMDB0031057 | | Down | |
| 3-Hydroxyisovaleric acid | HMDB0000754 | | Up | |
| 4-Guanidinobutanoic acid | HMDB0003464 | C01035 | Down | Arginine and proline metabolism |
| 5-Hydroxy-L-tryptophan | HMDB0000472 | C00643 | Up | Tryptophan metabolism |
| Abietic acid | | C06087 | Down | |
| Allothreonine | HMDB0004041 | C05519 | Up | Glycine, serine and threonine metabolism |
| Asymmetric dimethylarginine | HMDB0001539 | C03626 | Up | |
| Benzyl-HCH fr | HMDB0304271 | | Down | |
| beta-Cyanoalanine | METPA0300 | C02512 | Up | Cyanoamino acid metabolism |
| Betaine | HMDB0000043 | C00719 | Up | Glycine, serine and threonine metabolism |
| Caffeoyl glucose 3 | HMDB0036937 | C10433 | Down | |
| Caffeoyl quinic acid 4 | HMDB0003164 | C00852/( | Down | Phenylpropanoid biosynthesis<br>Flavonoid biosynthesis |
| Caffeoylshikimate | HMDB0304282 | | Down | |
| Catechin | HMDB0002780 | C06562 | Down | |
| chiro-Inositol | HMDB0240209 | C19891 | Down | |
| Cinnamoyl acetate fr | | | Down | |
| Citrulline | HMDB0000904 | C00327 | Up | Arginine biosynthesis |
| Coumaroyl quinic acid 2 | HMDB0029681 | C12208 | Down | Phenylpropanoid biosynthesis<br>Flavonoid biosynthesis |
| Coumaroyl quinic acid 3 | HMDB0301709 | C12208 | Down | Phenylpropanoid biosynthesis<br>Flavonoid biosynthesis |
| Dehydroabietate | HMDB0061925 | C12078 | Down | |
| Dehydroascorbic acid (DHAA) | HMDB0001264 | C00425 | Down | |
| D-Glucuronic acid | HMDB0000127 | C00191 | Down | Pentose and glucuronate interconversions |
| Dihydrouracil | HMDB0000076 | C00429 | Up | beta-Alanine metabolism<br>Pantothenate and CoA biosynthesis |
| Dihydroxy benzoic acid fr1 | | | Down | |
| Dihydroxy benzoic acid fr2 | | | Down | |
| Dihydroxy benzoic acid fr3 | | | Down | |
| Dimethoxybenzaldehyde fr 2 | | | Down | |
| D-threo-Isocitric acid | HMDB0001874 | C00451 | Down | |
| Epigallocatechin dimer | | | Down | |
| Ferulic acid fr | HMDB0000954 | C01494 | Up | Phenylpropanoid biosynthesis |
| Fructose | HMDB0000660 | C02336 | Down | Galactose metabolism<br>Starch and sucrose metabolism<br>Glycolysis/Gluconeogenesis<br>Fructose and mannose metabolism |
| Fucose | HMDB0029196 | C01018 | Down | |
| Galactinol | HMDB0005826 | C01235 | Down | Galactose metabolism |
| Galactose 1-phosphate | HMDB0000645 | C00446 | Down | Galactose metabolism |
| Gallocatechin | HMDB0038365 | C12127 | Down | |
| gamma-Aminobutyric acid (GABA) | HMDB0000112 | C00334 | Up | Alanine, aspartate and glutamate metabolism<br>Butanoate metabolism<br>Arginine and proline metabolism |
| gamma-Glutamylleucine / gamma-Gl | HMDB0011171 | | Up | |
| Gluconic acid | HMDB0000625 | C00257 | Down | |
| Glucose | HMDB0000122 | C00221 | Down | Galactose metabolism<br>Starch and sucrose metabolism<br>Glycolysis/Gluconeogenesis |

| | | | | |
|---|---|---|---|---|
| Indoleacetaldehyde | HMDB0001190 | C00637 | Up | Tryptophan metabolism |
| Isopropylmalate | HMDB0000402 | C02504 | Down | Valine, leucine and isoleucine biosynthesis<br>Pyruvate metabolism |
| Kaempferol | HMDB0005801 | C05903 | Down | Flavone and flavonol biosynthesis<br>Flavonoid biosynthesis |
| Kaempferol 3-O-b -rutinoside (Nicotiflorin) | | | Down | |
| Kaempferol-7-o-glucoside | | | Down | |
| Kaempherol fr | | | Down | |
| L-Arginine | HMDB0000517 | C00062 | Up | Arginine biosynthesis<br>Arginine and proline metabolism |
| L-Asparagine | HMDB0000168 | C00152 | Up | Alanine, aspartate and glutamate metabolism<br>Cyanoamino acid metabolism |
| L-Aspartic acid | HMDB0000191 | C00049 | Down | Alanine, aspartate and glutamate metabolism<br>Arginine biosynthesis<br>Glycine, serine and threonine metabolism<br>beta-Alanine metabolism<br>Cyanoamino acid metabolism<br>Carbon fixation in photosynthetic organisms<br>Glycolysis / Gluconeogenesis<br>Cysteine and methionine metabolism |
| Leucyl-Aspartate / gamma-Glutamylv | HMDB0028925 | | Up | |
| L-Glutamine | HMDB0000641 | C00064 | Down | Alanine, aspartate and glutamate metabolism<br>Arginine biosynthesis<br>Glyoxylate and dicarboxylate metabolism |
| L-Histidine | HMDB0000177 | C00135 | Up | |
| L-Leucine / L-Isoleucine | HMDB0000687 | C00123 | Up | Valine, leucine and isoleucine biosynthesis |
| L-Lysine | HMDB0000182 | C00047 | Up | |
| L-Phenylalanine | HMDB0000159 | C00079 | Up | Cyanoamino acid metabolism<br>Phenylalanine, tyrosine and tryptophan biosynthesis<br>Phenylpropanoid biosynthesis<br>Phenylalanine metabolism |
| L-Serine | HMDB0000187 | C00065 | Up | Glycine, serine and threonine metabolism<br>Cyanoamino acid metabolism<br>Glyoxylate and dicarboxylate metabolism<br>Cysteine and methionine metabolism |
| L-Threonine | HMDB0000167 | C00188 | Up | Glycine, serine and threonine metabolism<br>Valine, leucine and isoleucine biosynthesis |
| LysoPC(16:0/0:0) | HMDB0010382 | C04230 | Down | |
| LysoPC(18:1(9Z)/0:0) | HMDB0002815 | C04230 | Down | |
| LysoPC(18:2(9Z,12Z)/0:0) | HMDB0010386 | C04230 | Down | |
| LysoPE(0:0/18:2(9Z,12Z)) | HMDB0011477 | | Down | |
| LysoPE(16:0/0:0) | HMDB0011503 | | Down | |
| myo-Inositol | HMDB0000211 | C00137 | Down | Galactose metabolism |
| Myo-Inositol 1-Monophosphate | HMDB0000213 | C01177 | Down | |
| N-gamma-Glutamylglutamine | HMDB0029147 | | Down | |
| Ornithine | HMDB0000214 | C00077 | Up | Arginine biosynthesis<br>Arginine and proline metabolism |
| Oxoglutaric acid | HMDB0000208 | C00026 | Down | Alanine, aspartate and glutamate metabolism<br>Arginine biosynthesis<br>Butanoate metabolism<br>Glyoxylate and dicarboxylate metabolism<br>TCA cycle<br>Glycolysis/Gluconeogenesis |

| | | | | |
|---|---|---|---|---|
| Pantothenic acid | HMDB0000210 | C00864 | Up | beta-Alanine metabolism<br>Pantothenate and CoA biosynthesis |
| Pelargonic acid | HMDB0000847 | C01601 | Up | |
| Phosphorylcholine | HMDB0001565 | C00588 | Up | |
| Procyanidin B-type dimer 1 | HMDB0029754 | | Down | |
| Procyanidin B-type dimer 2 | HMDB0033973 | | Down | |
| Procyanidin B-type dimer 3 | HMDB0033974 | | Down | |
| Procyanidin B-type trimer 1 | | | Down | |
| Procyanidin B-type trimer 2 | | | Down | |
| Pyroglutamic acid | HMDB0000267 | C01879 | Down | |
| Pyruvic acid | HMDB0000243 | C00022 | Down | Alanine, aspartate and glutamate metabolism<br>Glycine, serine and threonine metabolism<br>Valine, leucine and isoleucine biosynthesis<br>Butanoate metabolism<br>Pantothenate and CoA biosynthesis<br>TCA cycle<br>Glycolysis/Gluconeogenesis<br>Tyrosine metabolism<br>Pyruvate metabolism<br>Carbon fixation in photosynthetic organisms |
| Quinic acid | HMDB0003072 | C00296 | Down | |
| Ribitol | HMDB0000508 | C00474 | Down | |
| Salicyl aldehyde/Benzoic acid fr 1 | HMDB0001870 | C00539 | Down | |
| Salicylic acid fr 3 | HMDB0001895 | C00805 | Down | |
| Salirepin 1 | | | Down | |
| Shikimic acid | HMDB0003070 | C00493 | Down | Phenylalanine, tyrosine and tryptophan biosynthesis |
| Spermidine | HMDB0001257 | C00315 | Down | beta-Alanine metabolism<br>Arginine and proline metabolism |
| Succinic acid | HMDB0000254 | C00042 | Down | Alanine, aspartate and glutamate metabolism<br>Butanoate metabolism<br>Glyoxylate and dicarboxylate metabolism<br>TCA cycle<br>Glycolysis/Gluconeogenesis |
| Sucrose | HMDB0000258 | C00089 | Down | Galactose metabolism<br>Starch and sucrose metabolism<br>Glycolysis/Gluconeogenesis |
| Trigonelline | HMDB0000875 | C01004 | Up | |
| Tryptophan | HMDB0000929 | C00078 | Up | Glycine, serine and threonine metabolism<br>Phenylalanine, tyrosine and tryptophan biosynthesis<br>Tryptophan metabolism |
| Tyrosine | HMDB0000158 | C00082 | Up | Phenylalanine, tyrosine and tryptophan biosynthesis<br>Tyrosine metabolism<br>Isoquinoline alkaloid biosynthesis |
| Valine | HMDB0000883 | C00183 | Up | Valine, leucine and isoleucine biosynthesis<br>Pantothenate and CoA biosynthesis |
| Vanillic acid fr | HMDB0000484 | C06672 | Down | |

**Table S6** **Details of PAL sequences from monocots and putative PTAL sequences from conifers used for the multiple alignment and phylogeny.**

| **Species** | **Sequence Id (TAIR/GenBank/Plaza)** | **Expression: Sun Versus Shade at South and North, respectively** | **Number of amino acids** | **Abbreviation** |
|---|---|---|---|---|
| Arabidopsis thaliana | AT2G37040 | | 725 | AtPAL |
| Brachypodium distachyon | XP_003575400.1 | | 717 | BdPAL |
| Bambusa oldhamii | ACN62413.1 | | 713 | BoPAL |
| Petroselinum crispum | P24481.1 | | 716 | PcPAL |
| Populus trichocarpa | XP_006381441.1 | | 714 | PtPAL |
| Oryza sativa | A2X7F7.1 | | 713 | OsPAL |
| Zea mays | NP_001151482.2 | | 718 | ZmPAL |
| Picea abies | PAB00008810/MA_10429279g0010 | Not significant at latitude South and North | 711 | PabPAL1 |
| | PAB00025303/MA_15852g0010 High confidence | Not significant at latitude South Shade>Sun at latitude North | 787 | PabPAL2 |
| Pseudotsuga menziesii | PME00016510 | | 712 | PmePAL1 |
| | PME00011612 | | 828 | PmePAL2 |
| Picea sitchensis | PSI00008374 | | 790 | PsiPAL |
| Pinus sylvestris | PSY00008254 | | 678 | PsyPAL |
| Pinus taeda | PTA00029770/ PITA_000038676 | Not significant at latitude South and North | 711 | PtaPAL1 |
| | PTA00066008/ PITA_000078355 | Shade<Sun at latitude South Not significant at latitude North | 711 | PtaPAL2 |
| | PTA00062798/ PITA_000074853 | Not significant at latitude South and North | 739 | PtaPAL3 |
| | PTA00046911/ PITA_000057642 | Not significant at latitude South and North | 696 | PtaPAL4 |

**Table S7** **Details of PTAL sequences from monocots and putative PTAL sequences from conifers used for the multiple alignment and phylogeny.**

| **Species** | **Sequence Id (TAIR/GenBank/Plaza)** | **Expression: Sun Versus Shade at South and North, respectively** | **Number of amino acids** | **Abbreviation** |
|---|---|---|---|---|
| Brachypodium distachyon | XP_003575396.1 | | 707 | BdPTAL |
| Bambusa oldhamii | ADE08261.1 | | 701 | BoPTAL |
| Oryza sativa | S06475 | | 701 | OsPTAL |
| Zea mays | NP_001105334.2 | | 703 | ZmPTAL |
| Picea abies | PAB00020676/MA_123220g0010 | Not significant at latitude South and North | 748 | PabPTAL1 |
| | PAB00042267/MA_44561g0010 High confidence | Shade<Sun at latitude South and North | 718 | PabPTAL2 |
| | PAB00052998/MA_73113g0010 | Not significant at latitude South and North | 736 | PabPTAL3 |
| Pseudotsuga menziesii | PME00018594 | | 753 | PmePTAL1 |
| | PME00002674 | | 724 | PmePTAL2 |
| | PME00065720 | | 765 | PmePTAL3 |
| | PME00143027 | | 687 | PmePTAL4 |
| Picea sitchensis | PSI00019215 | | 720 | PsiPTAL |
| Pinus sylvestris | PSY00017079 | | 718 | PsyPTAL |
| Pinus taeda | PTA00031900/ PITA_000041078 | Not significant at latitude South and North | 718 | PtaPTAL1 |
| | PTA00063025/ PITA_000075110 | Not significant at latitude South and North | 764 | PtaPTAL2 |
| | PTA00005492/ PITA_000005593 | Not expressed at latitude South and North | 686 | PtaPTAL3 |

| Table S8 Population-wise allele frequencies of PAL/PTAL SNPs detected in Norway spruce | | | | | | | | | | |
|---|---|---|---|---|---|---|---|---|---|---|
| Gene<br>Spruce gene id | Position | Mutation | Allele | Population-wise allele frequency | | | | | | Cline |
| | | | | S1 | S2 | S3 | S4 | S5 | S6 | |
| **PabPAL2**<br>**MA_15852g0010** | 35746 | missense | Reference (C) | 0.59 | 0.58 | 0.60 | 0.57 | 0.58 | 0.56 | No |
| | | | Alternate (T) | 0.41 | 0.42 | 0.40 | 0.43 | 0.42 | 0.44 | |
| | 35760 | missense | Reference (A) | 0.56 | 0.56 | 0.56 | 0.57 | 0.57 | 0.55 | No |
| | | | Alternate (T) | 0.44 | 0.44 | 0.44 | 0.43 | 0.43 | 0.45 | |
| | 35766 | missense | Reference (T) | 0.56 | 0.56 | 0.57 | 0.57 | 0.56 | 0.56 | No |
| | | | Alternate (G) | 0.44 | 0.44 | 0.43 | 0.43 | 0.44 | 0.44 | |
| | 35776 | missense | Reference (C) | 0.56 | 0.56 | 0.58 | 0.56 | 0.57 | 0.56 | No |
| | | | Alternate (A) | 0.44 | 0.44 | 0.43 | 0.44 | 0.43 | 0.44 | |
| | 35780 | synonymous | Reference (C) | 0.58 | 0.57 | 0.58 | 0.59 | 0.58 | 0.59 | No |
| | | | Alternate (T) | 0.42 | 0.43 | 0.42 | 0.41 | 0.42 | 0.41 | |
| | 35829 | missense | Reference (T) | 0.52 | 0.52 | 0.53 | 0.52 | 0.52 | 0.52 | No |
| | | | Alternate (G) | 0.48 | 0.48 | 0.47 | 0.48 | 0.48 | 0.48 | |
| | 35893 | missense | Reference (C) | 0.82 | 0.81 | 0.81 | 0.82 | 0.87 | 0.83 | No |
| | | | Alternate (T) | 0.18 | 0.19 | 0.19 | 0.18 | 0.13 | 0.17 | |
| | 35924 | synonymous | Reference (A) | 0.51 | 0.51 | 0.51 | 0.52 | 0.52 | 0.51 | No |
| | | | Alternate (G) | 0.49 | 0.49 | 0.49 | 0.48 | 0.48 | 0.49 | |
| | 35929 | missense | Reference (A) | 0.50 | 0.50 | 0.50 | 0.50 | 0.50 | 0.50 | No |
| | | | Alternate (T) | 0.50 | 0.50 | 0.50 | 0.50 | 0.50 | 0.50 | |
| | 35940 | synonymous | Reference (C) | 1.00 | 0.98 | 0.99 | 1.00 | 0.98 | 0.98 | No |
| | | | Alternate (A) | 0.00 | 0.02 | 0.01 | 0.00 | 0.02 | 0.02 | |
| | **36047** | **synonymous** | Reference (T) | **0.50** | **0.50** | **0.50** | **0.49** | **0.48** | **0.47** | **Yes** |
| | | | Alternate (C) | **0.50** | **0.50** | **0.50** | **0.51** | **0.52** | **0.53** | |
| | 36124 | missense | Reference (G) | 0.93 | 0.94 | 0.90 | 0.93 | 0.93 | 0.94 | No |
| | | | Alternate (A) | 0.07 | 0.06 | 0.10 | 0.07 | 0.07 | 0.06 | |
| | 36127 | missense | Reference (A) | 1.00 | 1.00 | 0.99 | 0.93 | 0.92 | 0.95 | No |
| | | | Alternate (G) | 0.00 | 0.00 | 0.01 | 0.07 | 0.08 | 0.05 | |
| | 36132 | missense | Reference (G) | 0.43 | 0.42 | 0.43 | 0.44 | 0.42 | 0.41 | No |
| | | | Alternate (A) | 0.57 | 0.58 | 0.57 | 0.56 | 0.58 | 0.59 | |
| | 36150 | missense | Reference (A) | 0.56 | 0.56 | 0.58 | 0.57 | 0.60 | 0.56 | No |
| | | | Alternate (C) | 0.44 | 0.44 | 0.42 | 0.43 | 0.40 | 0.44 | |
| | 36176 | synonymous | Reference (T) | 0.98 | 0.99 | 0.99 | 0.99 | 0.99 | 1.00 | No |
| | | | Alternate (A) | 0.02 | 0.01 | 0.01 | 0.01 | 0.01 | 0.00 | |
| | 36193 | missense | Reference (G) | 0.66 | 0.69 | 0.69 | 0.67 | 0.69 | 0.65 | No |
| | | | Alternate (C) | 0.34 | 0.31 | 0.31 | 0.33 | 0.31 | 0.35 | |
| | 36197 | synonymous | Reference (G) | 0.66 | 0.67 | 0.68 | 0.67 | 0.69 | 0.64 | No |
| | | | Alternate (A) | 0.34 | 0.33 | 0.32 | 0.33 | 0.31 | 0.36 | |
| | 36230 | synonymous | Reference (T) | 0.74 | 0.75 | 0.79 | 0.74 | 0.74 | 0.68 | No |
| | | | Alternate (C) | 0.26 | 0.25 | 0.21 | 0.26 | 0.26 | 0.32 | |
| | 36236 | synonymous | Reference (T) | 0.78 | 0.80 | 0.84 | 0.81 | 0.77 | 0.72 | No |
| | | | Alternate (C) | 0.22 | 0.20 | 0.16 | 0.19 | 0.23 | 0.28 | |
| | 36257 | synonymous | Reference (G) | 0.87 | 0.89 | 0.86 | 0.82 | 0.83 | 0.76 | No |
| | | | Alternate (A) | 0.13 | 0.11 | 0.14 | 0.18 | 0.17 | 0.24 | |
| | 36338 | synonymous | Reference (C) | 0.79 | 0.81 | 0.77 | 0.70 | 0.72 | 0.68 | No |
| | | | Alternate (T) | 0.21 | 0.19 | 0.23 | 0.30 | 0.28 | 0.32 | |
| | 36347 | synonymous | Reference (A) | 0.40 | 0.40 | 0.40 | 0.42 | 0.43 | 0.40 | No |
| | | | Alternate (G) | 0.60 | 0.60 | 0.60 | 0.58 | 0.57 | 0.60 | |
| | 36356 | synonymous | Reference (C) | 0.78 | 0.80 | 0.78 | 0.70 | 0.70 | 0.68 | No |
| | | | Alternate (T) | 0.22 | 0.20 | 0.22 | 0.30 | 0.30 | 0.32 | |

| Gene<br>Spruce gene id | Position | Mutation | Allele | Population-wise allele frequency | | | | | | Cline |
|---|---|---|---|---|---|---|---|---|---|---|
| | | | | S1 | S2 | S3 | S4 | S5 | S6 | |
| **PabPAL2**<br>**MA_15852g0010** | 36380 | synonymous | Reference (G) | 0.50 | 0.51 | 0.53 | 0.51 | 0.52 | 0.50 | No |
| | | | Alternate (C) | 0.50 | 0.49 | 0.47 | 0.49 | 0.48 | 0.50 | |
| | **36431** | **synonymous** | Reference (G) | **0.97** | **0.96** | **0.96** | **0.95** | **0.92** | **0.86** | **Yes** |
| | | | Alternate (C) | **0.03** | **0.04** | **0.04** | **0.05** | **0.08** | **0.14** | |
| | 36440 | synonymous | Reference (A) | 0.50 | 0.51 | 0.50 | 0.50 | 0.50 | 0.51 | No |
| | | | Alternate (G) | 0.50 | 0.49 | 0.50 | 0.50 | 0.50 | 0.49 | |
| | 36446 | synonymous | Reference (A) | 0.50 | 0.51 | 0.50 | 0.50 | 0.50 | 0.51 | No |
| | | | Alternate (G) | 0.50 | 0.49 | 0.50 | 0.50 | 0.50 | 0.49 | |
| | 36497 | synonymous | Reference (C) | 0.50 | 0.50 | 0.50 | 0.50 | 0.50 | 0.50 | No |
| | | | Alternate (T) | 0.50 | 0.50 | 0.50 | 0.50 | 0.50 | 0.50 | |
| | **36510** | **missense** | Reference (G) | **0.90** | **0.90** | **0.87** | **0.86** | **0.85** | **0.77** | **Yes** |
| | | | Alternate (A) | **0.10** | **0.10** | **0.13** | **0.14** | **0.15** | **0.23** | |
| | 36512 | synonymous | Reference (A) | 0.50 | 0.50 | 0.50 | 0.50 | 0.50 | 0.50 | No |
| | | | Alternate (C) | 0.50 | 0.50 | 0.50 | 0.50 | 0.50 | 0.50 | |
| | 36561 | synonymous | Reference (T) | 0.89 | 0.94 | 0.94 | 0.96 | 0.95 | 0.95 | No |
| | | | Alternate (C) | 0.11 | 0.06 | 0.06 | 0.04 | 0.05 | 0.05 | |
| | 36677 | synonymous | Reference (C) | 0.83 | 0.82 | 0.77 | 0.86 | 0.90 | 0.85 | No |
| | | | Alternate (T) | 0.17 | 0.18 | 0.23 | 0.14 | 0.10 | 0.15 | |
| | 36699 | missense | Reference (T) | 0.98 | 0.95 | 0.96 | 0.96 | 0.97 | 0.98 | No |
| | | | Alternate (A) | 0.02 | 0.05 | 0.04 | 0.04 | 0.03 | 0.02 | |
| **PabPTAL1**<br>**MA_123220g0010** | 3693 | synonymous | Reference (C) | 0.99 | 0.98 | 1.00 | 0.99 | 1.00 | 0.98 | No |
| | | | Alternate (A) | 0.01 | 0.02 | 0.00 | 0.01 | 0.00 | 0.02 | |
| | **3784** | **missense** | Reference (C) | **0.97** | **0.97** | **0.95** | **0.93** | **0.89** | **0.86** | **Yes** |
| | | | Alternate (T) | **0.03** | **0.03** | **0.05** | **0.07** | **0.11** | **0.14** | |
| | **3790** | **missense** | Reference (C) | **0.90** | **0.90** | **0.87** | **0.88** | **0.84** | **0.80** | **Yes** |
| | | | Alternate (T) | **0.10** | **0.10** | **0.13** | **0.12** | **0.16** | **0.20** | |
| | 3805 | missense | Reference (C) | 0.94 | 0.96 | 0.94 | 0.95 | 0.95 | 0.96 | No |
| | | | Alternate (T) | 0.06 | 0.04 | 0.06 | 0.05 | 0.05 | 0.04 | |
| | 3807 | synonymous | Reference (C) | 0.96 | 0.96 | 0.97 | 0.99 | 0.98 | 0.98 | No |
| | | | Alternate (T) | 0.04 | 0.04 | 0.03 | 0.01 | 0.02 | 0.02 | |
| | 3831 | synonymous | Reference (G) | 0.96 | 0.96 | 0.97 | 0.99 | 0.98 | 0.98 | No |
| | | | Alternate (A) | 0.04 | 0.04 | 0.03 | 0.01 | 0.02 | 0.02 | |
| | 3861 | synonymous | Reference (C) | 0.98 | 0.97 | 0.96 | 0.94 | 0.87 | 0.93 | No |
| | | | Alternate (T) | 0.02 | 0.03 | 0.04 | 0.06 | 0.13 | 0.08 | |
| | 3882 | synonymous | Reference (C) | 0.99 | 0.99 | 0.99 | 1.00 | 0.99 | 0.98 | No |
| | | | Alternate (T) | 0.01 | 0.01 | 0.01 | 0.00 | 0.01 | 0.02 | |
| **PabPTAL3**<br>**MA_73113g0010** | 6025 | missense | Reference (A) | 0.93 | 0.96 | 0.94 | 0.99 | 0.98 | 0.97 | No |
| | | | Alternate (G) | 0.07 | 0.04 | 0.06 | 0.01 | 0.02 | 0.03 | |
| | 6122 | missense | Reference (G) | 0.92 | 0.91 | 0.86 | 0.76 | 0.81 | 0.71 | No |
| | | | Alternate (A) | 0.08 | 0.09 | 0.14 | 0.24 | 0.19 | 0.29 | |
| | 6168 | synonymous | Reference (G) | 0.84 | 0.87 | 0.88 | 0.94 | 0.93 | 0.91 | No |
| | | | Alternate (A) | 0.16 | 0.13 | 0.12 | 0.06 | 0.07 | 0.09 | |
| | 6237 | synonymous | Reference (G) | 0.84 | 0.83 | 0.82 | 0.84 | 0.84 | 0.81 | No |
| | | | Alternate (A) | 0.16 | 0.17 | 0.18 | 0.16 | 0.16 | 0.19 | |
| | **6265** | **missense** | Reference (C) | **0.94** | **0.95** | **0.91** | **0.87** | **0.89** | **0.83** | **Yes** |
| | | | Alternate (T) | **0.06** | **0.05** | **0.09** | **0.13** | **0.11** | **0.17** | |
| | **6296** | **missense** | Reference (A) | **0.95** | **0.95** | **0.92** | **0.89** | **0.89** | **0.85** | **Yes** |
| | | | Alternate (T) | **0.05** | **0.05** | **0.08** | **0.11** | **0.11** | **0.15** | |

| Table S9 P-values from one-way ANOVA of the allele frequencies and genotype frequencies of missense SNPs in PAL/PTAL showing cline across the populations in Norway spruce in Sweden | | | | |
|---|---|---|---|---|
| **Gene**<br>**Spruce gene id** | **Variation** | **Reference allele frequency p-value** | **Alternate allele frequency p-value** | **Genotype frequency p-value** |
| **PabPAL2**<br>**MA_15852g0010** | Reference T, alternate C<br>GCT → GCC; Ala228Ala | T: 3.19E-05 | C: 0.0004 | 0.0002 |
| | Reference G, alternate C<br>CCG → CCC; Pro356Pro | G: 3.99E-12 | C: 3.99E-12 | 3.99E-12 |
| | Reference G, alternate A<br>GCA → ACA; Ala383Thr | G: 4.75E-10 | A: 4.75E-10 | 4.75E-10 |
| **PabPTAL1**<br>**MA_123220g0010** | Reference C, alternate T<br>GCC → GTC; Ala535Val | C: 3.34E-06 | T: 1.43E-06 | 1.73E-06 |
| | Reference C, alternate T<br>GCT → GCT; Ala537Val | C: 7.8E-15 | T: 1.56E-15 | 1.47E-15 |
| **PabPTAL3**<br>**MA_73113g0010** | Reference C, alternate T<br>TCC → TTC; Ser595Phe | C: 3.09E-06 | T: 1.3E-09 | 4.12E-09 |
| | Reference A, alternate T<br>TTA → TTT; Leu601Phe | A: 1.95E-06 | T: 7.36E-09 | 1.20E-07 |

**Table S10 One-way ANOVA and Tukey's post-hoc test for genotype frequencies of missense SNPs showing latitudinal cline in Norway spruce in Sweden.**
**Significant p-values (p-value<0.05) are highlighted**

| Gene<br>Spruce gene id | Variation | Tukey's p-value S1 vs S2 | Tukey's p-value S1 vs S3 | Tukey's p-value S1 vs S4 | Tukey's p-value S1 vs S5 | Tukey's p-value S1 vs S6 | Tukey's p-value S2 vs S3 | Tukey's p-value S2 vs S4 | Tukey's p-value S2 vs S5 | Tukey's p-value S2 vs S6 | Tukey's p-value S3 vs S4 | Tukey's p-value S3 vs S5 | Tukey's p-value S3 vs S6 | Tukey's p-value S4 vs S5 | Tukey's p-value S4 vs S6 | Tukey's p-value S5 vs S6 |
|---|---|---|---|---|---|---|---|---|---|---|---|---|---|---|---|---|
| **PabPAL2 MA_15852g0010** | Ala228Ala | 0.89999 | 0.89999 | 0.89999 | 0.14764 | **0.0016** | 0.89999 | 0.89999 | 0.38736 | **0.0085** | 0.89493 | 0.13527 | **0.0018** | 0.77999 | **0.0491** | 0.24361 |
| | Pro356Pro | 0.89999 | 0.89999 | 0.80186 | **0.0014** | **0.001** | 0.89999 | 0.89999 | **0.0358** | **0.001** | 0.89999 | **0.033** | **0.001** | 0.20479 | **0.001** | **0.001** |
| | Ala383Thr | 0.89999 | 0.7354 | 0.42999 | 0.10567 | **0.001** | 0.73089 | 0.43676 | 0.12345 | **0.001** | 0.89999 | 0.89999 | **0.001** | 0.89999 | **0.001** | **0.001** |
| **PabPTAL1 MA_123220g0010** | Ala535Val | 0.89999 | 0.52257 | 0.79511 | **0.0041** | **0.001** | 0.62482 | 0.89213 | **0.0131** | **0.001** | 0.89999 | 0.74499 | 0.06147 | 0.35208 | **0.01** | 0.30369 |
| | Ala537Val | 0.89999 | 0.79039 | 0.10416 | **0.001** | **0.001** | 0.89999 | 0.27973 | **0.001** | **0.001** | 0.80937 | **0.0018** | **0.001** | 0.12789 | **0.001** | 0.13467 |
| **PabPTAL3 MA_73113g0010** | Ser595Phe | 0.89999 | 0.68176 | **0.0082** | **0.0059** | **0.001** | 0.64761 | **0.0086** | **0.0072** | **0.001** | 0.4482 | 0.61754 | **0.0015** | 0.89999 | 0.30079 | **0.0165** |
| | Leu601Phe | 0.89999 | 0.43629 | **0.0123** | **0.0024** | **0.001** | 0.47522 | **0.018** | **0.0049** | **0.001** | 0.74936 | 0.76679 | **0.0369** | 0.89999 | 0.54461 | 0.17789 |
