## Supplementary file4 for "In *silico* characterisation of PAL homologs and metabolomic profiling of shade response indicates potential presence of PTALs and Tyr as a probable precursor of lignin biosynthesis in conifers"

**Figure S1** Norway Spruce, LCMS data: PCA shows separation between all groups – North_Shade, North_Sun, South_Shade and South_Sun; targeted + untargeted metabolites, K = 799.

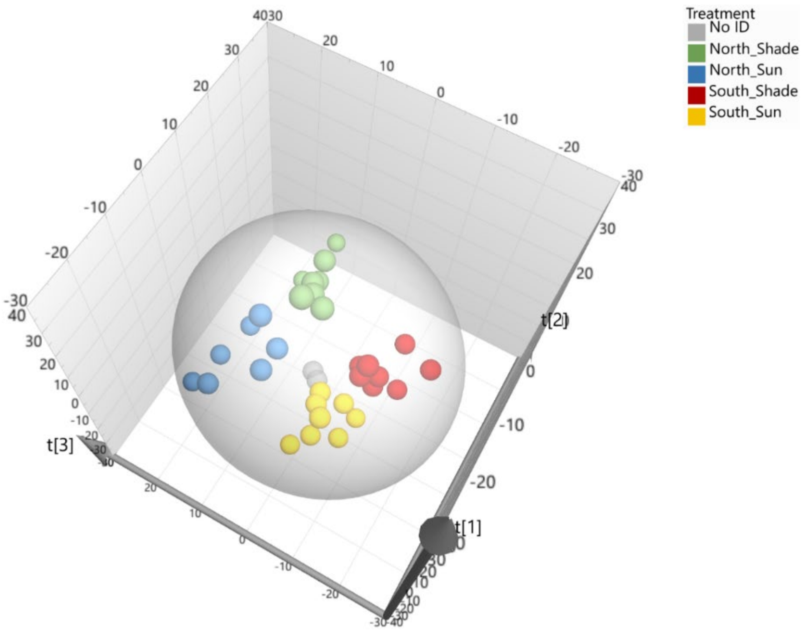


R2X[1] = 0.248; R2X[2] = 0.125; R2X[3] = 0.112; Ellipse: Hotelling's T2 (95%)

**Figure S2** Norway Spruce GCMS data: PCA shows separation between all groups – North_Shade, North_Sun, South_Shade and South_Sun; K = 69.

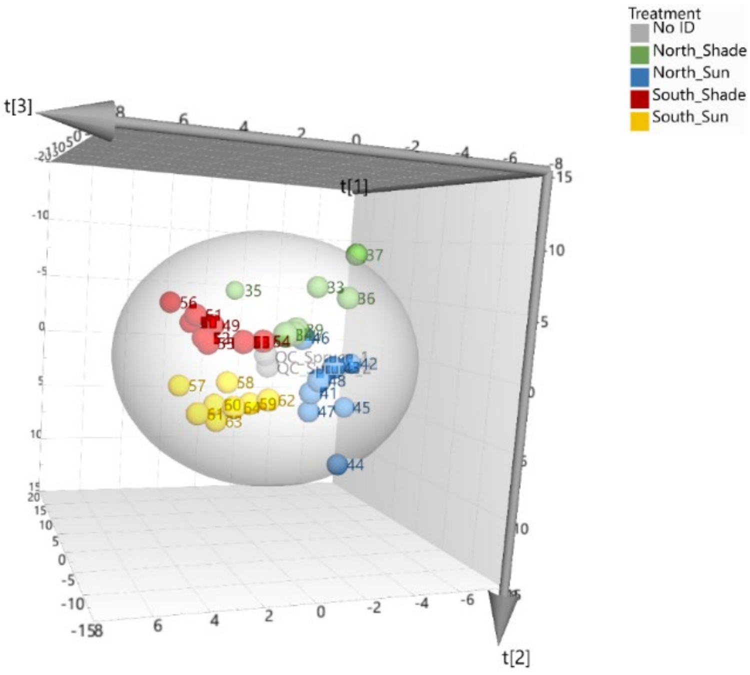


R2X[1] = 0.405; R2X[2] = 0.209; R2X[3] = 0.0859; Ellipse: Hotelling's T2 (95%)

**Figure S3** Scots pine LCMS data: PCA shows separation between all groups – North_Shade, North_Sun, South_Shade and South_Sun; targeted + untargeted metabolites, K = 781.

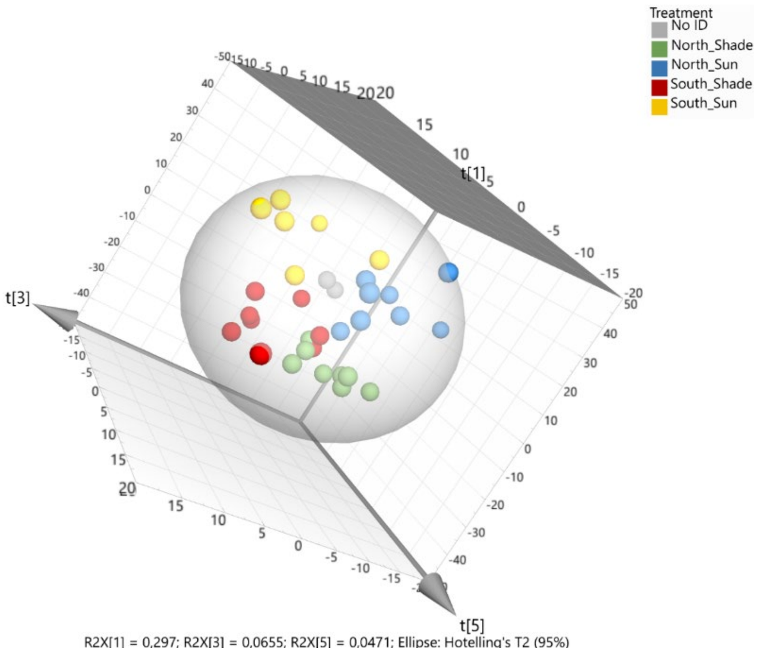


**Figure S4** Scots pine GCMS data: PCA shows clear separation between Shade and Sun conditions, but the Northern and Southern samples are not completely separated; K = 68.

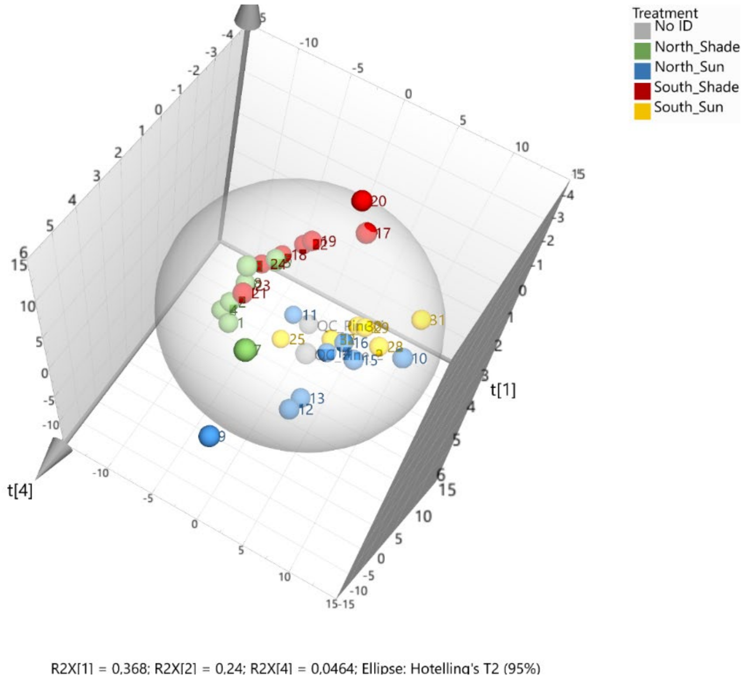

**Figure S5** Norway spruce samples, LCMS data: OPLS-DA shows differences between Shade and Sun conditions in both populations; targeted + untargeted metabolites, K = 799.

(a) Northern samples: R2Y = 0.97 Q2Y = 0.78
(b) Southern samples: R2Y = 0.98, Q2Y = 0.84

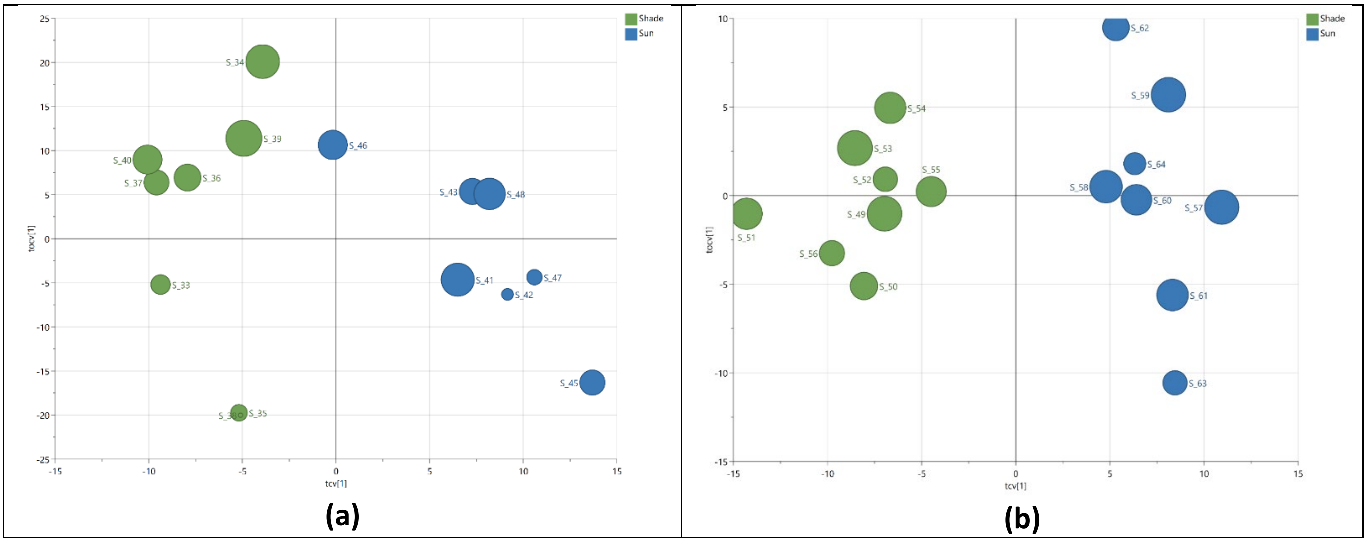


**Figure S6** Norway spruce samples, GCMS data: OPLS-DA shows differences between Shade and Sun conditions in both populations; K = 69.

(a) Northern samples: Clear separation between Shade and Sun conditions for all samples except S46. R2Y = 0.79 Q2Y = 0.58
(b) Southern samples: R2Y = 0.97, Q2Y = 0.94

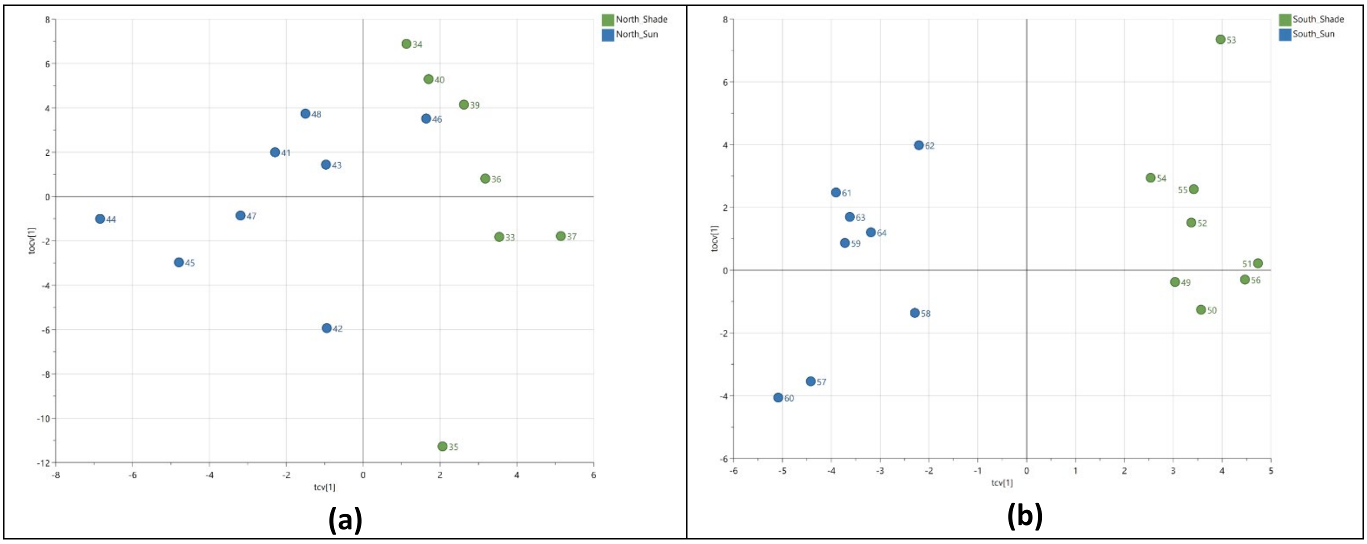

**Figure S7** Scots pine samples, LCMS data: OPLS-DA shows differences between Shade and Sun conditions in both populations; targeted + untargeted metabolites, K = 781.

(a) Northern samples: R2Y = 0.97, Q2Y = 0.87

(b) Southern samples: R2Y = 0.98, Q2Y = 0.92

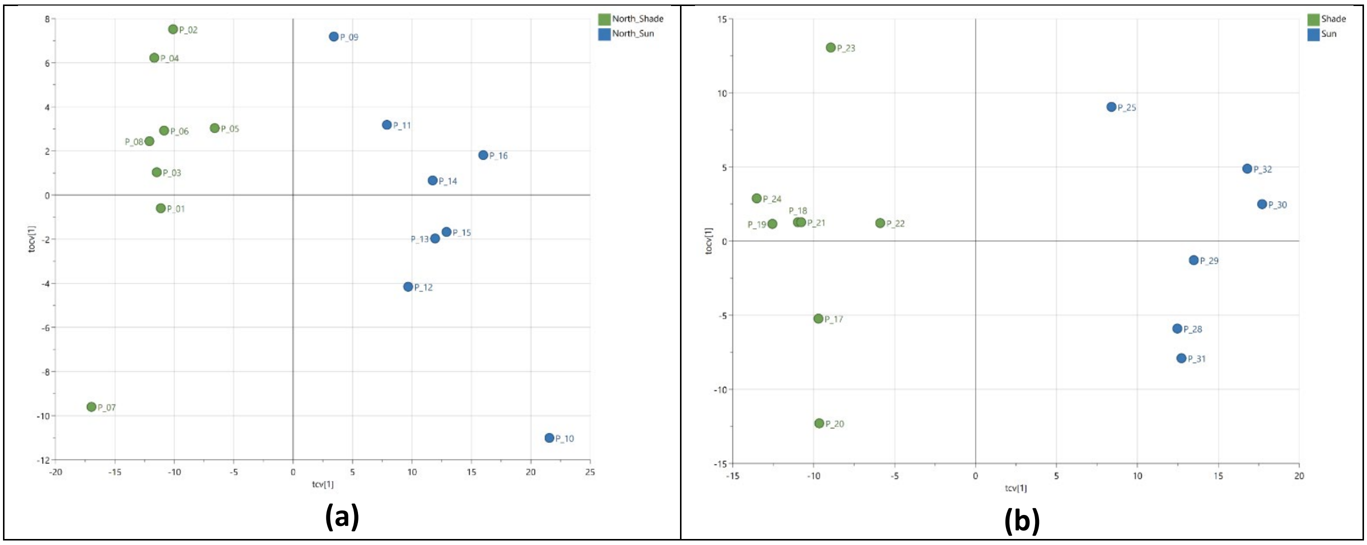


**Figure S8** Scots pine samples, GCMS data: OPLS-DA shows differences between Shade and Sun conditions in both populations; K = 68.

(a) Northern samples: R2Y = 0.93 Q2Y = 0.87

(b) Southern samples: R2Y = 0.96, Q2Y = 0.93

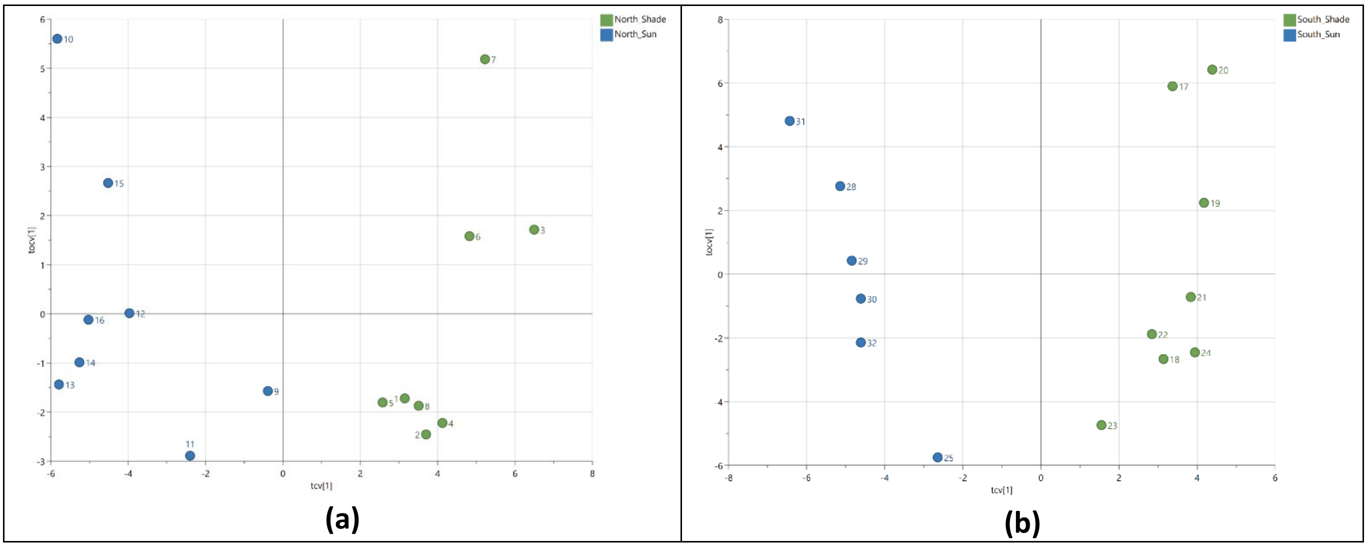

**Figure S9** Norway Spruce LCMS data: OPLS-DA, South vs North models; targeted + untargeted metabolites, K = 799.

- Metabolite loadings for Northern ecotype plotted against Southern ecotype.
- Shows similarity between models and up/down regulation in Sun & Shade conditions.
- Regression line, R2 = 0.44 – There is a slight similarity in metabolic response due to light condition for the two ecotypes.

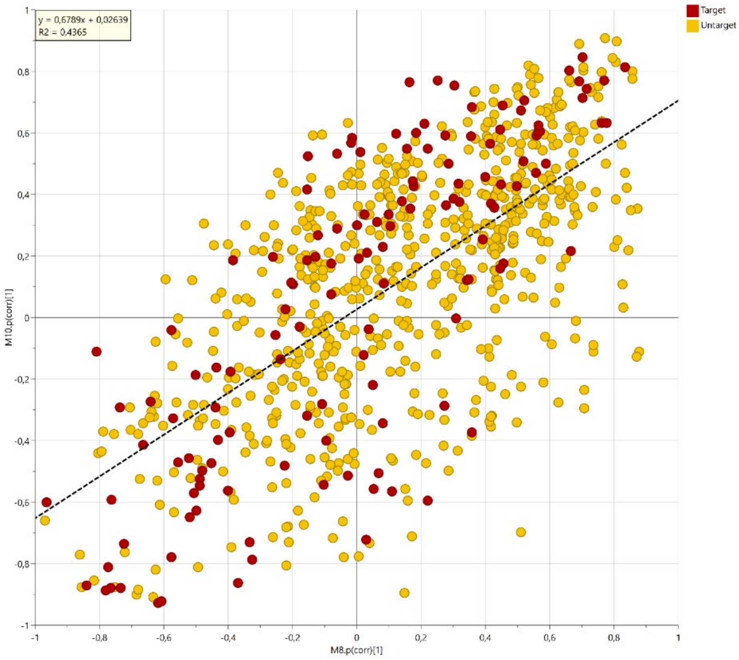


**Figure S10** Norway Spruce GCMS data: OPLS-DA, South vs North models; K = 69.

- Metabolite loadings for Northern ecotypes plotted against Southern ecotypes.
- Shows similarity between models and up/down regulation in Sun & Shade conditions.
- Regression lines, R2 = 0.75 – High similarity in metabolic response for Sun and Shade conditions for the two ecotypes.

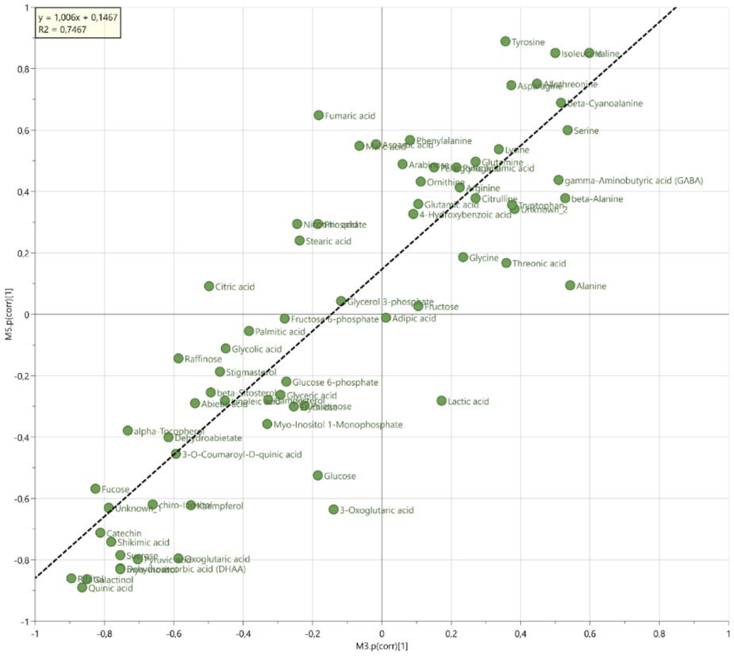

**Figure S11** Scots pine LCMS data: OPLS-DA, South vs North models; targeted + untargeted metabolites, K = 781.

- Metabolites for Northern ecotype plotted against Southern ecotype.
- Shows similarity between ecotype models and up/down regulation in Sun & Shade conditions.
- Regression line, R2 = 0.73 – Pine exhibits a more similar metabolic response in both light conditions compared to spruce.

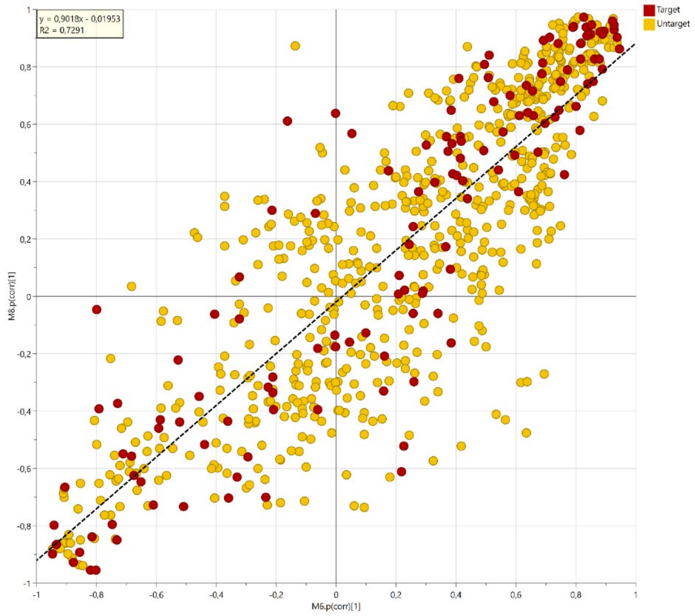


**Figure S12** Scots pine GCMS data: OPLS-DA, South vs North models; K = 68.

- Metabolite loadings for Northern ecotypes plotted against Southern ecotypes.
- Shows similarity between models and up/down regulation in Sun & Shade conditions.

Regression lines, R2 = 0.86 – High similarity in metabolic response for Sun and Shade conditions for the two ecotypes.

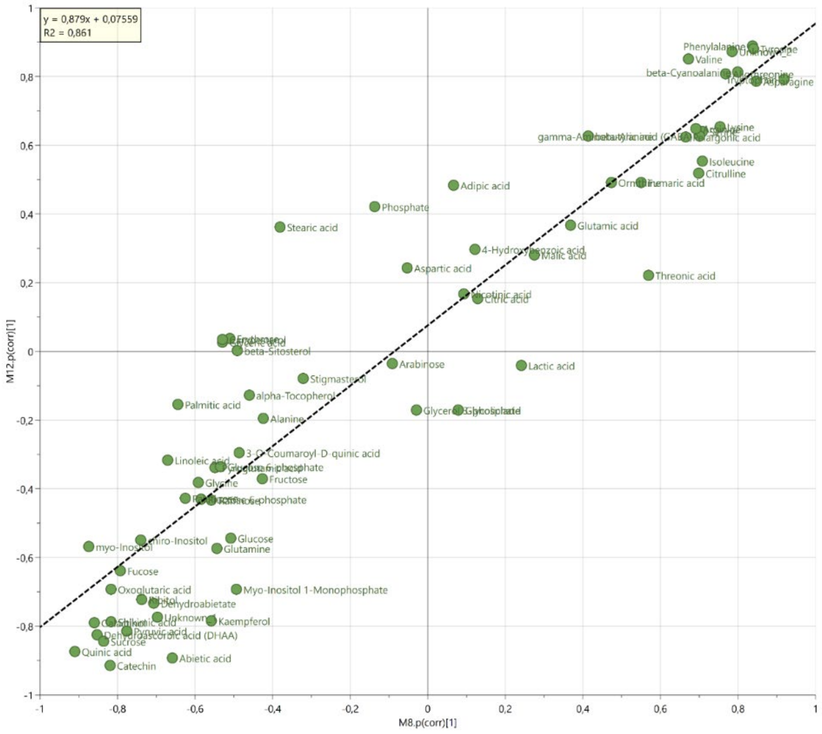

**Figure S13** Overview of Pathway Analysis: Northern Norway spruce, Sun vs Shade

The x axis shows pathway impact scores that summarize normalized topology measures of those perturbed metabolites in each pathway. The y axis shows −log10(P) values of the enrichment analysis results. The sizes of the data points are correlated with their x values, and the colour gradients correspond to their y values.

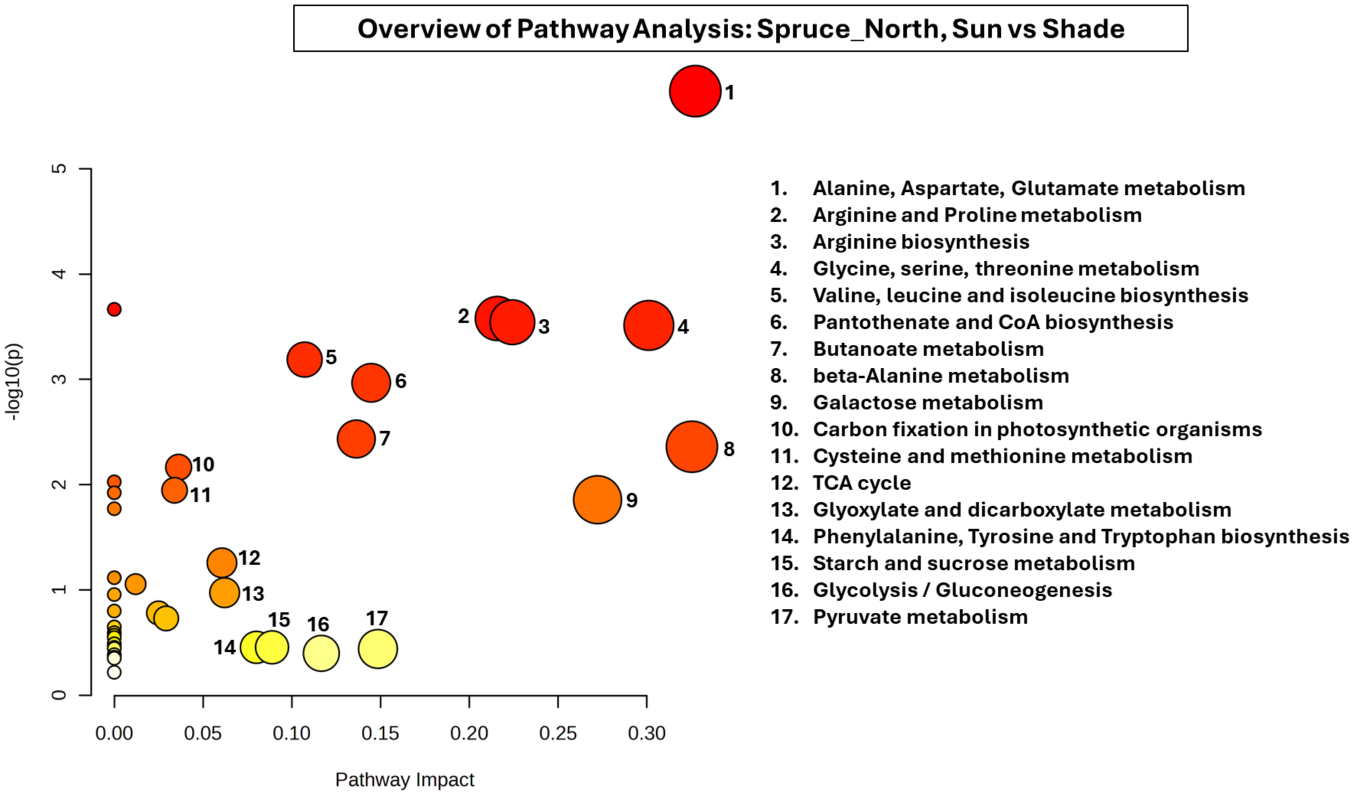

**Figure S14** Overview of Pathway Analysis: Southern Norway spruce, Sun vs Shade

The x axis shows pathway impact scores that summarize normalized topology measures of those perturbed metabolites in each pathway. The y axis shows −log10(P) values of the enrichment analysis results. The sizes of the data points are correlated with their x values, and the colour gradients correspond to their y values.

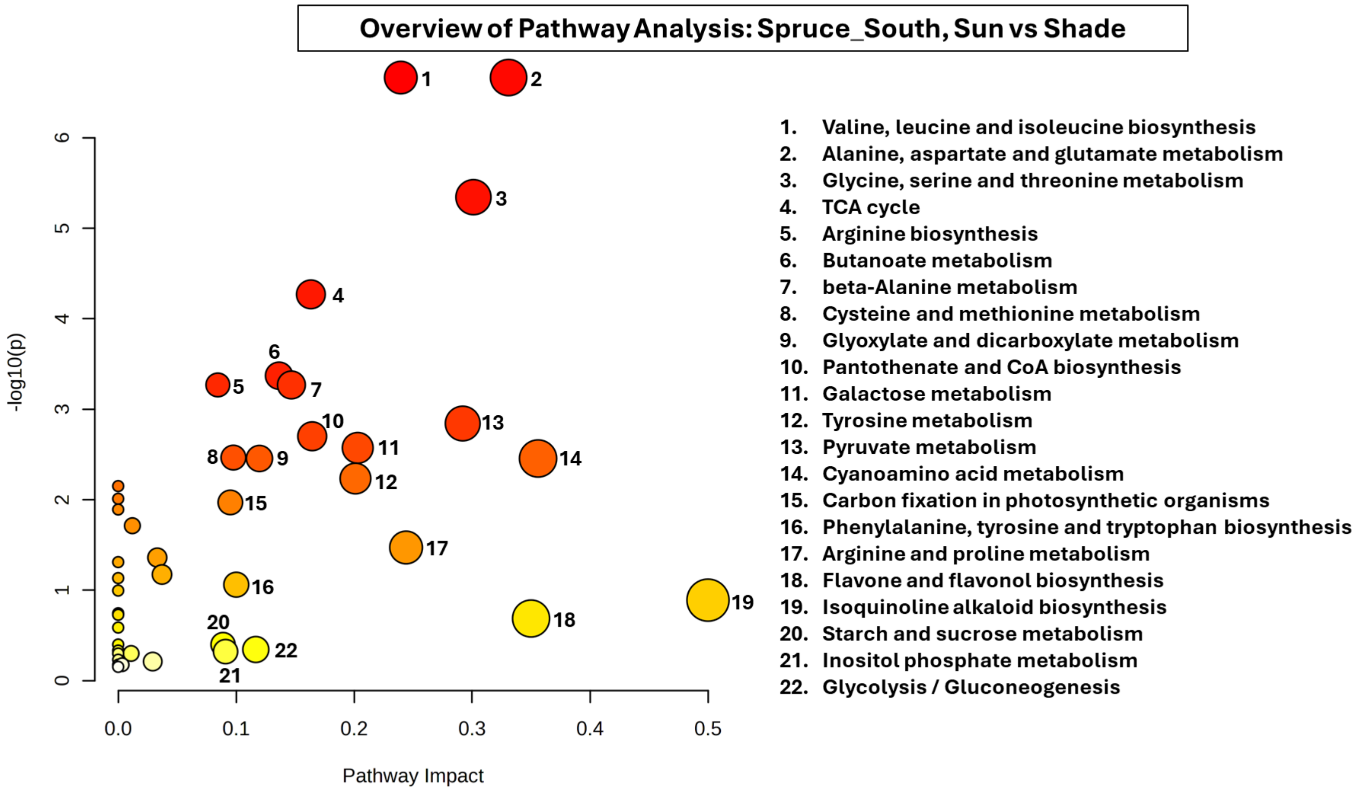

**Figure S15** Overview of Pathway Analysis: Northern Scots pine, Sun vs Shade

The x axis shows pathway impact scores that summarize normalized topology measures of those perturbed metabolites in each pathway. The y axis shows −log10(P) values of the enrichment analysis results. The sizes of the data points are correlated with their x values, and the colour gradients correspond to their y values.

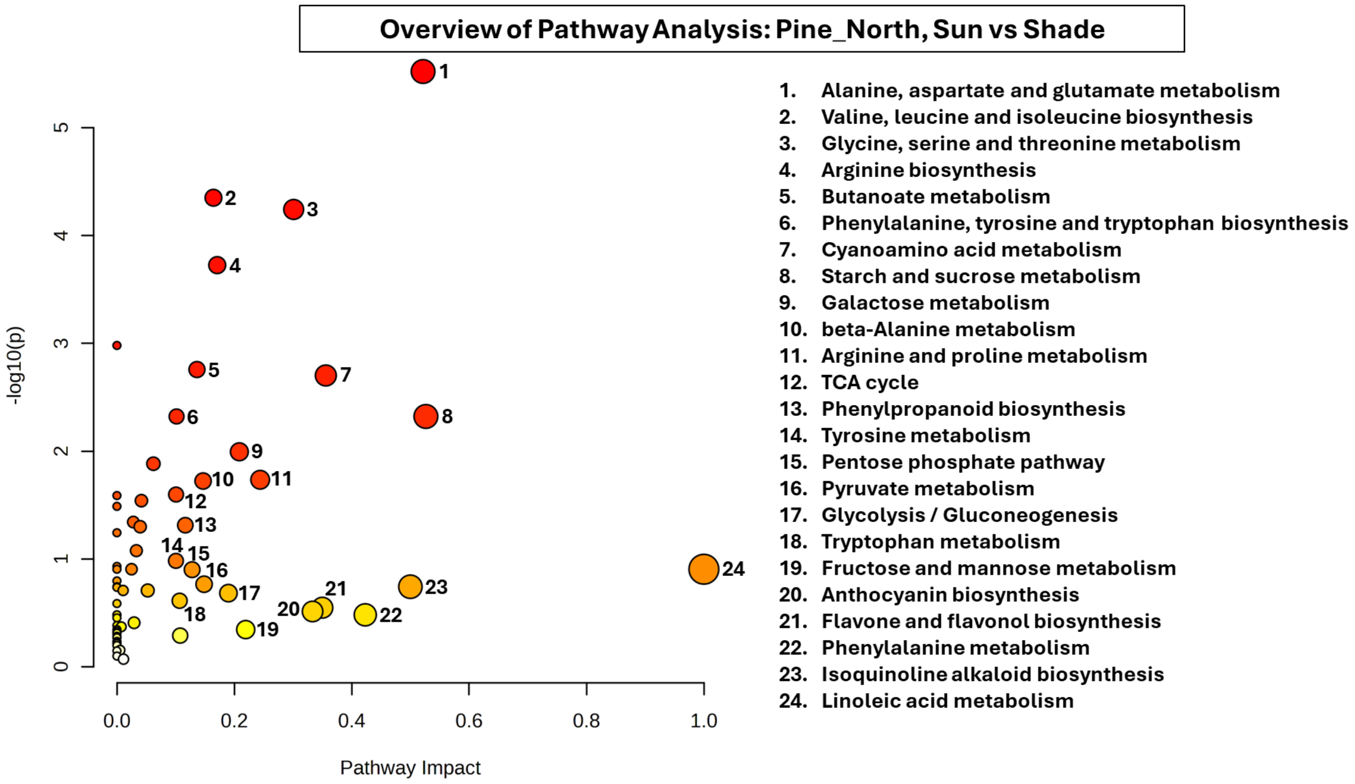

**Figure S16** Overview of Pathway Analysis: Southern Scots pine, Sun vs Shade

The x axis shows pathway impact scores that summarize normalized topology measures of those perturbed metabolites in each pathway. The y axis shows −log10(P) values of the enrichment analysis results. The sizes of the data points are correlated with their x values, and the colour gradients correspond to their y values.

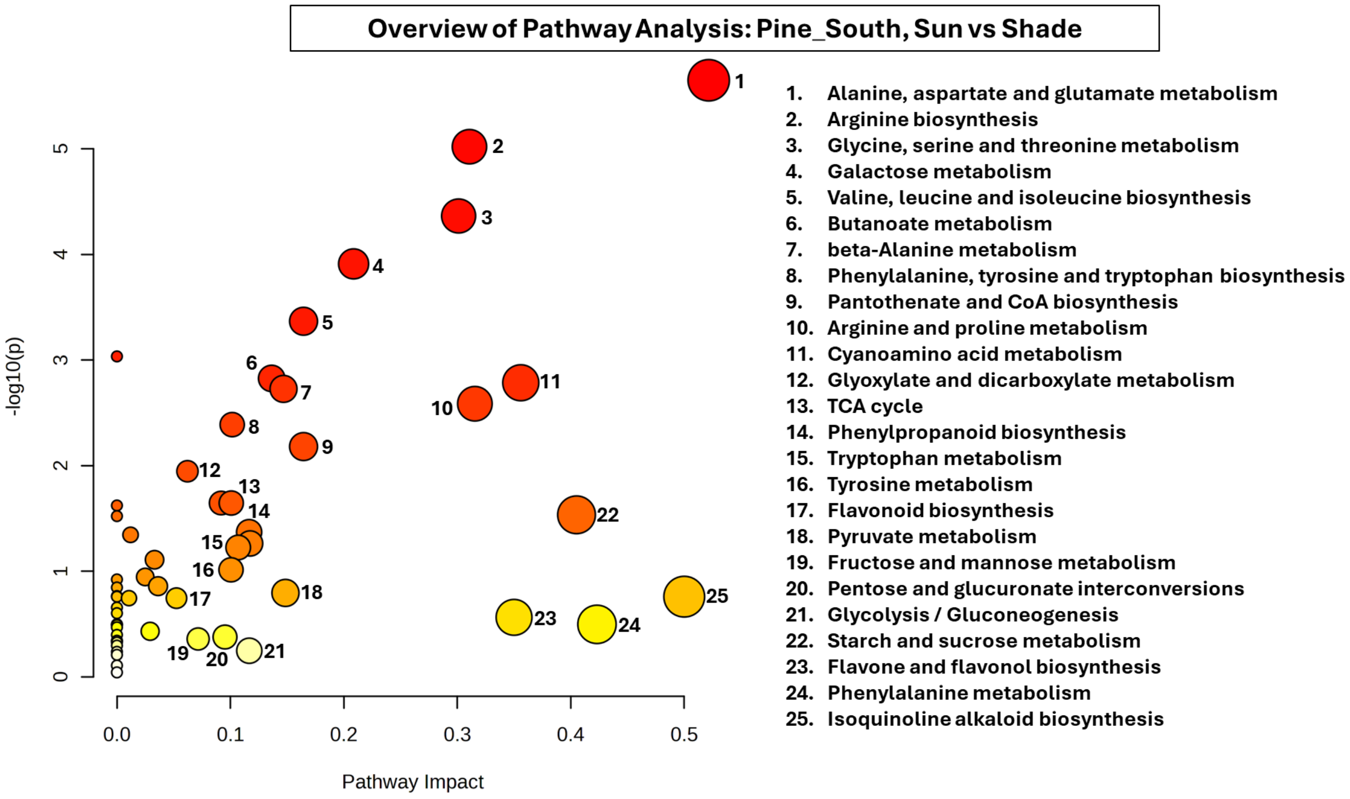

**Figure S17 Multiple sequence alignment of PALs and PTALs from plant model systems and putative PALs and PTALs from conifer species showing conserved amino acid residues (CLUSTAL O, 1.2.4). Colour codes – red: catalytically essential tyrosine; yellow: MIO region; green: asparagine and tyrosine stabilizing the MIO group; blue: arginine responsible for binding the carboxylic group of substrate. The conserved GTITASGDLVPLSYIA motif contains MIO region, marked in bold. Residues involved in substrate specificity for phenylalanine/tyrosine are marked with boxes (F/H, S/A, V/L, L/I, E/D).**

```
PtaPTAL3      ------------------------------------------------------------   0
PmePTAL4      ------------------------------------------------------------   0
PtaPTAL2      -----------------------------------------------MPE-------AAS   6
PmePTAL3      ------------------------------------------------------------   0
PabPTAL3      -------------------------------------------MVEVMAE-------AAP   10
PtaPAL4       ------------------------------------------------------------   0
PtaPAL3       ------------------------------------------------------------   0
PtaPAL1       ------------------------------------------------------------   0
PtaPAL2       ------------------------------------------------------------   0
PabPAL1       ------------------------------------------------------------   0
PmePAL1       ------------------------------------------------------------   0
OsPTAL        ------------------------------------------------------------   0
ZmPTAL        ------------------------------------------------------------   0
BdPTAL        ------------------------------------------------------------   0
BoPTAL        ------------------------------------------------------------   0
BoPAL         ------------------------------------------------------MECENG   6
OsPAL         ------------------------------------------------------MECENG   6
BdPAL         ------------------------------------------------------MECENG   6
ZmPAL         ------------------------------------------------------MESEAG   6
PcPAL         ------------------------------------------------------MENGNG   6
AtPAL         --------------------------------------------MEINGA---HKSNGGG   13
PtPAL         --------------------------------------------METITK----------   6
PabPAL2       --------------------------------------------MSSQGEEISGMENTNA   16
PsiPAL        --------------------------------------------MSSQGEEISRLENTNA   16
PmePAL2       ---------MLSLHYLAATSFLSAFSLRYCIAVPRPSASIHQIQEEIMSSTLGKENSVNG   51
PsyPAL        ------------------------------------------------------------   0
PsiPTAL       -----------------------------------------MVAA---AAEM---TQTNE   13
PabPTAL1      -----------------------------------------MVAA---AAEM---TQTNE   13
PmePTAL1      VSNKHSSFQLPCLALLCSAQFLDCL-VHFALQDHK------NMVA---GAEI---MQTNE   47
PsyPTAL       ------------------------------------------MVA---AAGM---TQASE   12
PtaPTAL1      ------------------------------------------MVA---AAEI---TQANE   12
PmePTAL2      ---------------------MVAG-VDMAVQANG------NGN-----------GNGNG   21
PabPTAL2      ---------------------MVAG-IDLAVQANG------NQNGDLAVQVN---GNQNG   29


PtaPTAL3      ------------MEFCQLNS--------------------NSCITARSRSCHVEDIILLI   28
PmePTAL4      ------------MECSQSKG--------------------KCC--KSAHSCHVKDIILLI   26
PtaPTAL2      TVMI--ASIV--SSLAIKDDINI----Q-GCHEYDHDSLNWSRVADAMKGSHANEVKNMI   57
PmePTAL3      --VM--GSMV--SLLAIKNDINM----E-GVHAYDHDRLNWSRAADAMKGSHGKEVKNMI   49
PabPTAL3      TVVM--GSM-------VKDDINM----E-SAHEYDHDGLNWSRAADAMKGSHAKEVRNMI   56
PtaPAL4       ---MASQEFTGLMNLCAGN-----------------DPLNWASVAESMKGSHFEEVKRMV   40
PtaPAL3       ---MAPQEFTGEVKFCAGND---------RV-SSLHDPLNWEAAAEPMKGSHFDEVKRMV   47
PtaPAL1       ---MAPQGITGEVKFCAGNG---------RV-SSLHDPLNWEAAAESMKGSHFDEVKRMV   47
PtaPAL2       ---MAPQGITGEVKFCAGNG---------RV-SSLHDPLNWEAAAESMKGSHFDEVKRMV   47
PabPAL1       ---MAPQEFTGEVKFCAGNG---------GTA-SLHDPLNWVAAAESMKGSHFEEVKRMV   47
PmePAL1       ---MAPQEFNGEVKFCAGNG---------GVGSNLHDPLNWAAAAESMKGSHFEEVKRMV   48
OsPTAL        ---------------MAGNG---------P--INKEDPLNWGAAAAEMAGSHLDEVKRMV   34
ZmPTAL        ---------------MAGNG---------A--IVESDPLNWGAAAAELAGSHLDEVKRMV   34
BdPTAL        ---------------MAGNG---------A--ISEKDPLNWGAAAAELTGSHLDEVKRMV   34
BoPTAL        ---------------MAGNG---------P--IVKDDPLNWGAAAAELTGSHFDEVKRMV   34
BoPAL         QVA--SNG----NGLCMATP--------------RADPLNWGKAAEELMGSHLEEVKRMV   46
OsPAL         RVS--ANGM---SGLCVAAP--------------RADPLNWGKATEEMTGSHLDEVKRMV   47
BdPAL         L----VGSL-NGEGLCMSAP---------P--RAAADPLNWAKTAEELAGSHLEEVKKMV   50
ZmPAL         LLV--RSSL-NGEGLCMPAP--------------RADPLNWGKAAEGLSGSHLDEVKRMV   49
PcPAL         --ATTNGHV-NGNGM---DF---------C--MKTEDPLYWGIAAEAMTGSHLDEVKKMV   49
AtPAL         VDAMLCGGD-IK-TK---NM---------V--INAEDPLNWGAAAEQMKGSHLDEVKRMV   57
PtPAL         -----NGYQ-NG-SS---ES---------L--CTQRDPLSWGVAAEAMKGSHLDEVKRMV   45
PabPAL2       L--M------RPMELCRPPILPLPESFGGKSPDHVIIPTHWKKAAEAMQCSHYEEVRKMI   68
PsiPAL        L--M------RPMELCRPPTLPLPESFGGKSPDHVIIPTHWKKAAEAMQCSHYEEVRKMI   68
PmePAL2       F--K------RPMELCRPPTLPLPGTLGNKTPDHVIIPTHWKKAAEALQCSHYEEVRKMI   103
PsyPAL        ------------------------------------------------------------   0
PsiPTAL       VQQV------KSTGLC--------TGLSSSS----SDPLNWIRAAKAMEGSHFEEVKTMV   55
PabPTAL1      VQQV------KSTGLC--------TGFSSSS----SDPLNWIRAAKAMEGSHFEEVKTMV   55
PmePTAL1      -VQV------KSTGLC--------TSFGSST----SDPLNWIRAAKAMEGSHFEEVKAMV   88
PsyPTAL       -VQV------KSTGLC--------TDFGSSG----SDPLNWVRAAKAMEGSHFEEVKAMV   53
PtaPTAL1      -VQV------KSTGLC--------TDFGSSG----SDPLNWVRAAKAMEGSHFEEVKAMV   53
PmePTAL2      FHHV------NAVDLC--------IQ---NG----PDPLNWGQAAKELQGSHFQEVKLMV   60
PabPTAL2      FHQV------HSVDLC--------IQ---NG----PDPLNWGRAAKALQGSHFEEVKLMV   68
```

```
PtaPTAL3        KTFNETNCINVDGSRITVAHVTALGRR---PQVKVTLDNHGGNCRERVERCS--------  77
PmePTAL4        NTFNEIQCINVDGSHITVAHVTALGRR---PQVKVALDDHGGHCRDNVERCS--------  75
PtaPTAL2        EEFQGIKVVSLTGFDLNIAQVTALARR---PDVKVVLDELN--AKQRVDASS--------  104
PmePTAL3        EEFQGIKVVSLTGSNLSIAQVTALARR---PDVKVVLDKLN--CKDRVDASS--------  96
PabPTAL3        EEFQGIKVVSLTGSSLSIAQVAALARR---PDVKVVLDELN--SKDGVDASS--------  103
PtaPAL4         EEFRA-PVVRLQGSGLTIAQVAAVARR--LGSVRVEFDT-G--ARARVEESS--------  86
PtaPAL3         EEFRA-PVVRLQGSGLTIAQVAAVARR--MGSVRVELET-G--AKARVDESS--------  93
PtaPAL1         EEFRA-PVVRLQGSGLTIAQVAAVARR--MGSVRVELET-G--AKARVDESS--------  93
PtaPAL2         EEFRA-PVVRLQGSGLTIAQVAAVARR--MGSVRVELET-G--AKARVDESS--------  93
PabPAL1         EEFRA-PVVKLQGSDLTIAQVAAVARR--LGSVRVELET-G--AKARVDESS--------  93
PmePAL1         EEFRS-PVVKLQGSGLTIAQVAAVARK--LGSVRVELET-C--AKARVDQSS--------  94
OsPTAL          AQFRE-PLVKIQGATLRVGQVAAVAQAKDAARVAVELDE-E--ARPRVKASS--------  82
ZmPTAL          AQARQ-PVVKIEGSTLRVGQVAAVASAKDASGVAVELDE-E--ARPRVKASS--------  82
BdPTAL          AQFRE-PVVKIEGASLRVGQVAAVAQAKDAAGVSVELDE-E--ARPRVKASS--------  82
BoPTAL          AQFRE-PVIKIEGASLRVGQVAAVAQAKDVSGVAVELDE-E--ARPRVKASS--------  82
BoPAL           AEYRQ-PVVKIEGASLRIAQVAAVA--AGAGEAKVQLDD-S--ARGRVKESS--------  92
OsPAL           AEYRQ-PLVKIEGASLRIAQVAAVA---AAGEARVELDE-S--ARERVKASS--------  92
BdPAL           AQFRM-PLVKIEGATLGIAQVAAVA--AGAGEARVELDE-S--ARGRVKESS--------  96
ZmPAL           AEFRD-PLVKIQGASLSVAQVAAVAVGAGGGEARVELDE-S--ARERVRASS--------  97
PcPAL           AEYRK-PVVKLGGETLTISQVAAISAR-DGSGVTVELSE-A--ARAGVKASS--------  96
AtPAL           AEFRK-PVVNLGGETLTIGQVAAISTI--GNSVKVELSE-T--ARAGVNASS--------  103
PtPAL           AEYRK-PVVNLAGQTLTIAQVASIAGH-DASNVKVELSE-S--ARPRVKASS--------  92
PabPAL2         GQFKATHKVVLRGTTLTVAEVTAVTRR---AEVKVELDEVS--AKQRVERSY--------  115
PsiPAL          GQFKATHKVVLRGTTLTVAEVTAVTRR---AEVKVELDEVS--AKQRVERSY--------  115
PmePAL2         AQFNATHKVVLRGTTLTVAEVTAVTRK---ATVKVELDEAS--AKERVQESY--------  150
PsyPAL          ------------------------------------------------------------  0
PsiPTAL         DSYFESKEIFIEGKTLTIADVTAVARR---SQVKVKLDAAA--AKSRVEESS--------  102
PabPTAL1        DSYFESKEIFIEGKTLTIADVTAVARR---SQVKVKLDAAA--AKSRVEESSNWVLTQMT  110
PmePTAL1        NSYLGSKEISIEGKSLTISDVAAVARR---SQVKVKLDAAA--AKSRVEESS--------  135
PsyPTAL         DSYLGVKEIFIEGKSLTISDVAAVARR---SQVKVKLDAEA--AKSRVEESS--------  100
PtaPTAL1        DSYFGAKEISIEGKSLTISDVAAVARR---SQVKVKLDAAA--AKSRVEESS--------  100
PmePTAL2        ESYFGSQEVSIEGKTLTIADVAAVARR---PEAKVRLDAVS--AKSRVDESS--------  107
PabPTAL2        ESYFGSKEVSIEGKSLTIADVAAVARR---PEAKVRLDAVA--AKARVDESS--------  115

                                                             F/H

PtaPTAL3        ------LWVQEKAKGGADIYGVTTGFGACSSKRTNQLSELQEALIRCLLAGVFTGP--SS  129
PmePTAL4        ------LWVREKAKEGADIYGVTTGFGACSSKRANQLSVLQEALIRCLLAGVFTGGPASS  129
PtaPTAL2        ------EWVMNNINRGTDTYGITTGFGATSHRRTNQAVELQRELIRFLNAGVMGKRDS--  156
PmePTAL3        ------DWVMNNINKGTDTYGVTTGFGATSHRRTNQAVELQRELIRFLNAGVMGKGGS--  148
PabPTAL3        ------DWVMNNINKGTDTYGVTTGFGATSHRRTNQAVELQKELIRFLNAGVMGKGDS--  155
PtaPAL4         ------NWVMDSIANGKAIYGVTTGFGASSHRRTSHGEALQKEMARFLNAGIFGGCGD--  138
PtaPAL3         ------NWVMDSMANGTDSYGVTTGFGATSHRRTRQGEALQKELIRFLNAGIFGACGD--  145
PtaPAL1         ------NWVMDSMANGTDSYGVTTGFGATSHRRTRQGEALQKELIRFLNAGIFGACGD--  145
PtaPAL2         ------NWVMDSMANGTDSYGVTTGFGATSHRRTRQGEALQKELIRFLNAGIFGACGD--  145
PabPAL1         ------NWVMDSMANGTDSYGVTTGFGATSHRRTRQGEALQKELIRFLNAGIFGGCGD--  145
PmePAL1         ------NWVMDSMANGTDSYGVTTGFGATSHRRTRQGEALQKELIRFLNAGIFGGCGD--  146
OsPTAL          ------EWILTCIAHGGDIYGVTTGFGGTSHRRTKDGPALQVELLRHLNAGIFGTGSD--  134
ZmPTAL          ------EWILDCIAHGGDIYGVTTGFGGTSHRRTKDGPALQVELLRHLNAGIFGTGSD--  134
BdPTAL          ------EWILSCLAAGGDIYGVTTGFGGTSHRRTKDGPALQVELLRHLNAGIFGTGSD--  134
BoPTAL          ------EWILNCLAHGGDIYGVTTGFGGTSHRRTKDGPALQVELLRHLNAGIFGTGTD--  134
BoPAL           ------DWVMNSMMNGTDSYGVTTGFGATSHRRTKEGGALQRELIRFLNAGAFGTGSD--  144
OsPAL           ------DWVMNSMMNGTDSYGVTTGFGATSHRRTKEGGALQRELIRFLNAGAFGTGTD--  144
BdPAL           ------DWVMNSMMNGTDSYGVTTGFGATSHRRTKEGGALQRELIRFLNAGAFGTGTD--  148
ZmPAL           ------DWVMGSMMNGTDSYGVTTGFGATSHRRTKEGGALQRELIRFLNAGAFGTGAD--  149
PcPAL           ------DWVMDSMNKGTDSYGVTTGFGATSHRRTKQGGALQKELIRFLNAGIFGNGSD--  148
AtPAL           ------DWVMESMNKGTDSYGVTTGFGATSHRRTKNGVALQKELIRFLNAGIFGSTKE--  155
PtPAL           ------DWVMDSMDKGTDSYGVTTGFGATSHRRTKQGGALQKELIRFLNAGIFGNGTE--  144
PabPAL2         ------QWVANNIAKGTDTYGVTTGFGATSHRRTDKAAELQKELIRFLNAGVVGKER---  166
PsiPAL          ------QWVANNIAKGTDTYGVTTGFGATSHRRTDKAAELQKELIRFLNAGVVGKER---  166
PmePAL2         ------QWVANNVARGTDTYGVTTGFGATSHRRTDKAEDLQKELIRFLNAGVVGKAR---  201
PsyPAL          ------QWVAKNVARGTDTYGVTTGFGATSHRRTDKAADLQKELIRFLNAGVVGREC---  51
PsiPTAL         ------NWVLTQMTKGTDTYGVTTGFGATSHRRTNQGAELQKELIRFLNAGVLGKCQD--  154
PabPTAL1        RVEESSNWVLTQMTKGTDTYGVTTGFGATSHRRTNQGAELQKELIRFLNAGVLGKCQD--  168
PmePTAL1        ------NWVLTQMTKGTDTYGVTTGFGATSHRRTNQGAELQKELIRFLNAGVLGKCQD--  187
PsyPTAL         ------NWVLTQMTKGTDTYGVTTGFGATSHRRTNQGAELQKELIRFLNAGVLGKCPE--  152
PtaPTAL1        ------NWVLTQMTKGTDTYGVTTGFGATSHRRTNQGAELQKELIRFLNAGVLGKCPE--  152
PmePTAL2        ------NWVLQNMLKGTDTYGVTTGFGATSHRRTNQGAELQKELIRFLNAGVLQ-AED--  158
PabPTAL2        ------NWVLQNMLKGTDTYGVTTGFGATSHRRTSQGAELQKELIRFLNSGVLT--EG--  165
                       *:      *   **:*****. * :*: .   **  : * * :* .
```

```
                                                                        MIO
PtaPTAL3     SPGELSPITARCAMFLRMSSFIYGCSGIRWEIMEALQQLINSHITPKCPLRGSVSASGDL  189
PmePTAL4     SPGELSPTTTRCAMFLRMNSFIYGCSGIRWEIMEALKKLINTHVTPKCPLRGSVSASGDL  189
PtaPTAL2     --NCLSVSATRAAMLVRTNTLMQGFSGIRWEILEAMQKLLDSHITPKLPLRGTITASGDL  214
PmePTAL3     --NCLSVSATRAAMLVRTNTLMQGFSGIRWEILEALQKLLDSHITPKLPLRGTITASGDL  206
PabPTAL3     --NCLSVSSTRAAMLVRTNTLMQGFSGIRWEILEAMQKLLDSHVTPKLPLRGTITASGDL  213
PtaPAL4      S-NTLPRDATRATMLVRTNTLLQGYSGIRWGILEAMTGLLNAGITPRLPLRGSITASGDL  197
PtaPAL3      S-NSLPRDTTRAAMLVRANTLLQGYSGIRWGILEAMSGLLNAGITPRLPLRGTITASGDL  204
PtaPAL1      S-NSLPRDTTRAAMLVRANTLLQGYSGIRWGILEAMSGLLNAGITPRLPLRGTITASGDL  204
PtaPAL2      S-NSLPRDTTRAAMLVRANTLLQGYSGIRWGILEAMSGLLNAGITPRLPLRGTITASGDL  204
PabPAL1      S-NSLPRDTTRAAMLVRANTLLQGYSGIRWGILEAMSGLLNAGITPRLPLRGTITASGDL  204
PmePAL1      S-NSLPRETTRAAMLVRANTLLQGYSGIRWEILEAMSGLLNAGITPRLPLRGTITASGDL  205
OsPTAL       G-HTLPSETVRAAMLVRINTLLQGYSGIRFEILEAITKLLNTGVTPCLPLRGTITASGDL  193
ZmPTAL       G-HTLPSEVTRAAMLVRINTLLQGYSGIRFEILEAITKLLNTGVSPCLPLRGTITASGDL  193
BdPTAL       G-HSLPAEVTRAAMLVRINTLLQGYSGIRFEILEAITKLLNTGVSPCLPLRGTITASGDL  193
BoPTAL       G-HTLPSEVTRAAMLVRINTLLQGYSGIRFEILEAITKLINTGVTPCLPLRGTITASGDL  193
BoPAL        G-HVLAAEATRAAMLVRINTLLQGYSGIRFEILEAIAKLLNANVTPCLPLRGTITASGDL  203
OsPAL        G-HVLPAEATRAAMLVRINTLLQGYSGIRFEILEAIAKLLNANVTPCLPLRGTITASGDL  203
BdPAL        G-HVLPAEATRAAMLVRINTLLQGYSGIRFEILEAITKLLNANVTPCLPLRGTITASGDL  207
ZmPAL        G-HVLPAEATRAAMLVRVNTLLQGYSGIRFEILEAIAKLLNANVTPCLPLRGTITASGDL  208
PcPAL        --NTLPHSATRAAMLVRINTLLQGYSGIRFEILEAITKFLNQNITPCLPLRGTITASGDL  206
AtPAL        TSHTLPHSATRAAMLVRINTLLQGFSGIRFEILEAITSFLNNNITPSLPLRGTITASGDL  215
PtPAL        TCHTLPHSATRAAMLVRINTLLQGYSGIRFEILEAITKLLNNNITPCLPLRGTITASGDL  204
PabPAL2      --LCLPAEYTKAAMLVRTNTLMQGYSGIRWEILDAMRKLMDCNITPKLPLRGTIE---AL  221
PsiPAL       --LCLPAEYTKAAMLVRTNTLMQGYSGIRWEILDAMRKLMDCNITPKLPLRGTITASGDL  224
PmePAL2      --LCLSAEYTKAAMLVRTNTLMQGYSGIRWEILDAMRKLMDCNIIPKLPLRGTITASGDL  259
PsyPAL       --LCLPAEYTKAAMLVRTNTLMQGYSGIRWEILDALRKLMDCNITPKLPLRGTITASGDL  109
PsiPTAL      --NVLSEDTTRAAMLVRTNTLLQGYSGIRWDILETVEKLLNAGLTPKLPLRGTITASGDL  212
PabPTAL1     --NVLSEDTTRAAMLVRTNTLLQGYSGIRWDILETVEKLLNAGLTPKLPLRGTITASGDL  226
PmePTAL1     --NILSEDTTRAAMLVRTNTLLQGYSGIRWDILETVEKLLNAGLTPKLPLRGTITASGDL  245
PsyPTAL      --NVLSEDTTRAAMLVRTNTLLQGYSGIRWDILETVEKLLNAGLTPKLPLRGTITASGDL  210
PtaPTAL1     --NVLSEDTTRAAMLVRTNTLLQGYSGIRWDILETVEKLLNAGLTPKLPLRGTITASGDL  210
PmePTAL2     --NVLPQATTRAAMLVRTNTLMQGYSGIRWEILETIQKLLNAGITPKLPLKGTITASGDL  216
PabPTAL2     --NVLPQATTRAAMLVRTNTLMQGYSGIRWDILETIQKLLNAGITPKLPLKGTITASGDL  223
                 *    .:.:*::* .::: * ****: *::::  :::  : *  **:*::     *
```

```
                                                                          A/S
PtaPTAL3     IPLAYIAGLLIGNPQVKARIGAHGEEQEVPAPEALMKAGLQ--PFKLQAKEGLALVNGTS  247
PmePTAL4     IPLAYIAGLLIGNPHVKARIGHHDGHEELSAPEALVKTGLQ--PFKLQAKEGLALVNGTS  247
PtaPTAL2     VPLSYIAGLLIARPNSVVVGI---DGNEMGAEEGLKLAGIDK-PFELNPKEGLALVNGTA  270
PmePTAL3     VPLSYIAGLLTARPNSVATRA---NGIEMGAEEALRMAGIDK-SFELNPKEGLALVNGTA  262
PabPTAL3     VPLSYIAGLLTARSNSVAIGV---NGNEMGPKEALILAGIDK-PFELNPKEGLALVNGTA  269
PtaPAL4      VPLSYIAGLLIGRPNARAVMA---DGTEVGAAEALAAAGVGNGPFVLRPKEGVALVNATA  254
PtaPAL3      VPLSYIAGLLTGRSNARAVTA---NGTELGAAEALAAAGVENGPFELRPKEGLALVNGTA  261
PtaPAL1      VPLSYIAGLLTGRSNARAVTA---NGTELGAAEALAAAGVENGPFELRPKEGLALVNGTA  261
PtaPAL2      VPLSYIAGLLTGRSNARAVTA---NGTELGAAEALAAAGVENGPFELRPKEGLALVNGTA  261
PabPAL1      VPLSYIAGLLTGRPNARAVTA---DGRELGAAEALAAAGVENGPFELRPKEGLALVNGTA  261
PmePAL1      VPLSYIAGLLTGRPNARAVTA---DGKELGSAEALAAAGVENGPFELRPKEGLALVNGTA  262
OsPTAL       VPLSYIAGLITGRPNAQAISP---DGRKVDAAEAFKLAGIEGGFFTLNPKEGLAIVNGTS  250
ZmPTAL       VPLSYIAGLITGRPNAQAVTV---DGRKVDAAEAFKIAGIEGGFFKLNPKEGLAIVNGTS  250
BdPTAL       VPLSYIAGLITGRPNAQATTA---DGRKVDAAEAFKVAGIEGGFFTLNPKEGLAIVNGTS  250
BoPTAL       VPLSYIAGLITGRPNAQAVAP---DGRKVDAAEAFKIAGIEGGFFKLNPKEGLAIVNGTS  250
BoPAL        VPLSYIAGLVTGRENSVAVAP---DGRKVNAAEAFKIAGIQGGFFELQPKEGLAMVNGTA  260
OsPAL        VPLSYIAGLVTGRENAVAVAP---DGSKVNAAEAFKIAGIQGGFFELQPKEGLAMVNGTA  260
BdPAL        VPLSYIAGLITGRQNSVAVAP---DGSKVSAAEAFKIAGIEHGFFELQPKEGLAMVNGTA  264
ZmPAL        VPLSYIAGLITGRQNSVAVDP---DGRKVGAAEAFKIAGIEHGFFELQPKEGLAMVNGTA  265
PcPAL        VPLSYIAGLLTGRPNSKAVGP---TGVILSPEEAFKLAGVEGGFFELQPKEGLALVNGTA  263
AtPAL        VPLSYIAGLLTGRPNSKATGP---NGEALTAEEAFKLAGISSGFFDLQPKEGLALVNGTA  272
PtPAL        VPLSYIAGLLTGRPNSKATGP---NGEVLDAVEAFKAAGIDSGFFELQPKEGLALVNGTA  261
PabPAL2      VPLSYIAGLLTARPNSKALAP---DGHVLDAMDALRKAGIPE-PFKLQPKEGLALVNGTG  277
PsiPAL       VPLSYIAGLLTARPNSKALAP---DGQVLDAMDALRKASIPE-PFKLQPKEGLALVNGTG  280
PmePAL2      VPLSYIAGLLTARPNSRALTP---DGHELDAMAALRKASIPE-PFKLQPKEGLALVNGTA  315
PsyPAL       VPLSYIAGLLTARPNSKALSP---DGHLLDAMEALRKAGILE-PFKLQPKEGLALVNGTA  165
PsiPTAL      VPLSYIAGLLTGRPNSRVRSR---DGTEMSGAEALKKVGLEK-PFELQPKEGLAIVNGTS  268
PabPTAL1     VPLSYIAGLLTGRPNSRVRSR---DGTEMSGAEALQKIGLEK-PFELQPKEGLAIVNGTS  282
PmePTAL1     VPLSYIAGLLTGRPNSKVRSR---DGTEMSGAEALKKVGVEK-PFELAPKEGLAIVNGTS  301
PsyPTAL      VPLSYIAGLLTGRPNSRVRSR---DGIEMSGAEALKKVGLEK-PFELQPKEGLAIVNGTS  266
PtaPTAL1     VPLSYIAGLLTGRPNSRVRSR---DGIEMSGAEALKKVGLEK-PFELQPKEGLAIVNGTS  266
PmePTAL2     VPLSYIAGLLTGRPNSKARCR---DGKEIGALEALQQVGVEK-PFELQPKEGLAIVNGTS  272
PabPTAL2     VPLSYIAGFLTGRPNSKGRCR---DGKELGALEALQQIGVEK-PFELQPKEGLAIVNGTS  279
             :**:****:: .. :              :    .:   .:    * *  ***:*:**.*.
```

```
PtaPTAL3      FATALAATVMYDANVLLLLVEMLCGMFCEVIFGREEFAHPLIHEMKPHPGQRESAALLEW   307
PmePTAL4      FATALASTVMYDANVLLLLVETLCGMFCEVIFGREEFAHPLIHTMKPHVGQIQSAALLEW   307
PtaPTAL2      VGSAVACTVCYDANVLAVFAEIGSAFFCEVMQGKPEFTDPLTHRLKHHPGQMEAGAVMEW   330
PmePTAL3      VGAAVACTVCYDANVLAVFAEIGSAFFCEVMQGKPEFTDPLTHRLKHHPGQIEAGAVMEW   322
PabPTAL3      VGAAVACTVCYDANVLAVFAEIG---------------------------QMEAGAVMEW   302
PtaPAL4       VGSALAATVLFDANVVVLLSEVLSGLFCEVMQGDPGFTNHLIHRLKDHPGQIEAAAIMEH   314
PtaPAL3       VGSALAATVLFDANVVALLSEVLSAMFCEVMQGNPEFTDHLTHRLKHHPGQIEAAAIMEH   321
PtaPAL1       VGSALAATVLFDANVVALLSEVLSAMFCEVMQGNPEFTDHLTHRLKHHPGQIEAAAIMEH   321
PtaPAL2       VGSALAATVLFDANVVALLSEVLSAMFCEVMQGNPEFTDHLTHRLKHHPGQIEAAAIMEH   321
PabPAL1       VGSALAATVLFDANVVVLLSEVLSALFCEVMQGNPEFTDHLTHRLKHHPGQIEAAAIMEH   321
PmePAL1       VGSALAATVLFDANVVVLLSEVVSALFCEVMQGNPEFTDNLTHRLKHHPGQIEAAAIMEH   322
OsPTAL        VGSALAATVMFDANILAVLSEVLSAVFCEVMNGKPEYTDHLTHKLKHHPGSIDAAAIMEH   310
ZmPTAL        VGSALAATVMYDANVLAVLSEVLSAVFCEVMNGKPEYTDHLTHKLKHHPGSIEAAAIMEH   310
BdPTAL        VGSALAATVLFDCNVLAVLSEVLSAVFCEVMNGKPEFTDHLTHKLKHHPGSIEAAAIMEH   310
BoPTAL        VGSALAATVLYDCNVLAVLSEVLSAVFCEVMNGKPEYTDHLTHKLKHHPGSIEAAAIMEH   310
BoPAL         VGSGLASTVLFEANILAILAEVLSAVFCEVMNGKPEYTDHLTHKLKHHPGQIEAAAIMEH   320
OsPAL         VGSGLASTVLFEANILAILAEVLSAVFCEVMNGKPEYTDHLTHKLKHHPGQIEAAAIMEH   320
BdPAL         VGSGLASTVLFEANIQAILAEVLSAVFCEVMTGKPEFTDHLTHKLKHHPGQIEAAAIMEH   324
ZmPAL         VGSGLASTVLFEANVLAVLAEVISAVFCEVMTGKPEFTDHLTHKLKHHPGQIEAAAVMEH   325
PcPAL         VGSGMASMVLFEANILAVLAEVMSAIFAEVMQGKPEFTDHLTHKLKHHPGQIEAAAIMEH   323
AtPAL         VGSGMASMVLFETNVLSVLAEILSAVFAEVMSGKPEFTDHLTHRLKHHPGQIEAAAIMEH   332
PtPAL         VGSGLASMVLFETNVLAVLSELISAIFAEVMNGKPEFTDHLTHKLKHHPGQIEAAAIMEH   321
PabPAL2       VGSAVAASVCFDANVLVVLAEILSAFFCEVMQGKPEFVDPLTHQLKHHPGQIEAAAVMEY   337
PsiPAL        VGSAVAASVCFDANVLVVLAEILSAFFCEVMQGKPEFVDPLTHQLKHHPGQIEAAAVIEY   340
PmePAL2       VGSAVAASVCFDANVLGVLAEILSALFCEVMQGKPEFVDPLTHQLKHHPGQIEAAAVMEY   375
PsyPAL        VGSAVAASVCFDANVLGVLAEILSALFCEVMQGKPEFVDPLTHQLKHHPGQIEAAAVMEF   225
PsiPTAL       VGAALASIVCFDANVLALLSEVISALFCEVMNGKPEFTDPLTHKLKHHPGQMEAAAIMEY   328
PabPTAL1      VGAALASIVCFDANVLALLSEVISALFCEVMNGKPEFTDPLTHKLKHHPGQMEAAAIMEY   342
PmePTAL1      VGAALASIVCFDANVLALLSEVISAMFCEVMNGKPEFTDPLTHKLKHHPGQMEAAAIMEY   361
PsyPTAL       VGAALASIVCFDANVLALLSEVISAMFCEVMNGKPEFTDPLTHKLKHHPGQMEAAAIMEY   326
PtaPTAL1      VGAALASIVCFDANVLALLSEVISAMFCEVMNGKPEFTDPLTHKLKHHPGQMEAAAIMEY   326
PmePTAL2      VGAALASIVCFDANVICILAEVLSAMFCEVMLGKPEFTDPLTHKLKHHPAQMEAAAIMEY   332
PabPTAL2      VGAALASIVCFDANVICILAEVLSAMFCEVMLGKPEFTDPLTHRLKHHPAQMEAAAIMEY   339
              ..:.:*. * :: *:  :: *                             . ::.*::*

PtaPTAL3      LLRDSPFQELS-REYYSINSLKKPKQDRYALRSSPQWLAPLVQTIRE-------------   353
PmePTAL4      LLRDSPFQELS-REYYSINSLKKPKQDRYALRSSPQWLAPIVQTIRE-------------   353
PtaPTAL2      LLDGSSYSF---LKLAETDPLKKPKQDRYALRTSPQWLGPQIEVIRM-------------   374
PmePTAL3      ILDGSPYVN---PKLAETDPLKKPKQDRYALRTSPQWLGPQIEVIRM-------------   366
PabPTAL3      LLDGSSYSY---LKLVETDPLKKPKQDRYALRTSPQWLGPQIEVIRM-------------   346
PtaPAL4       LLDGSSYMKAAAAKNKEADPMSKPKKDRYALYTSPQWLGPQVEVIRA-------------   361
PtaPAL3       LLEGSSYMKAAAAKHHEADALSKPKQDRYALRTAPQWLGPQIEVIRA-------------   368
PtaPAL1       LLEGSSYMKAAAAKHHEADALSKPKQDRYALRTAPQWLGPQIEVIRA-------------   368
PtaPAL2       LLEGSSYMKAAAAKHHEADALSKPKQDRYALRTAPQWLGPQIEVIRA-------------   368
PabPAL1       LLDGSSYMKAAAAKHQEADALSKPKQDRYALRTAPQWLGPQVEVLRA-------------   368
PmePAL1       LLDGSSYMKAMAAKHQEADALSKPKQDRYALRTSPQWLGPQVEVIRA-------------   369
OsPTAL        ILAGSSFMSHA-KKVNEMDPLLKPKQDRYALRTSPQWLGPQIQVIRA-------------   356
ZmPTAL        ILDGSSFMKQA-KKLNELDPLLKPKQDRYALRTSPQWLGPQIEVIRA-------------   356
BdPTAL        ILAGSSFMSHA-KKVNEIDPQLKPKQDRYALRTSPQWLGPQIEVIRS-------------   356
BoPTAL        ILAGSSFMSHA-KKVNEMDPLLKPKQDRYALRTSPQWLGPQIEVIRA-------------   356
BoPAL         ILEGSSYMKLA-KKLGELDPLMKPKQDRYALRTSPQWLGPQIEVIRA-------------   366
OsPAL         ILEGSSYMKHA-KKLGELDPLMKPKQDRYALRTSPQWLGPQIEVIRA-------------   366
BdPAL         ILEGSSYMKEA-KKQGELDPLMKPKQDRYALRTSPQWLGPQIEVIRF-------------   370
ZmPAL         ILEGSSYMKLA-KRLGELDPLMKPKQDRYALRTSPQWLGPQIEVIRF-------------   371
PcPAL         ILDGSAYVKAA-QKLHEMDPLQKPKQDRYALRTSPQWLGPQIEVIRS-------------   369
AtPAL         ILDGSSYMKLA-QKLHEMDPLQKPKQDRYALRTSPQWLGPQIEVIRY-------------   378
PtPAL         ILDGSAYMKAA-KKLHEMDPLQKPKQDRYALRTSPQWLGPQIEVIRF-------------   367
PabPAL2       LLDGSDYVKEA-ARLHERDPLSKPKQDRYALRTSPQWLGPQIEVIRA-------------   383
PsiPAL        LLDGSDYVKEA-ARLHERDPLSKPKQDRYALRTSPQWLGPQIEVIRA-------------   386
PmePAL2       LLDGSDYVKEA-ARLHESDPLSKPKQDRYALRTSPQWLGPQIEVIRA-------------   421
PsyPAL        LLDGSDYVKEA-ARLHERDPLSKPKQDRYALRTSPQWLGPPIEVIRA-------------   271
PsiPTAL       VLDGSSYMKHA-AKLHEMNPLQKPKQDRYALRTSPQWLGPQAEVIRS-------------   374
PabPTAL1      VLDGSSYMKHA-AKLHEMNPLQKPKQDRYALRTSPQWLGPQVEVIRSATHMIESQVEVIR   401
PmePTAL1      VLDGSSYMKHA-AKLHEMNPLQKPKQDRYALRTSPQWLGPQVEVIRS-------------   407
PsyPTAL       VLDGSSYMKHA-AKLHEMNPLQKPKQDRYALRTSPQWLGPQVEIIRS-------------   372
PtaPTAL1      VLDGSSYMKHA-AKLHEMNPLQKPKQDRYALRTSPQWLGPQVEIIRS-------------   372
PmePTAL2      VLDGSSYMKNA-AKKHEMNPLQKPKQDRYALRTSPQWLGPQIEVIRA-------------   378
PabPTAL2      ---------------KHEMNPLQKPKQDRYALRTSPQWLGPQIEVIRA-------------   372
                              . :   ***:***** ::****.*  : :*
```

```
                                    L/V         I/L
PtaPTAL3     -ATATIETEINSANDNPIIDPVNGKALHGANFQGSAVGFYMDYVRVALAGLGKLIFAQFT  412
PmePTAL4     -ATATIETEINSANDNPIIDHVNGKALHGANFQGSAVGFYMDYVRVAVAGLGKLLFAQFT  412
PtaPTAL2     -STHAIAREINSVNDNPLIDVVRDRALHGGNFQGTPIGVYMDNIRLALAAIAKLIFAQFS  433
PmePTAL3     -ATHAIAREINSVNDNPLIDVSRDRALHGGNFQGTPIGVSMDNIRLALAAIAKLIFAQFS  425
PabPTAL3     -ATHAIAREINSVNDNPLIDVVRDRALHGGNFQGTSIGVSMDNIRLALAAIAKLIFAQFS  405
PtaPAL4      -STHMVQREINSVNDNPIIDAAGNKALNGGNFQGTPVGVAMDNVRLALAAVGKLIFAQMS  420
PtaPAL3      -STHMVQREINSVNDNPLIDAARNKALHGGNFQGTPVGVAMDNARLALAAVGKLMFAQML  427
PtaPAL1      -STHMVQREINSVNDNPLIDAARNKALHGGNFQGTPVGVAMDNARLALAAVGKLMFAQMS  427
PtaPAL2      -STHMVQREINSVNDNPLIDAARNKALHGGNFQGTPVGVAMDNARLALAAVGKLMFAQMS  427
PabPAL1      -STHMVQREINSVNDNPLIDAARNKALHGGNFQGTPVGVAMDNARLALAAVGKLMFAQMS  427
PmePAL1      -STHMVQREINSVNDNPLIDAARNKALHGGNFQGTPVGVAMDNARLALAAVGKLLFAQMS  428
OsPTAL       -ATKSIEREVNSVNDNPVIDVHRGKALHGGNFQGTPIGVSMDNARLAIANIGKLMFAQFS  415
ZmPTAL       -ATKSIEREVNSVNDNPVIDVHRGKALHGGNFQGTPIGVSMDNARLAIANIGKLMFAQFS  415
BdPTAL       -ATKSIEREVNSVNDNPVIDVHRGKALHGGNFQGTPIGVSMDNTRLAIANIGKLMFAQFS  415
BoPTAL       -ATKSIEREVNSVNDNPVIDVHRGKALHGGNFQGTPIGVSMDNTRLAIANIGKLMFAQFS  415
BoPAL        -ATKSIEREINSVNDNPLIDVSRGKALHGGNFQGTPIGVSMDNTRLAIAAVGKLMFAQFS  425
OsPAL        -ATKSIEREINSVNDNPLIDVSRGKALHGGNFQGTPIGVSMDNTRLAIAAIGKLMFAQFS  425
BdPAL        -ATKSIEREINSVNDNPLIDVSRGKAIHGGNFQGTPIGVSMDNTRLAIAAIGKLMFAQFS  429
ZmPAL        -ATKSIEREINSVNDNPLIDVSRGKALHGGNFQGTPIGVSMDNTRLALAAIGKLMFAQFS  430
PcPAL        -STKMIEREINSVNDNPLIDVSRNKAIHGGNFQGTPIGVSMDNTRLAIAAIGKLMFAQFS  428
AtPAL        -ATKSIEREINSVNDNPLIDVSRNKAIHGGNFQGTPIGVSMDNTRLAIAAIGKLMFAQFS  437
PtPAL        -STKSIEREINSVNDNPLIDVSRNKALHGGNFQGTPIGVSMDNVRLAIASIGKLLFAQFS  426
PabPAL2      -ATHSIEREINSVNDNPLIDLSRDMALHGGNFQGTPIGVSMDNMRISLAAIGKLIFAQFS  442
PsiPAL       -ATHSIEREINSVNDNPLIDVSRDMALHGGNFQGTPIGVSMDNMRISLAAIGKLIFAQFS  445
PmePAL2      -ATHSIEREINSVNDNPLIDVSRDMALHGGNFQGTPIGVSMDNMRIALAAIGKLLFAQFS  480
PsyPAL       -ATHSIEREINSVNDNPLIDVSRDMALHGGNFQGTPIGVSMDNMRISLAAVGKLLFAQFS  330
PsiPTAL      -ATHMIEREINSVNDNPVIDVARDLALHGGNFQGTPIGVSMDNLRLAISAIGRLMFAQFS  433
PabPTAL1     SATHMIEREINSVNDNPVIDVARDLALHGGNFQGTPIGVSMDNLRLAISAIGRLMFAQFS  461
PmePTAL1     -ATHMIEREINSVNDNPVIDVARDKALHGGNFQGTPIGVSMDNLRLAISAIGKLMFAQFS  466
PsyPTAL      -ATHMIEREINSVNDNPVIDVARDKALHGGNFQGTPIGVSMDNLRLSISAIGKLMFAQFS  431
PtaPTAL1     -ATHMIEREINSVNDNPVIDVARDKALHGGNFQGTPIGVSMDNLRLSISAIGKLMFAQFS  431
PmePTAL2     -ATHMIQREINSVNDNPVIDVARDKALHGGNFQGTPIGVSMDNLRLALSAIGKLMFAQFS  437
PabPTAL2     -ATHMIQREINSVNDNPVIDVARDKALHGGNFQGTPIGVSMDNLRLALAAIGKLMFAQFS  431
              :*  :  *:**.****:**   .  *::*.****: :*. **  *:::: :.:*:***:

                 D/E
PtaPTAL3     ELMIEFYSNGLPGNLSLGPDLSLDYGFKGIDIAMAAYSSELQFLANPVTTHVHSAEQHNQ  472
PmePTAL4     ELMIEFYSNGLPSNLALGPDLSLDYGFKGVDIAMAAYSSELQFLANPVTTHVHSAEQHNQ  472
PtaPTAL2     ELVNEYYNNGLPSNLSGGADPSLDYGLKGAEIAMASYTSEILFLANPVTNHVQSAEQHNQ  493
PmePTAL3     ELVNEYYNNGLPSNLSGGADPSLDYGLKGAEIAMASYTSEILFLANPVTSHVQSAEQHNQ  485
PabPTAL3     ELVNEYYNNGLPSNLSGGADPSLDYGLKGAEIAMASYTSEIMFLANPVTNHVQSAEQHNQ  465
PtaPAL4      ELVNDFYNKGLPSNLSGGPDPSLDYGFKGAEVAMASYTSELQFLANPVTTHVQSAEQHNQ  480
PtaPAL3      ELVNDFYNNGLPSNLSGGPDPSLDYGFKGAEIAMAAYTSELQFLANPVTTHVQSAEQHNQ  487
PtaPAL1      ELVNDFYNNGLPSNLSGGPDPSLDYGFKGAEIAMAAYTSELQFLANPVTTHVQSAEQHNQ  487
PtaPAL2      ELVNDFYNNGLPSNLSGGPDPSLDYGFKGAEIAMAAYTSELQFLANPVTTHVQSAEQHNQ  487
PabPAL1      ELVNDFYNNGLPSNLSGGPDPSLDYGFKGAEIAMAAYTSELQFLANPVTTHVQSAEQHNQ  487
PmePAL1      ELVNDFYNNGLPSNLSGGPDPSLDYGFKGAEIAMAAYTSELQFLANPVTTHVQSAEQHNQ  488
OsPTAL       ELVNEFYNNGLTSNLAGSRNPSLDYGFKGTEIAMASYSSELQYLANPITNHVQSAEQHNQ  475
ZmPTAL       ELVNEFYNNGLTSNLAGSRNPSLDYGFKGTEIAMASYCSELQYLGNPITNHVQSAEQHNQ  475
BdPTAL       ELVNEFYNNGLTSNLAGSRNPSLDYGFKGTEIAMASYCSELQYLANPVTNHVQSAEQHNQ  475
BoPTAL       ELVNEFYNNGLTSNLAGSRNPSLDYGFKGTEIAMASYCSELQYLANPITNHVQSAEQHNQ  475
BoPAL        ELVNDFYNNGLPSNLSGGRNPSLDYGFKGAEIAMASYCSELQFLGNPVTNHVQSAEQHNQ  485
OsPAL        ELVNDFYNNGLPSNLSGGRNPSLDYGFKGAEIAMASYCSELQFLGNPVTNHVQSAEQHNQ  485
BdPAL        ELVNDFYNNGLPSNLSGGRNPSLDYGFKGAEIAMASYCSELQFLGNPVTNHVQSAEQHNQ  489
ZmPAL        ELVNDYYNNGLPSNLSGGRNPSLDYGFKGAEIAMASYCSELQFLGNPVTNHVQSAEQHNQ  490
PcPAL        ELVNDFYNNGLPSNLSGGRNPSLDYGFKGAEIAMASYCSELQFLANPVTNHVQSAEQHNQ  488
AtPAL        ELVNDFYNNGLPSNLTASRNPSLDYGFKGAEIAMASYCSELQYLANPVTSHVQSAEQHNQ  497
PtPAL        ELVNDFYNNGLPSNLTASRNPSLDYGFKGAEIAMASYCSELQYLANPVTSHVQSAEQHNQ  486
PabPAL2      ELVCDYYNNGLPSNLSGGPNPSLDYGFKGAEIAMAAYTSELQYLANPVTTHVQSTDQHNQ  502
PsiPAL       ELVCDYYNNGLPSNLSGGPNPSLDYGFKGAEIAMAAYTSELQYLANPVTTHVQSAEQHNQ  505
PmePAL2      ELVCDYYNNGLPSNLSGGPNPSLDYGFKGAEIAMAAYTSELQYLANPVTTHVQSAEQHNQ  540
PsyPAL       ELVCDYYNNGLPSNLSGGPNPSLDYGFKGAEIAMAAYTSELQYLANPVTTHVQSAEQHNQ  390
PsiPTAL      ELVNDYYNGGLPSNLSGGPNPSLDYGLKGAEIAMASYTSELLYLANPVTSHVQSAEQHNQ  493
PabPTAL1     ELVNDYYNGGLPSNLSGGPNPSLDYGLKGAEIAMASYTSELLYLANPVTSHVQSAEQHNQ  521
PmePTAL1     ELVNDYYNGGLPSNLSGGPNPSLDYGLKGAEIAMASYTSELLYLANPVTSHVQSAEQHNQ  526
PsyPTAL      ELVNDYYNGGLPSNLSGGPNPSLDYGLKGAEIAMASYTSELLYLANPVTSHVQSAEQHNQ  491
PtaPTAL1     ELVNDYYNGGLPSNLSGGPNPSLDYGLKGAEIAMASYTSELLYLANPVTSHVQSAEQHNQ  491
PmePTAL2     ELVNDYYNGGLPSNLSGGPNPSLDYGFKGAEIAMAAYTSELLYLANPVTTHVQSAEQHNQ  497
PabPTAL2     ELVNDYYNGGLPSNLSGGPNPSLDYGFKGAEIAMAAYTSELQYLASPVTTHVQSAEQHNQ  491
             **: ::*. ** .**: . : *****:** ::***:* **: :*..*:*.**:*::****
```

```
PtaPTAL3     DINSLALISARKTEEALDILKLMLASHLYALCQAIDLRQLEQILLNIVLGIISSVSDECH  532
PmePTAL4     DINSLALISARKTEEALDILKLMVASHLSALCQAIDLRQLEQMLVKTVLGVISSVSDACH  532
PtaPTAL2     DVNSLGLISARKTAEAVDILKLMVSTYLIALCQAVDLRHLEENFHGAVKQIVCQAVRRIL  553
PmePTAL3     DVNSLGLISARKTAEAVEILELMVSTYLIALCQAVDLRHLEENFHGGVKQIVCQAAGRVL  545
PabPTAL3     DVNSLGLMSARKTAEAVDILKLMVSTYLIALCQAVDLRHLEENFHSGVKQIVCQAVRTIL  525
PtaPAL4      DVNSLGLISARMTAQAVEILKLMTSTYLVALCQAIDLRHLEENLHAAVRQAVGEACKKTL  540
PtaPAL3      DVNSLGLISARMTAQAVEILKLMTSTYLVALCQAIDLRHLEENLQGTVRQAVGQTFKNTL  547
PtaPAL1      DVNSLGLISARMTAQAVEILKLMTSTYLVALCQAIDLRHLEENLQGAVRQAVGQTFKKTL  547
PtaPAL2      DVNSLGLISARMTAQAVEILKLMTSTYLVALCQAIDLRHLEENLQGTVRQAVGQTFKKTL  547
PabPAL1      DVNSLGLISARMTAQAVEILKLMTSTYLVALCQAIDLRHLEENLQTAVRQAVAQACKKTL  547
PmePAL1      DVNSLGLISARMTAQAVEILKLMSSTYLVALCQAIDLRHLEENLQTAVRQAVGQACKKTL  548
OsPTAL       DVNSLGLVSARKTLEAVDILKLMTSTYIVALCQAVDLRHLEENIKSSVKNCVTQVAKKVL  535
ZmPTAL       DVNSLGLVSARKTAEAIDILKLMSSTYIVALCQAVDLRHLEENIKASVKNTVTQVAKKVL  535
BdPTAL       DVNSLGLVSARKTAEAVDILKLMSSTYMVALCQAVDLRHLEENIKASVKNCVTQVSKKVL  535
BoPTAL       DVNSLGLVSARKTAEAVDILKLMSSTYMVALCQAVDLRHLEENIKSSVKNCVTQVAKKVL  535
BoPAL        DVNSLGLISSRKTAEAIDILKIMSSTFLVALCQAIDLRHIEENVKSAVKSCVMTVAKKTL  545
OsPAL        DVNSLGLISSRKTAEAIDILKLMSSTFLIALCQAVDLRHIEENVKSAVKSCVMTVAKKTL  545
BdPAL        DVNSLGLISSRKTAEAIDILKIMSSTFLIALCQAIDLRHLEENMKTAVRNCVMQVAKKTL  549
ZmPAL        DVNSLGLISSRKTAEAIDILKLMSSTFLIALCQAIDLRHLEENVKAAVKNCVTQVAKKSL  550
PcPAL        DVNSLGLISSRKTSEAVEILKLMSTTFLVGLCQAIDLRHLEENLKSTVKNTVSSVAKRVL  548
AtPAL        DVNSLGLISSRKTSEAVDILKLMSTTFLVAICQAVDLRHLEENLRQTVKNTVSQVAKKVL  557
PtPAL        DVNSLGLISSRKTAESVDILKLMSTTFLVALCQAIDLRHLEENLRSAVKNTVSHVSKRVL  546
PabPAL2      DVNSLGLISARKTAEAVEILKLMFATYLVGLCQAVDLRHLEENMRSVVKHIVLQAARKTL  562
PsiPAL       DVNSLGLISARKTAEAVEILKLMFATYLVGLCQAVDLRHLEENMRSVVKHIVLQAARKTL  565
PmePAL2      DVNSLGLISARKTAEAIEILKLMFSTYMVGLCQAIDLRHLEENMRSVVKHIVLQAARKTL  600
PsyPAL       DVNSLGLISARKTAEAVEILKLMFATYLVALCQAIDLRHLEENMRSVVKHVVLQAARKTL  450
PsiPTAL      DVNSLGLISARKSAEAVDILKLMLSTYLTALCQAVDLRHLEENMLATVKQIVSQVAKKTL  553
PabPTAL1     DVNSLGLISARKSAEAVDILKLMLSTYLTALCQAVDLRHLEENMLATVKQIVSQVAKKTL  581
PmePTAL1     DVNSLGLVSARKSAEAIDILKLMLSTYLTALCQAVDLRHLEENMLATVKQIVSQVAKKTL  586
PsyPTAL      DVNSLGLVSARKSAEAIDILKLMLSTYLTALCQAVDLRHLEENMLATVKQIVSQVAKKTL  551
PtaPTAL1     DVNSLGLVSARKSAEAIDILKLMLSTYLTALCQAVDLRHLEENMLATVKQIVSQVAKKTL  551
PmePTAL2     DVNSLGLVSARKSAEAIDILNLMVSTYLLAICQAADLRHLEENMLSTVKRVVSQVAKKTL  557
PabPTAL2     DVNSLGLISARKSAEAIDILNMMVSTFLLALCQAADLRHLEENMLSTVKRVVSQVAKKTL  551
             *:***.*:*:* : ::::**::* ::.: .:*** ***::*: .   *   :  .

PtaPTAL3     -----------LQQSIKEQLISVASGIPVYTYLESPCNPSLPLVSALKQTFLDAIVTSH-  580
PmePTAL4     -----------LPESIKVQLVNVARGIPIYTYLESPCDPSLPLLSAIKQTFLDSILTFH-  580
PtaPTAL2     YSTTEQGILVLPFGFYENKLLQVVDCLPVFSYMENPTSASSPLTLQLRHVLVEQALKADT  613
PmePTAL3     YSTTDQGILLSPFRFCEKELLQVVDRLPVFSYIEDPTGPSSPLMLQLRQVLVEQALKGNT  605
PabPTAL3     YSTTDQGISLLPFRFCENELMQVVDRLPVFSYVEDPAGPSSPLMLQLRHVLVELALKANS  585
PtaPAL4      VVGPRG--ELL--------LLKAVDREPVFSYIDNPFSATSVLTTVLRQVLFEHALEKTT  590
PtaPAL3      VVGSRG--ELLNSRFCEKDLLKVVDLEAVFSYIDNPCSATSVLTTKLRQVLLEHALENKT  605
PtaPAL1      VVGSRG--ELLNSRFCEKDLLRVVDREAVYSYIDNPCSATSVLTTKLRQVLLEHALESKT  605
PtaPAL2      VVGSRG--ELLNSRFCEKDLLKVVDLEAVFSYIDNPCSATSVLTTKLRQVLLEHALENKT  605
PabPAL1      VVGPQG--ELLDSRFCEKDLLKAVEREPVFSYIDNPCSATSVLTTKLRQVLFEHAVEKTT  605
PmePAL1      AVGPQG--ELLPSRFCEKDLLMAVERETVFSYIDNPCSATSVLTTKLRQVLFEHALEKTT  606
OsPTAL       TMNPTG--DLSSARFSEKNLLTAIDREAVFSYADDPCSANYPLMQKLRAVLVEHALTSGD  593
ZmPTAL       TMNPSG--ELSSARFSEKELISAIDREAVFTYAEDAASGSLPLMQKLRAVLVDHALSSGD  593
BdPTAL       TMNPTG--DLSSARFSEKSLLTAIDREAVFSYADDACSANYPLMQKLRAVLVDHALTSSG  593
BoPTAL       TMNPTG--DLSSARFSEKNLLTAIDREAVFTYADDPCSANYPLMQKLRAVLVDHALTSGD  593
BoPAL        STNSTG--DLHVARFCEKDLLKEIDREAVFAYADDPCSPNYPLMKKMRNVLVERALANGM  603
OsPAL        STNSTG--DLHVARFCEKDLLKEIDREAVFAYADDPCSHNYPLMKKLRNVLVERALANGA  603
BdPAL        SMNAMG--GLHIARFCEKDLLTAIDREAVFAYADDPCSPNYPLMQKLRAVLIEHALANGD  607
ZmPAL        SLNARG--GLHNARFCEKDLQTAIDREAVFAYADDPCSPNYALMQKLRAVLVEHALANGD  608
PcPAL        TMGVNG--ELHPSRFCEKDLLRVVDREYIFAYIDDPCSATYPLMQKLRQTLVEHALKNGD  606
AtPAL        TTGVNG--ELHPSRFCEKDLLKVVDREQVYTYADDPCSATYPLIQKLRQVIVDHALINGE  615
PtPAL        TTGANG--ELHPSRFCEKELLKVVDREDVFAYADDPCSATYPLMQKLRQVLVDHALANGE  604
PabPAL2      YTAEDG--SLLDTRFCEKELLQVIDHQPVFSYIDDPTNPSYALMLQLREVLVDESLRLSC  620
PsiPAL       YTAEDG--SLLDTRFCEKELLQVIDHQPVFSYIDDPTNPSYALMLQLREVLVDESLRLSC  623
PmePAL2      YTAEDG--SLLDTRFCEKELLQVIDHQPVFSYIDDPTNPSYALMLQLREVLVDEALKLNC  658
PsyPAL       CTAEDG--SLQDTGFCEKELLQVIDHQPVFSYIDDPTNPSYALMLQLREVLVDEALKSSC  508
PsiPTAL      STGLNG--ELLPGRFCEKDLLQIVDNEHVFSYIDDPCNASYPLTQKLRNVLVEHAFKNTD  611
PabPTAL1     STGLNG--ELLPGRFCEKDLLQIVDNEHVFSYIDDPCNASYPLTQKLRNVLVEHAFKNTD  639
PmePTAL1     STGLNG--ELLPGRFCEKDLLQIVDNEHVFAYIDDPCNASYPLTQKLRNVLVEHAFKNTE  644
PsyPTAL      STGLNG--ELLPGRFCEKDLLQVVDNEHVFSYIDDPCNASYPLTQKLRNILVEHAFKNGE  609
PtaPTAL1     STGLNG--ELLPGRFCEKDLLQVVDNEHVFSYIDDPCNASYPLTQKLRNILVERAFKNAE  609
PmePTAL2     STHNGE--LLTAGRFCEKDLLQAVENLHVFAYVDDPCNENYPLMQQLRQVLVAHALNET-  614
PabPTAL2     STHNGE--LLTAGRFCEKDLLQAVENMHVFAYIDDPCNENYPLMQQLRQVLVAHALSES-  608
                                *        :::* :.  . .  *   ::  :.   .
```

```
PtaPTAL3      -----------------------DIQIVEQITEFECHLKQRLEEEITAIVLSYEERTN-S   616
PmePTAL4      -----------------------DIQIVEKIKEFESHLKQRLEEEITATRLSYEQRTN-I   616
PtaPTAL2      ----------------------EQYSLLSKIPIFEEEVRRKLAIEVPLLRQRY-ERGD-F   649
PmePTAL3      ----------------------EQYSLLGKISMFEDELRRILAIEVARIRQRC-ERGD-F   641
PabPTAL3      ----------------------DQYSLLGKISMFEEELRRKLAIEVPLIRQKC-ERGD-F   621
PtaPAL4       ---------------------DNDGSILTRVPAFEEELKARIVADVHETRAAC-EKGT-A   627
PtaPAL3       ---------------------DNDASILTRIPAFEKELKAQIVAGVQETRAAC-EKDT-A   642
PtaPAL1       ---------------------DNDASILARIPEFEEELKAQIVAEVQERRAAC-EKGT-A   642
PtaPAL2       ---------------------DNDASILTRIPAFEEELKAQIVAGVQERRAAC-EKGT-A   642
PabPAL1       ---------------------DNDASILTRIPAFEEELKARMVAEVQETREAF-EKRT-A   642
PmePAL1       ---------------------DNDASLLTKIPAFEAELKARIVAEVHEKRDAY-EKGA-A   643
OsPTAL        RRA----------------RGLRVLQDH---QVRGGAPLCAAPGDR-GRPRRR-RQRT-A   631
ZmPTAL        --------------------AEREPSVFSKITRFEEELRAVLPQEVEAARVAV-AEGT-A   631
BdPTAL        VDN----------------AGESEATVFSKINKFEEELRAALPREIEAARVAF-EKGT-A   635
BoPTAL        --------------------AEREPSVFSKITKFEEELRSALPREIEAARVAV-ADGT-A   631
BoPAL         AEF------------------NAETSVFAKVAQFEEELRATLPRAVEAARAAV-ENGT-A   643
OsPAL         AEF------------------NADTSVFAKVAQFEEELRATLPGAIEAARAAV-ENGT-A   643
BdPAL         GER------------------ALETSIFAKVAEFEQNLRAALPKEVEAARASV-ENGT-P   647
ZmPAL         AER------------------DVDTSIFAKVAEFEQQVRAALPKEVEAARAAV-ENGS-P   648
PcPAL         NER------------------NLSTSIFQKIATFEDELKALLPKEVESARAAL-ESGN-P   646
AtPAL         SEK------------------NAVTSIFHKIGAFEEELKAVLPKEVEAARAAY-DNGT-S   655
PtPAL         NEK------------------NTSTSVFQKITAFEEELKALLPKEVESARAAY-DSGN-S   644
PabPAL2       LGKDGESDHNSEAAD---ASGVLSNWVFGRIPLFQQELKARLEDEVPKARERF-DKGD-F   675
PsiPAL        LGKDGESDHNLEAAD---ASGVLSNWVFGRIPLFQQELKARLEDEVPKARERF-DKGD-F   678
PmePAL2       TVKDAESGHKLEAGDGAGAAGTIPNWVFCKIPIFQQELKARLEEEVPKARERF-DRGD-F   716
PsyPAL        PEGNAESDRNLQAAESAGAAGILPNWVFSRIPIFQEELKARLEEEVPKARERF-DNGD-F   566
PsiPTAL       GEK------------------DPNTSIFNKITLFEAELKTQLELQVNLARESY-DKGI-S   651
PabPTAL1      GEK------------------DPNTSIFNKITLFEAELKTQLELQVNLARESY-DKGI-S   679
PmePTAL1      GEK------------------DPNTSIFNKITLFEAELKTQLESQVNLARDSY-DKGI-S   684
PsyPTAL       GEK------------------DPNTSIFNKIPLFEAELKAQLELQVSLARESY-DKGT-S   649
PtaPTAL1      GEK------------------DPNTSIFNKIPVFEAELKAQLEPQVSLARESY-DKGT-S   649
PmePTAL2      ---------------------ESQSSIFDKIPVFEKELKEQMEAEIGRARNDYYEKGIAG   653
PabPTAL2      ---------------------EIQSSIFNKIPVFEKELKDQMEAEIGRARNDYYEKGIAG   647


PtaPTAL3      H-MLEGSCCRTLIHGSKFFPLYAFIREELNAIMMTPRTDHTPQEGT----------QKVF   665
PmePTAL4      HMMLEGSCCRTLIHGSKFFPLYAFIREELNAKLMTPRTDNTPQEDI----------QKVF   666
PtaPTAL2      D-------LPNKIRECRTYPLYKFVREELGTSLLSGPRGRTPGEDI----------DKVF   692
PmePTAL3      D-------FPNKIRECRTYPLYDFVREELGTSLLSGPRGRTPGEDI----------DKVF   684
PabPTAL3      D-------LPNKVRECRTYPLYEFVREELGTSLLSGPRGRTPGEDI----------DRVF   664
PtaPAL4       L-------VPNRIKDCRSYPLYEFVRAELGTSLLVGTDSRSPGEDF----------DKVF   670
PtaPAL3       L-------VPNRIKDCCSYPLYEFVRVELGTSLLVGTNSRSPGEDFDKVRSSGEDFEKVF   695
PtaPAL1       P-------VPNRIKDCRSYPLYEFVRVELGTSLLVGTNSHSPGEDF----------DKVF   685
PtaPAL2       L-------VPNRIKDCRSYPLYEFVRVELGTSLLVGTNSRSPGEDF----------DKVF   685
PabPAL1       L-------VPNRIKDCRSYPLYEFVRLELGTSLLVGTNSHSPGEDF----------DKVF   685
PmePAL1       F-------VPNRIKDCRSYPLYEFVRLELGTSLLVGTNSHSPGEDF----------DKVF   686
OsPTAL        P-------VANRIVESRSFPLYRFVREELGCVFLTGEKLKSPGEEC----------NKVF   674
ZmPTAL        P-------VANRIADSRSFPLYRFVREELGCVFLTGERLKSPGEEC----------NKVF   674
BdPTAL        P-------IPNLIKDSRSFPLYRFVREELGCVYLTGEKLLSPGEEC----------NKVF   678
BoPTAL        P-------IANRIKESRSFPVYRFVREELGCVYLTGEKLKSPGEEC----------NKVF   674
BoPAL         A-------TPNRITECRSYPLYRFVREELGTAYLTGEKTRSPGEEL----------NKVL   686
OsPAL         A-------IPSRITECRSYPLYRFVREELGTKYLTGEKTRSPGEEL----------NKVL   686
BdPAL         L-------APNRIKDCRSYPLYRFVREELGTEYLTGEKTRSPGEEL----------NKVL   690
ZmPAL         L-------VPNRIKECRSYPLYRFVREEVGTKYLTGEKTRSPGEEL----------NKVL   691
PcPAL         A-------IPNRIEECRSYPLYKFVRKELGTEYLTGEKVTSPGEEF----------EKVF   689
AtPAL         A-------IPNRIKECRSYPLYRFVREELGTELLTGEKVTSPGEEF----------DKVF   698
PtPAL         A-------IENKIKECRSYPLYKFVREELGTGLLTGEKVRSPGEEF----------DKVF   687
PabPAL2       P-------IANRINKCRTYPIYRFVRSELGTDLLTGPKWRSPGEDI----------EKVF   718
PsiPAL        P-------IANRINKCRTYPIYRFVRSELGTDLLTGPKWRSPGEDI----------EKVF   721
PmePAL2       P-------IANRINKCRTYPIYRFVRSELGTDLLTGPKWRSPGEDI----------EKVY   759
PsyPAL        P-------IANRINKCRTYPIYRFVRSELGTDLLTGPKWRSPGEDI----------EKVF   609
PsiPTAL       P-------LPNRIQECRSYPLYEFVRTQLGTKLLSGTRTTSPGEVI----------ELVY   694
PabPTAL1      P-------LPNRIQECRSYPLYEFVRTQLGTKLLSGTRTISPGEVI----------ELVY   722
PmePTAL1      P-------LPNRIQECRSYHLYEFVRNQLGTKLLSGARTTSPGEVI----------ELVY   727
PsyPTAL       P-------LPNRIQECRSYPLYEFVRKQLGTKLLSGTRTISPGEVI----------EVVY   692
PtaPTAL1      P-------LPNRIQECRSYPLYEFVRNQLGTKLLSGTRTISPGEVI----------EVVY   692
PmePTAL2      S-------IPNRIQDCRSFPLYDFARSQLGTQLLSGDRTTSPGEYI----------GKVY   696
PabPTAL2      S-------VSNRIQECRSFPLYDFVRSQLGTQLLSGDRVTSPGEYI----------EKVY   690
                        . :  .  : :* * * ::.   :      :* *              *
```

```
PtaPTAL3      DAIVDGRITAPLLQCLNGFMN---------------------------------------  686
PmePTAL4      DAIVDGRITVPLLLCLNGFMK---------------------------------------  687
PtaPTAL2      IAITEGKLEGRLMECLDGWNESPGPFNDLKKNANHYDILKSHNNNSCVWSWFQQIGGPQV  752
PmePTAL3      IAITEGKLEGRLMECLDGWNESPGPFNDLKKNANNDDVVRTHSKNSCVWSWFQHIGGPPV  744
PabPTAL3      IAITEGKLEGRLMECLDGWNESPGPFNGMKKNANNYDIVKTPNNNSCVWSWFQKMGGPQV  724
PtaPAL4       VAINEGKAVAPLFKCLEGWNGAPIPI----------------------------------  696
PtaPAL3       VAINEGKAVEPLFKCLEEWNGAPIPILNLNITN------RNPKESECCRS----------  739
PtaPAL1       VAINEGKTVEPLFKCLEKWNGAPIPI----------------------------------  711
PtaPAL2       VAINEGKAVEPLFKCLEKWNGVPIPI----------------------------------  711
PabPAL1       VAINEGKAVEPLFKCLERWNGAPIPI----------------------------------  711
PmePAL1       VAINEGKAVEPLLKCLERWNGAPIPI----------------------------------  712
OsPTAL        LGISQGKLIDPMLDCLKEWNGEPLPIN---------------------------------  701
ZmPTAL        VGISQGKLVDPMLECLKEWDGKPLPINVK-------------------------------  703
BdPTAL        IGISQGKLIDPMLECLKEWNGEPLPINVV-------------------------------  707
BoPTAL        IGISQGKLIDPMLECLKEWNGEPLPIN---------------------------------  701
BoPAL         LAINQGKHIDPLLECLKEWNGEPLPIN---------------------------------  713
OsPAL         VAINEGKHIDPLLECLKEWNGEPLPIC---------------------------------  713
BdPAL         VAMNQRKHIDPLLECLKEWNGEPLPLC---------------------------------  717
ZmPAL         VAINQRKHIDPLLECLKEWNGEPLPLC---------------------------------  718
PcPAL         IAMSKGEIIDPLLECLESWNGAPLPIC---------------------------------  716
AtPAL         TAICEGKIIDPMMECLNEWNGAPIPIC---------------------------------  725
PtPAL         TAMCQGKIIDPMLECLGEWNGAPLPIC---------------------------------  714
PabPAL2       EGICEGKMGEVILKCLDAWRGCAGPFTPRAYPA------SPAAFNTSYWAWFDNTKSPSA  772
PsiPAL        EGICEGKMGEVILKCLDAWRGCAGPFTPRAYPA------SPAAFNTSYWAWFDNTKSPSA  775
PmePAL2       EGICDGKIGDVILKCLDAWRGCAGPFTPRAYPA------SPAAFNTSYWAWFDSTKSPSA  813
PsyPAL        EGICQGKIGDVILKCLDAWGGCAGPFTPRAYPA------SPAAFNASYWAWFDSTKSPSA  663
PsiPTAL       DAISEDKIIGPLLKCVEGWKATPGPF----------------------------------  720
PabPTAL1      DAISEDKIIGPLLKCVEGWKATPGPF----------------------------------  748
PmePTAL1      DAISEDKIIGPLFKCLDGWKATPGPF----------------------------------  753
PsyPTAL       DAISEDKVIVPLFKCLDGWKGTPGPF----------------------------------  718
PtaPTAL1      DAISEDKVIVPLFKCLDGWKGTPGPF----------------------------------  718
PmePTAL2      DGIREGKIIAPLLKCLDGWSGTPGPFPS--------------------------------  724
PabPTAL2      VGICEGKIISPLFKCLDGWSGTPGPFQS--------------------------------  718
               .: . .    :: *:  :

PtaPTAL3      ---------------     686
PmePTAL4      ---------------     687
PtaPTAL2      NGGKGYWLLTIA---     764
PmePTAL3      NGIKGYWLLSIA---     756
PabPTAL3      NGGKGYWLLSIA---     736
PtaPAL4       ---------------     696
PtaPAL3       ---------------     739
PtaPAL1       ---------------     711
PtaPAL2       ---------------     711
PabPAL1       ---------------     711
PmePAL1       ---------------     712
OsPTAL        ---------------     701
ZmPTAL        ---------------     703
BdPTAL        ---------------     707
BoPTAL        ---------------     701
BoPAL         ---------------     713
OsPAL         ---------------     713
BdPAL         ---------------     717
ZmPAL         ---------------     718
PcPAL         ---------------     716
AtPAL         ---------------     725
PtPAL         ---------------     714
PabPAL2       TSGRGFWSAQQQQVL     787
PsiPAL        TSGRGFWSAQQQQVL     790
PmePAL2       TSGRGFWSAQQQQIL     828
PsyPAL        TSGRGFWSAQQQQVL     678
PsiPTAL       ---------------     720
PabPTAL1      ---------------     748
PmePTAL1      ---------------     753
PsyPTAL       ---------------     718
PtaPTAL1      ---------------     718
PmePTAL2      ---------------     724
PabPTAL2      ---------------     718
```

**Figure S18** Latitudinal variation in the SNP (Ala228Ala) from PabPAL2 gene in Norway spruce populations across Sweden. (a) Cline in the allele frequencies of Ala228Ala. (b) Cline in the genotype frequencies of Ala228Ala. One-way ANOVA and Tukey's posthoc test was performed with the genotype frequencies. Tukey's posthoc categorization is indicated above the bars.

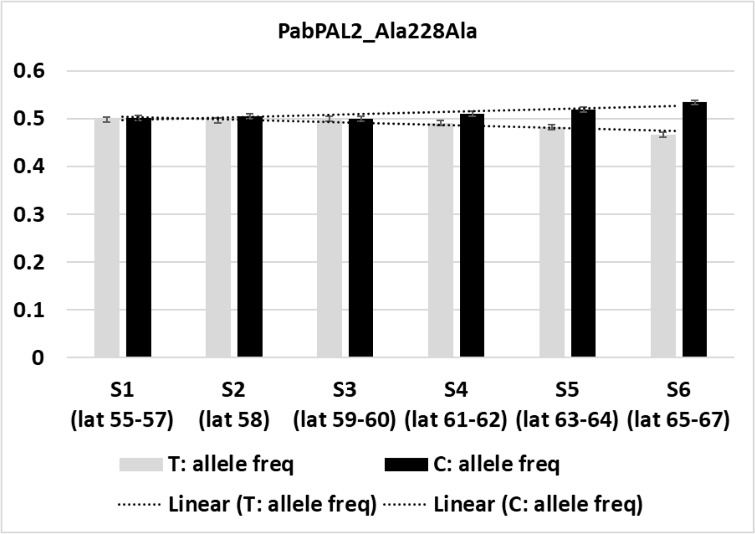


(a)

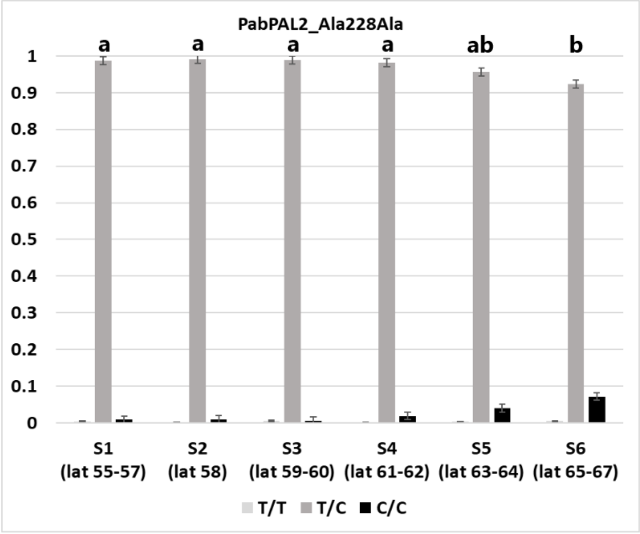


(b)

**Figure S19** Latitudinal variation in the SNP (Pro356Pro) from PabPAL2 gene in Norway spruce populations across Sweden. (a) Cline in the allele frequencies of Pro356Pro. (b) Cline in the genotype frequencies of Pro356Pro. One-way ANOVA and Tukey's posthoc test was performed with the genotype frequencies. Tukey's posthoc categorization is indicated above the bars.

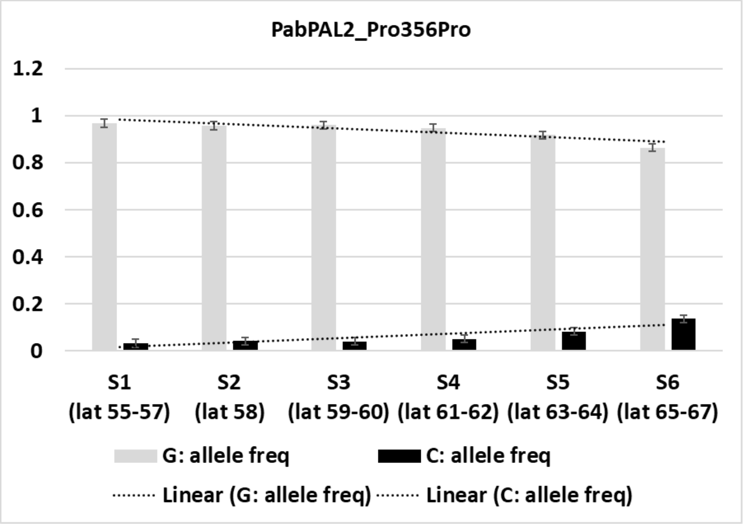


(a)

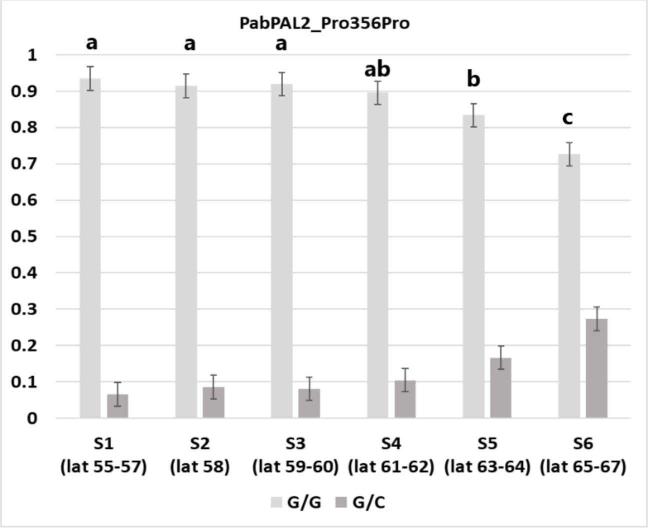


(b)

**Figure S20** Latitudinal variation in the SNP (Ala383Thr) from PabPAL2 gene in Norway spruce populations across Sweden. (a) Cline in the allele frequencies of Ala383Thr. (b) Cline in the genotype frequencies of Ala383Thr. One-way ANOVA and Tukey's posthoc test was performed with the genotype frequencies. Tukey's posthoc categorization is indicated above the bars.

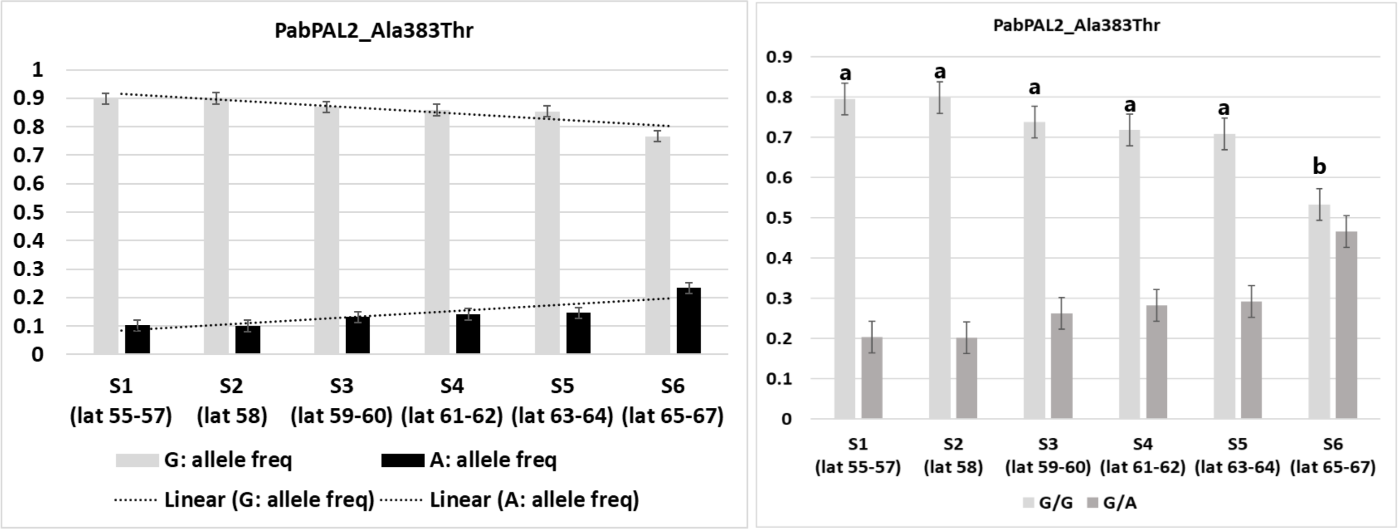


(a) (b)

**Figure S21** Latitudinal variation in the SNP (Ala535Val) from PabPTAL1 gene in Norway spruce populations across Sweden. (a) Cline in the allele frequencies of Ala535Val. (b) Cline in the genotype frequencies of Ala535Val. One-way ANOVA and Tukey's posthoc test was performed with the genotype frequencies. Tukey's posthoc categorization is indicated above the bars.

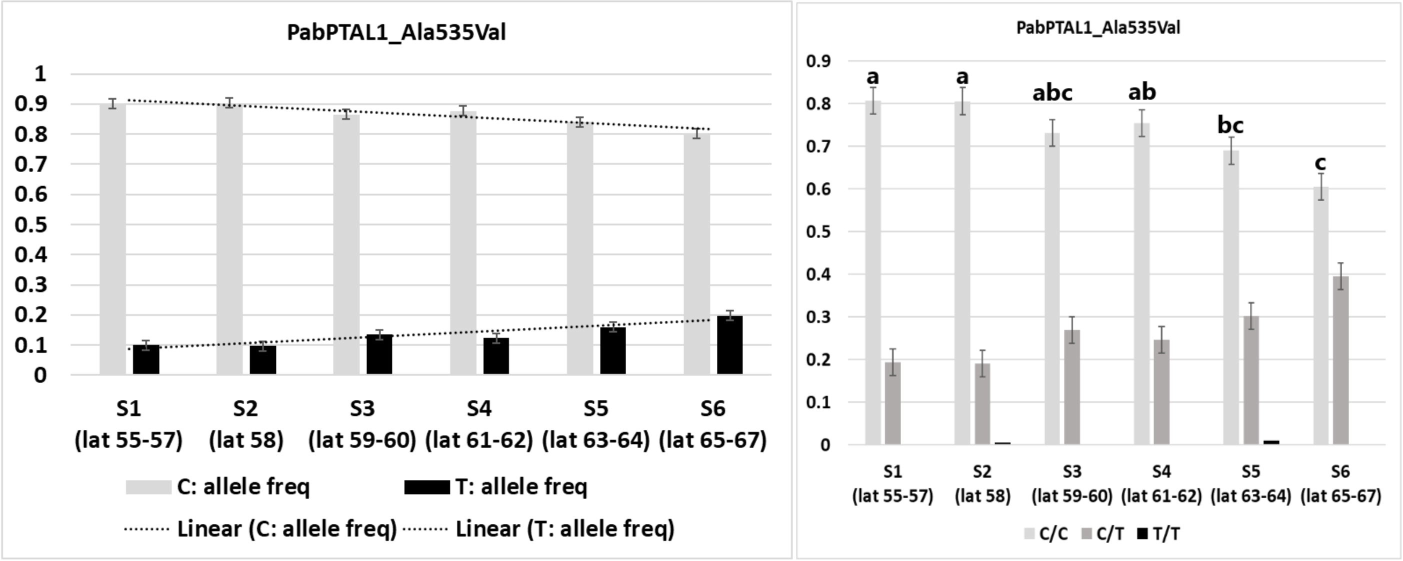


(a) (b)

**Figure S22** Latitudinal variation in the SNP (Ser595Phe) from PabPTAL3 gene in Norway spruce populations across Sweden. (a) Cline in the allele frequencies of Ser595Phe. (b) Cline in the genotype frequencies of Ser595Phe. One-way ANOVA and Tukey's posthoc test was performed with the genotype frequencies. Tukey's posthoc categorization is indicated above the bars.

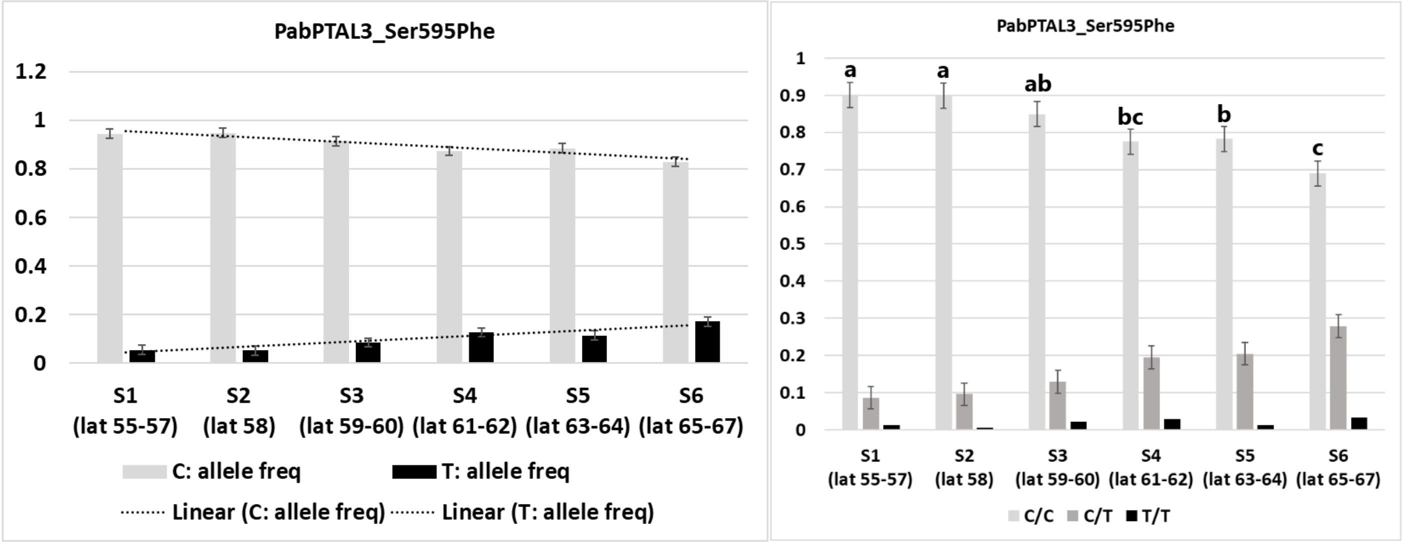


(a) (b)

**Figure S23** Latitudinal variation in the SNP (Leu601Phe) from PabPTAL3 gene in Norway spruce populations across Sweden. (a) Cline in the allele frequencies of Leu601Phe. (b) Cline in the genotype frequencies of Leu601Phe. One-way ANOVA and Tukey's posthoc test was performed with the genotype frequencies. Tukey's posthoc categorization is indicated above the bars.

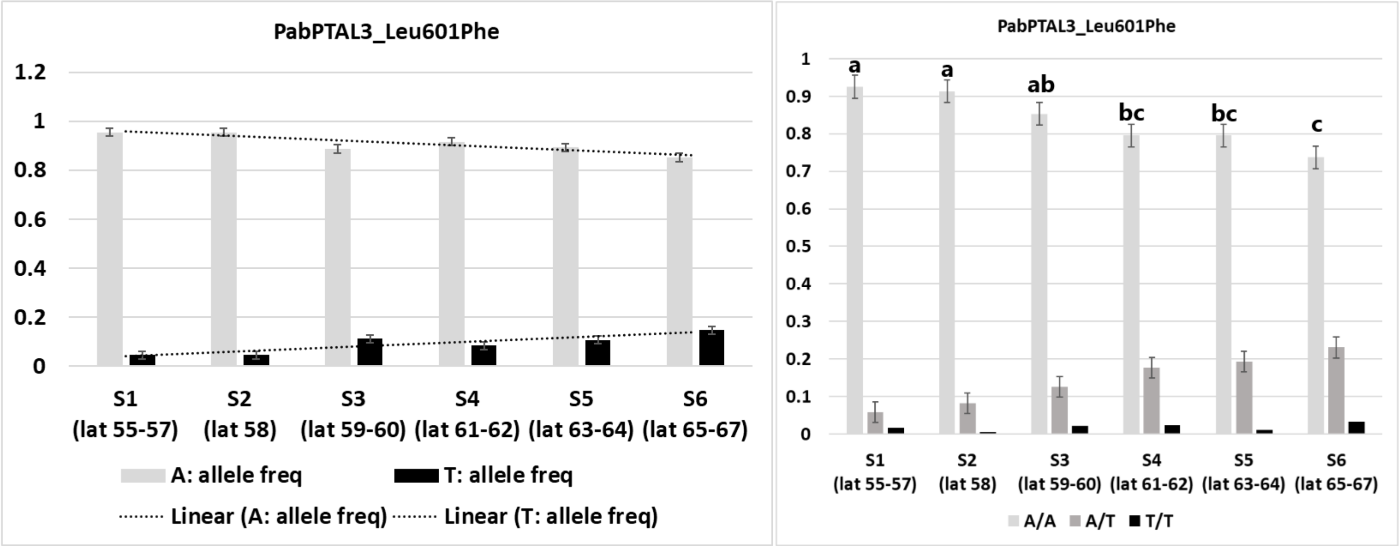


(a) (b)
